# scOnion: Cross-Modal Concordance-Aware Joint Analysis of Cells and Features in Single-Cell Multiomic Data

**DOI:** 10.64898/2026.11.24.769043

**Authors:** Ziqi Fu, Junhao Zhu, Nicholas A Pease, Jishnu Das, Harinder Singh, Xihong Lin, Rong Ma

## Abstract

Single-cell multiomic assays jointly profile gene expression and chromatin accessibility in the same cells, enabling direct analysis of cross-layer coordination. However, agreement between modalities varies across cells, and existing integration methods often emphasize shared structure, potentially averaging away cell-specific cross-modal discordance that carries biological signal. Here we present scOnion, a concordance-aware computational framework that quantifies cross-modality agreement and propagates this information into downstream analyses. Starting from paired low-rank representations obtained by spectral integration, scOnion computes per-cell concordance scores, a concordance-aware joint cell embedding, and an induced cross-modal feature embedding. The cell embedding preserves structure strongly represented in either modality, improving cell-type and state resolution, while the feature embedding scores candidate regulatory element(rE)-to-gene (rE2G) pairs. Across immune, neural, and developmental functional genomic datasets, scOnion’s concordance patterns are reproducible and cell-resolved, with low-concordance cells enriched for modality-specific structure. scOnion also prioritizes candidate rE2G pairs, including distal links, with stronger support from external Hi-C contacts and fine-mapped eQTL data than the baseline linkage methods. In a B-cell fate bifurcation study, scOnion reveals remodeling of concordance and rE2G coupling during lineage commitment. scOnion embeds 70,000 cells and computes rE2G links on CPU in 30 minutes, extends naturally to other multiomic assays such as CITE-seq, and is available as open-source software.

## 1 Introduction

Single-cell multiomic technologies capture multiple molecular layers within the same cells [1–4], providing a direct view of how cellular programs are coordinated across modalities. This coordination is often partial, cell-state-dependent, and dynamically changing, yet the degree to which the modalities agree is often absorbed into integrated representations without direct quantification. Such partial decoupling may reflect biologically meaningful differences in how cellular programs are encoded, primed, delayed, or resolved across molecular layers. Characterizing this cell-resolved het-erogeneity, and determining when it should be preserved in downstream analysis, remains an open challenge. This gap has implications at two linked levels: at the cell level, concordance can reveal heterogeneity in how cellular states are manifested across modalities, while at the feature level, it can illuminate how genes and regulatory elements participate in coordinated or modality-specific programs.

Current analysis of cell-paired single-cell multiomic data is typically organized around two computational scaffolds: cell embeddings and feature linkage [5]. Cell-centric methods can be viewed as emphasizing either the intersection or the union of modality-specific information. Intersection-type methods, such as the common representations produced by Tilted-CCA [6] and MOJITOO [7], recover shared structure but may suppress modality-specific biological variation in the joint embedding. Union-type methods [8], including MOFA2 [9] and MIRA [10], retain broader variation but usually do not quantify whether a local structure is supported by both modalities or mainly by one. However, cross-modality coupling is often graded rather than binary, and downstream analyses may benefit from knowing where modalities agree and diverge, and whether such divergence is biologically structured. Feature-linkage approaches address a complementary problem by relating molecular features across layers, but they are usually developed separately from cell embeddings. In scRNA+scATAC data, candidate regulatory elements (REs) are the accessible-chromatin peaks measured by ATAC, and rE2G linkage is commonly inferred using window-based correlation meth-ods, such as Signac [11] and FigR [12], which are restricted to predefined *cis* windows and obtain per-link uncertainty from permutation nulls. Graph-embedding methods, such as SIMBA [13] and scGLUE [14], are not restricted to predefined *cis* windows, but require trained models and can be harder to interpret. Thus, current methods lack a single geometry that makes cross-modal concordance an explicit, cell-resolved quantity and connects it to both the cell embedding and feature-level regulatory coupling.

To address these limitations, here we present scOnion, a concordance-aware computational framework for paired single-cell multiomic data. scOnion first uses a spectral objective to learn comparable low-dimensional representations across modalities, from which it defines per-cell concordance as a cell-specific measure of cross-modality agreement. These representations are then combined into a joint cell embedding that preserves broad cell-type organization shared across modalities while retaining finer state or subtype structure that may be more strongly supported by one modality than the other. The concordance scores indicate where such refinements are supported by both modalities or by modality-specific structure. At the feature level, scOnion in-duces a shared embedding of features across modalities, enabling broad candidate searches beyond predefined pairwise tests. Feature proximity in this embedding highlights candidate cross-modal relationships, while a per-link uncertainty estimate helps prioritize well-supported associations over high-magnitude but unstable links. scOnion therefore combines the broad candidate search of embedding-based approaches with the uncertainty-aware prioritization of correlation-based meth-ods for efficient linkage prioritization. Because the construction reduces to a sequence of low-rank matrix operations, scOnion produces a genome-wide cis linkage map for a 20K-cell multiome in 18 minutes on 16 CPU cores, faster than the other ten linkage methods evaluated. It further scales to 70K cells, producing both the joint cell embedding and genome-wide linkage map in 32 minutes without a GPU. Together, these properties make scOnion suitable for large-scale multiomic analyses in which the degree of cross-modal agreement varies across cells and is itself of biological interest.

Across ten paired scRNA+scATAC immune, neural, and developmental datasets comprising more than 72K cells, scOnion recovers reproducible, cell-resolved transcriptome–epigenome concor-dance patterns that are not readily accessible by existing methods, improves cell-type and cell-state resolution through concordance-guided joint embedding, and prioritizes cis rE2G links up to 1 Mb from the TSS, including distal links, with stronger support from independent chromatin-contact and eQTL evidence than window-based and embedding baselines across four immune cell types, and enrichment for lineage transcription-factor motifs. In a B-cell fate bifurcation study consisting of scRNA+scATAC profiles of 28K cells, scOnion further reveals remodeling of modality concordance and rE2G coupling during lineage commitment. Because the framework is not specific to a particular modality pair, it extends naturally to other multiomic assays, including CITE-seq, where it links surface proteins to transcriptional programs, and to broader integration settings in which cells are computationally paired across modalities.

## 2 Results

### 2.1 Overview of scOnion

scOnion takes paired high-dimensional single-cell multiomic profiles as input and returns three linked analysis objects: cell-wise concordance scores, a joint cell embedding, and a cross-modal feature map with reliability estimates for candidate feature links (Fig. 1a). The workflow first learns paired low-dimensional coordinates for the two modalities through spectral integration. This paired construction preserves strong within-modality structure while making coordinates directly comparable across modalities (Fig. 1b). Using these matched coordinates, scOnion defines per-cell concordance as a local measure of cross-modality agreement. Concordance indicates whether the structure around a cell is jointly supported by both modalities or driven primarily by one. This in-formation is not directly visible from embedding geometry alone. Similar separations in a union-like embedding may reflect strong agreement between modalities in one setting but uneven modality support in another (Fig. 1c). Low concordance can therefore flag either biologically meaningful asymmetry between molecular layers or residual technical effects that warrant follow-up. scOnion then combines the paired coordinates to construct a joint cell embedding that retains broad cell-type organization as well as finer state or subtype structure that may be stronger in one modality than the other. The embedding captures candidate biological structure, while the concordance scores indicate where that structure is strongly supported by both modalities or driven mostly by one (Fig. 1d). The same joint cell space also anchors feature-level analysis. Modality-specific features are projected into a shared feature embedding, where embedding proximity ranks candidate cross-modal couplings. Each candidate link additionally carries a reliability estimate, obtained efficiently in closed form, enabling uncertainty-adjusted ranking of links by stability as well as magnitude. Because the cell and feature analyses derive from the same paired low-dimensional co-ordinates, scOnion provides a unified workflow for concordance-aware cell representation to support interpretable feature-coupling analysis within any user-defined cell subpopulation.

**Figure 1:**
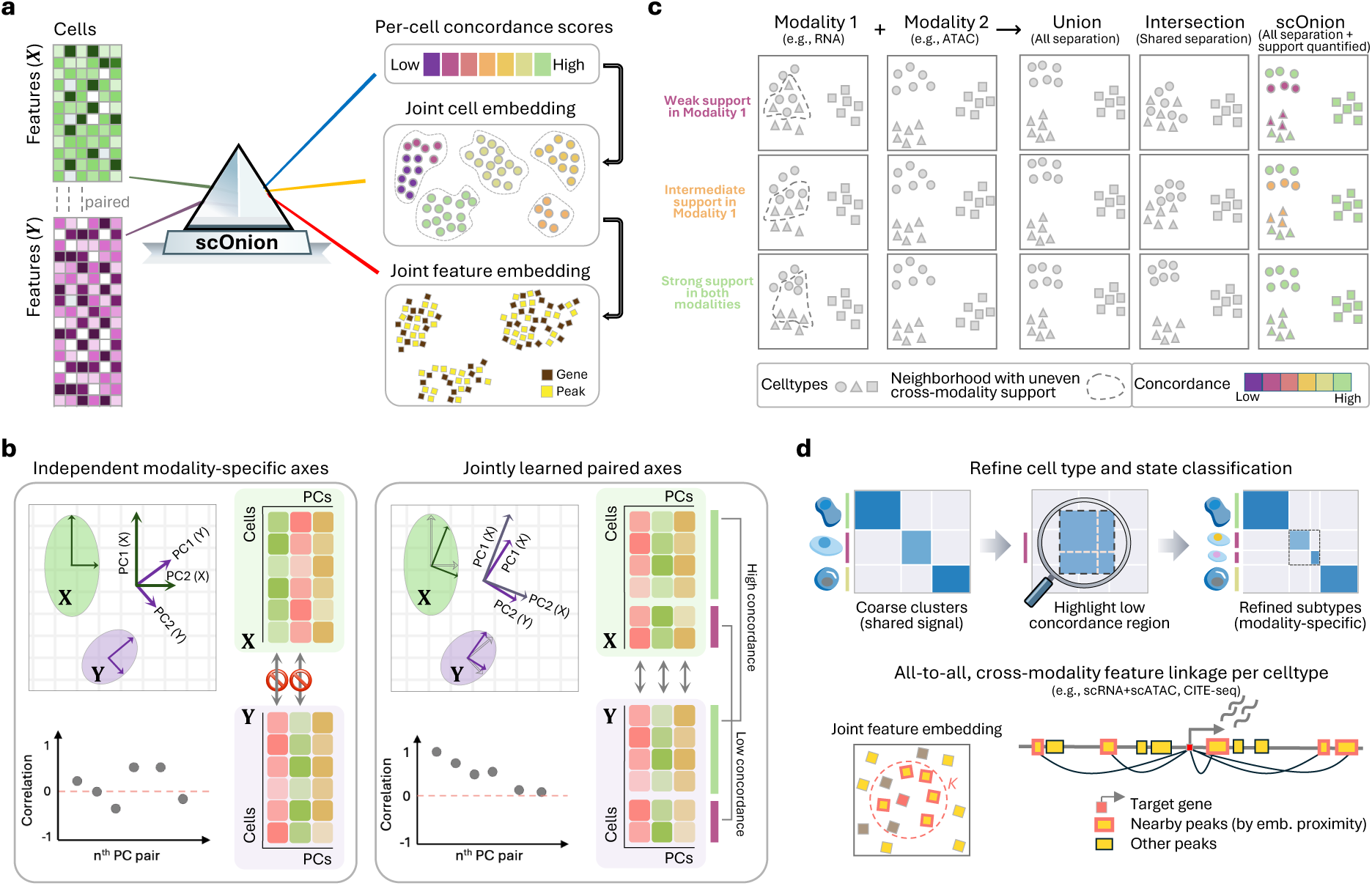
Schematic overview of scOnion. **a**, scOnion takes as input cell-paired measurements across modalities and jointly learns a cell embedding together with a per-cell concordance score that quantifies the agreement between a cell’s two modality representations. The joint cell embed-ding provides a shared coordinate system for cells and is used to project modality-specific feature matrices into a joint feature embedding. **c**, Comparison of union-based and intersection-based joint cell embeddings under different strengths of shared modality structure. scOnion retains separations that are strong in either modality, as the union does, and its per-cell concordance (color) additionally makes the degree of cross-modality support explicit, so a neighborhood separated in only one modality (dashed outline) is marked as low concordance. **b**, scOnion summarizes each modality’s cell-cell similarity structure with leading low-dimensional components. Unlike conventional PCA and other approaches that optimize each modality independently, scOnion captures dominant shared structure with paired leading axes while allowing strong but modality-specific structure to persist in less aligned directions. This pairing yields comparable coordinates across modalities and thereby enables per-cell concordance to quantify cell-level divergence between the two modality representations. **d**, The per-cell concordance score supports a pragmatic clustering workflow that prioritizes low-concordance neighborhoods in the joint cell embedding for modality-specific interrogation and refined subtyping. The joint feature embedding supports cross-modality feature linkage over broad candidate sets by embedding proximity. This is a schematic overview and contains no experimental data.

### 2.2 scOnion quantifies cell-resolved cross-modality concordance

scOnion starts from two cell-paired multiomic matrices that share the same cells but may contain different feature sets. After modality-appropriate normalization and denoising, scOnion learns paired low-dimensional coordinate matrices **U** and **V** for the two modalities through a spectral alignment objective (Methods). This objective captures the dominant variation within each modality while prioritizing latent directions that covary across modalities. The resulting alignment matches the dimensions of **U** and **V** one to one, making the two representations directly comparable. For each cell *i*, scOnion measures cross-modality agreement by the cosine similarity between its paired coordinate vectors, **U***_i_*_·_ and **V***_i_*_·_, and rescales this value to [0, 1] to define the cell’s concordance score (Fig. 2a and Methods).

**Figure 2:**
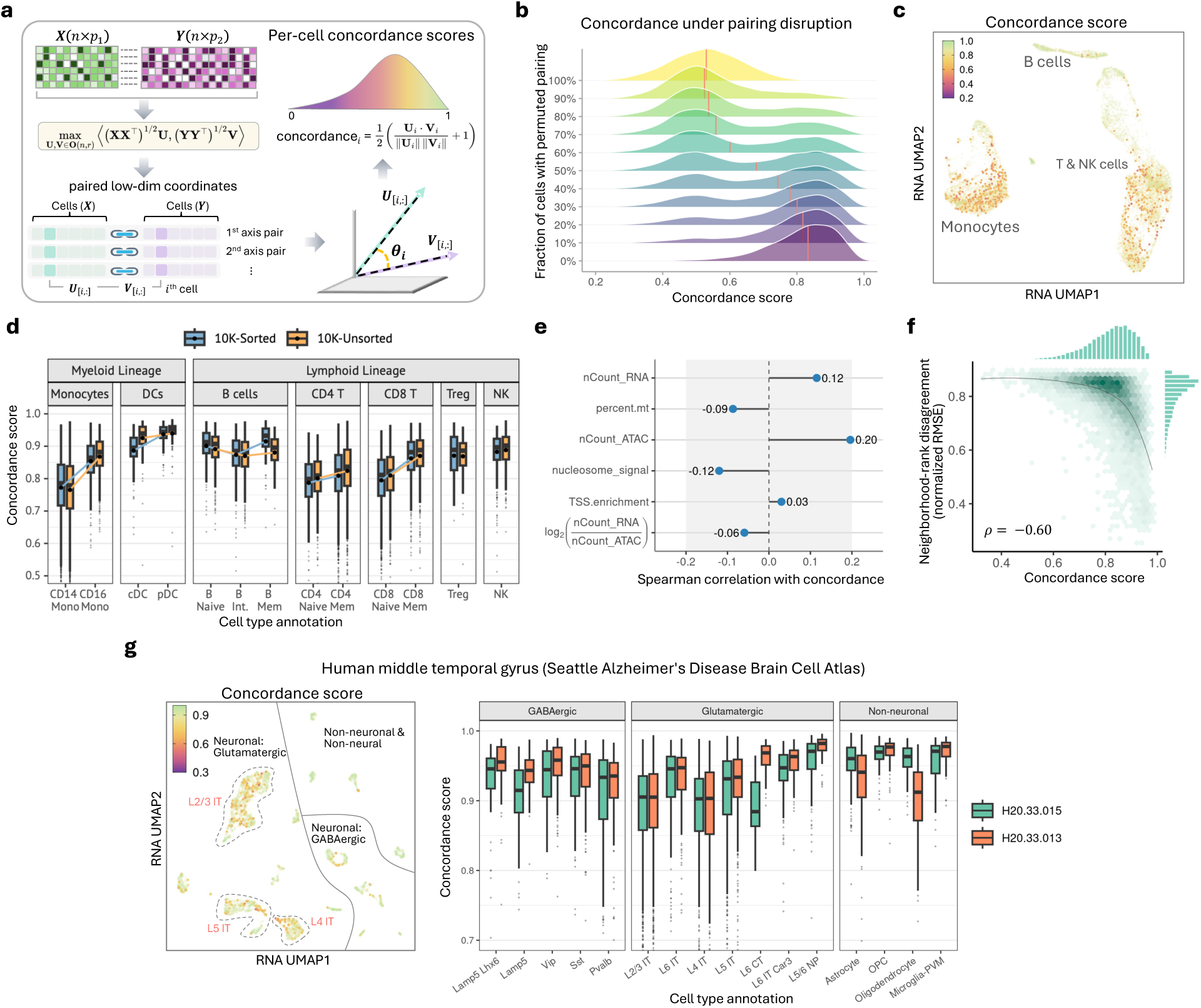
scOnion quantifies cell-specific concordance across modalities. **a**, Schematic of concordance score computation. Cell-paired matrices are jointly optimized by the scOnion objective to yield modality-specific spectral embeddings with explicitly matched axes. For each cell, the cosine similarity between its paired spectral coordinates, rescaled to [0, 1], is its concordance score, which measures how similar the cell’s representations are across the two jointly learned latent spaces (Methods). **b**, Concordance distributions in the unsorted PBMC 10K sample (*n* = 10,489 cells) under progressive pairing disruption, with the RNA identities of 0% to 100% of cells permuted in 10% increments (vertical lines, medians). The lower-scoring mode comprises the artificially un-paired cells, which separate from the true-paired cells. **c**, Cells colored by concordance score on the RNA-derived UMAP of the granulocyte-sorted PBMC 10K sample (*n* = 10,280 cells). Low-scoring cells (red and orange) are enriched within specific monocyte and T cell subpopulations. **d**, Per-cell-type concordance in the sorted and unsorted PBMC 10K samples, two independent datasets with comparable cell-type composition. The characteristic high- and low-scoring populations are preserved across the two samples (one-sided Wilcoxon rank-sum tests, Supplementary Table 4). **e**, Association between per-cell concordance and six standard per-cell quality-control metrics (RNA and ATAC library size, percent mitochondrial reads, nucleosome signal, TSS enrichment, and the log_2_ RNA/ATAC depth ratio), quantified by two-sided Spearman correlation in the sorted PBMC 10K sample (*n* = 10,280 cells). Associations are weak, the strongest being with ATAC library size (*ρ* = 0.20). **f**, Per-cell concordance versus cross-modality neighborhood-rank disagreement in the sorted PBMC 10K sample. Disagreement is the normalized-rank root-mean-square error be-tween a cell’s RNA and ATAC neighbor rankings over the union of its two 25-nearest-neighbor sets, with ranks capped at 125 and scaled by the maximum (Methods). Higher-concordance cells show lower neighborhood disagreement (two-sided Spearman *ρ* = −0.60). Marginal histograms show the two distributions. **g**, Left, concordance scores of a human middle temporal gyrus (MTG) multiome sample (SEA-AD donor H20.33.015) on the RNA-derived UMAP. Right, subclass-level concordance distributions for two donors (H20.33.015 and H20.33.013), with largely consistent high- and low-scoring subclasses across individuals. Panels d and g contain box plots of per-cell concordance by cell type (d) or subclass (g). The center line is the median, the box limits are the first and third quartiles (interquartile range, IQR), and the whiskers extend to the most extreme value within 1.5× the IQR. Outliers beyond the whiskers are plotted as individual points. Statistical tests and sample sizes are reported in Methods. Source data are provided with this paper.

To evaluate whether concordance captures the degree of agreement between modalities, we progressively disrupted cross-modal pairing in the unsorted PBMC 10K multiome sample [15] by randomly permuting RNA profiles of an increasing fraction of cells. As the permuted fraction increased, concordance scores shifted toward lower values and became bimodal at intermediate perturbation levels, with disrupted cell pairs moving toward low concordance while intact pairs remained highly concordant (Fig. 2b). Per-cell concordance distinguished permuted from intact cells with AUROC values of 0.87–0.97 across all permutation fractions and all four PBMC samples, showing that concordance reliably detects disrupted cross-modal pairing (Extended Data Fig. 1a). We complemented the permutation control with a noise-free simulation in which the two modality representations of each cell differ by a known rotation angle (Methods). Concordance decreased monotonically with this angle, from a median of 0.98 for cells rotated by 5^◦^ to 0.79 for cells rotated by 45^◦^, and tracked the true per-cell angle with a Spearman correlation of 0.90 (Extended Data Fig. 1c). Because the rotation angle was assigned independently of cell group, this correlation was the same within groups as across them, indicating that concordance captures graded cross-modality difference at the level of individual cells, independent of group identity. We next computed concordance scores in real datasets. In the granulocyte-sorted PBMC 10K multiome sample [16], low-concordance cells localized to discrete regions of the RNA-derived UMAP (Fig. 2c), indicating that cross-modality agreement varies systematically across cell populations. For example, CD14 monocytes showed lower concordance than CD16 monocytes (median 0.79 versus 0.86), and naive CD4 and CD8 T cells tended to score lower than their memory counterparts (0.80 versus 0.82 and 0.81 versus 0.88; Fig. 2d). These cell-type- and state-specific patterns were reproducible across four independent PBMC samples, with the same high- and low-concordance populations recurring across samples (Fig. 2d and Supplementary Fig. 2a,b). Together, these reproducible differences suggest that concordance may capture biologically meaningful variation in how cell states are coordinated across modalities. However, technical properties also vary across cell populations, so we tested how strongly concordance tracks per-cell data quality. Its correlations with standard per-cell QC metrics were weak, with |*ρ*| ≤ 0.20 across the six metrics tested (Fig. 2e). Sequencing depth showed the strongest of these correlations. Down-sampling both modalities to a common per-cell depth and refitting scOnion preserved the cell-type concordance ordering and removed the concordance-depth association (Extended Data Fig. 1b and Methods), indicating that sequencing depth alone does not explain the observed concordance differences.

In contrast, concordance was associated with cross-modality neighborhood agreement, with higher-concordance cells showing lower neighborhood-rank disagreement (normalized RMSE, Spear-man *ρ* = −0.60, Fig. 2f) and higher Jaccard overlap (Spearman *ρ* = 0.48, Supplementary Fig. 2c and Methods), indicating that it captures local cross-modality structure but is not fully explained by any single conventional neighborhood statistic. To relate concordance to existing per-cell quantities, we compared it with two independent measures, the synchrony score of Tilted-CCA [6], a per-cell measure of how strongly the two modalities share local structure, and the RNA modality weight of weighted-nearest-neighbor integration [8], a per-cell measure of which modality dominates locally. Within each immune cell type, concordance was moderately correlated with Tilted-CCA synchrony (Spearman *ρ* = 0.28 to 0.46) but weakly and inconsistently correlated with the WNN modality weight (Extended Data Fig. 1d,e and Methods). Within finer cell types the correlation with synchrony was indistinguishable from zero in naive B cells and conventional dendritic cells, so the two measures overlap partially and are not interchangeable (Methods). We next examined a human middle temporal gyrus (MTG) multiome dataset [17], where cellular heterogeneity reflects neuronal and glial subclass structure, providing a distinct biological setting from the immune activation and lineage state. Here, low-concordance cells again localized to regions of the RNA-derived UMAP and showed reproducible subclass-level differences across two donors (Fig. 2g). Relative to PBMC, the MTG dataset showed higher median concordance across individuals (0.92 and 0.93 versus 0.83 to 0.87 in the four PBMC samples, Supplementary Fig. 2a), showing that overall concordance can differ substantially across biological systems.

### 2.3 scOnion’s joint embedding retains modality-strong structure for concordance-guided subtyping

scOnion constructs a joint cell embedding by concatenating the paired modality-specific rank-*r* coordinates **U** and **V** learned from the scOnion objective. Given both coordinates, scOnion defines a 2*r*-dimensional embedding **W** = [**U**, **V**] that provides a single integrated coordinate system for downstream cell-level analysis, including nearest-neighbor graph construction, visualization, cell clustering, subtype refinement, and marker identification (Fig. 3a). **W** retains separations that are strongly represented in either modality, including structure supported mainly by one modality. To evaluate how union- and intersection-based methods behave under controlled signal imbalance, we simulated cell-paired profiles with two orthogonal factors, producing four ground-truth groups (Fig. 3b and Extended Data Fig. 2a). Each modality carried one factor as its dominant axis. The strength of the other factor was varied in increments, with modality-specific noise added to mimic real measurement differences (Methods). Across all signal-sharing combinations, the true cell-state structure always contained four groups, while the degree of cross-modality sharing varied. When both axes were strongly shared across modalities, all methods recovered the four groups. However, when one axis was weakly shared or supported by only one modality, intersection-based methods such as MOJITOO [7] and Tilted-CCA’s common embedding [6] recovered mainly the remaining shared axis, whereas MOFA2 [9] and MIRA [10] were less sensitive to this imbalance and WNN [8] behaved asymmetrically across reciprocal settings, consistent with its reported sensitivity to modality bias [18]. When shared signal was weak for both axes, only scOnion and MIRA retained all four separations, indicating stronger union-preserving behavior under extreme imbalance. scOnion achieved this with a linear construction, whereas MIRA relies on a more complex deep-learning model. In practice, scOnion built the joint embedding of the 10,333-cell granulocyte-sorted PBMC multiome in 4.5 minutes on a single CPU core from the processed matrices, about 28 times faster than MIRA run from raw counts, and with no GPU required (Methods). Scoring each method against the true group label by adjusted Rand index (ARI) quantified this behavior across all 25 signal-sharing conditions, with scOnion remaining near-perfect as shared signal weakened while the intersection-based methods declined (Extended Data Fig. 2b). Shared structure is captured by the top, most aligned directions in **U** and **V**, whereas strong modality-uneven separations can remain represented in less-aligned directions. On the granulocyte-sorted PBMC 10K multiome the scOnion joint embedding recovered the reference cell types better than six alternative integration methods on all three metrics scored (ARI 0.64 versus at most 0.53, Extended Data Fig. 3a,b and Methods). We complemented this reference-based comparison of the same seven methods on an independent dataset, the NeurIPS 2021 bone-marrow multiome (24,997 cells), using annotation-free metrics designed to reward embeddings that preserve both modalities (Methods). scOnion retained the most local structure at every clustering resolution and every neighborhood size (Extended Data Fig. 3c–e). Because union-preserving embeddings can contain both jointly supported and modality-uneven signals, scOnion interprets **W** together with per-cell concordance, which provides a local indicator of whether retained structure is broadly supported by both modalities or primarily by one. Major cell classes are often supported by both RNA and ATAC, whereas finer state or subtype differences may be clearer in one modality than the other. Consistent with this expectation, the paired modality-specific coordinates **U** and **V** showed similar coarse organization of major immune populations in the granulocyte-sorted PBMC 10K sample, with local differences within several lineages (Fig. 3c). The concatenated joint embedding preserved this coarse cell-type organization and revealed localized regions of low concordance (Fig. 3d). Next, we used this concordance-guided view to analyze subtypes within coarse immune compartments. After assigning major cell classes in the joint embedding, we subclustered cells within each compartment and quantified how strongly the resulting scOnion-derived subtype labels were separated in RNA and in ATAC (Fig. 3e). Across compartments, concordance was associated with the mean within-modality silhouette score across RNA and ATAC (Spearman *ρ* = 0.68, Supplementary Fig. 3a), indicating that high-concordance populations tend to contain subtype separations supported by both modalities. In contrast, low-concordance compartments were enriched for uneven support, where a retained subtype separation was clearer in one modality than the other. For example, within low-concordance naive CD4 T cells, one subcluster showed stronger silhouette separation in ATAC than in RNA (silhouette 0.29 versus 0.07, Fig. 3f), indicating that the joint embedding retained a modality-uneven separation without forcing agreement between layers. High-concordance compartments such as memory CD8 T cells instead showed subtype separation supported by both modalities (silhouette 0.31 and 0.34; Fig. 3f and Supplementary Fig. 3b–d). The joint embedding therefore retains candidate subtype structure, while concordance distinguishes separations supported by both modalities from those supported more strongly by one.

**Figure 3:**
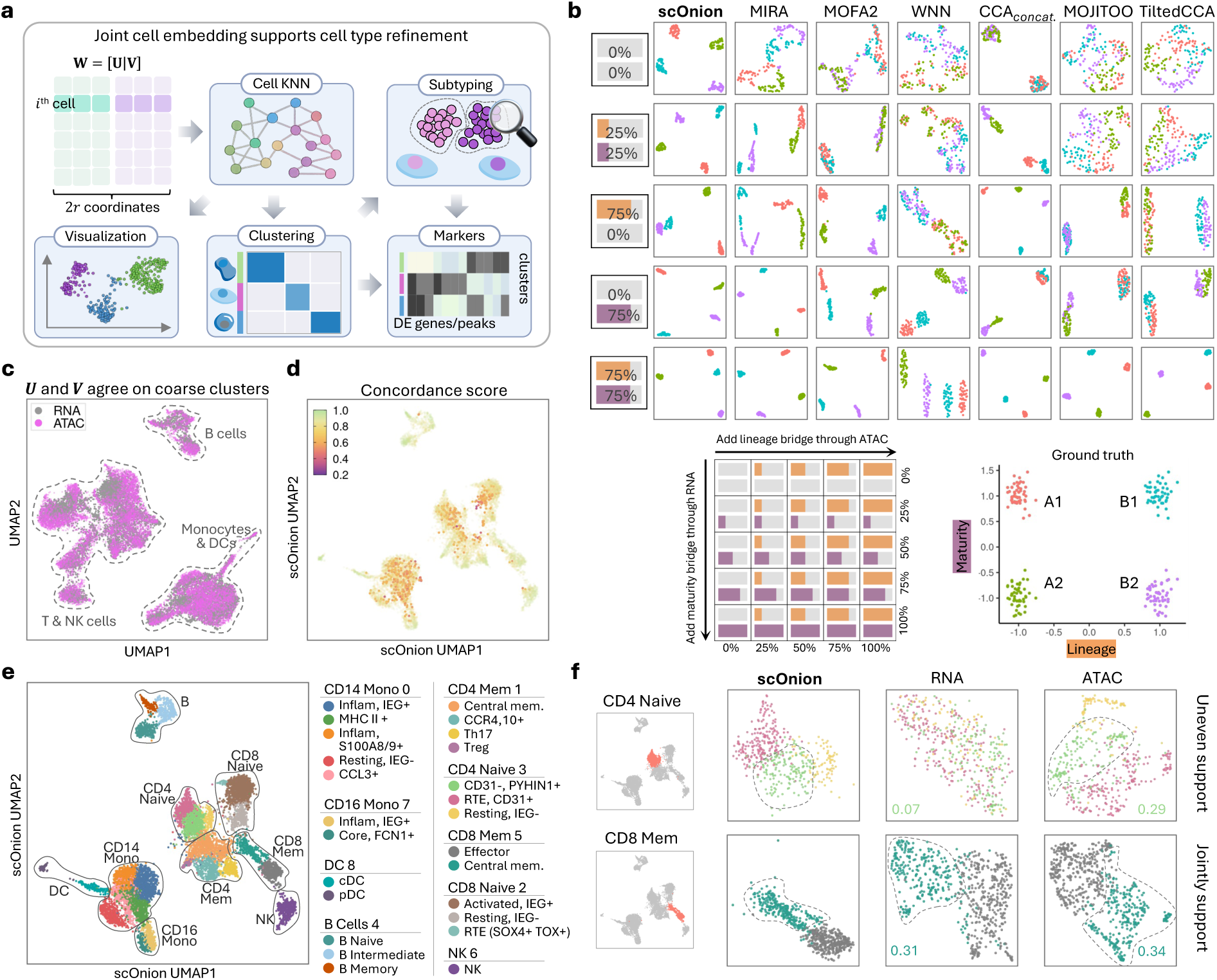
scOnion joint cell embedding retains modality-strong structure for concordance-guided subtyping. **a**, Overview of the scOnion joint cell embedding construction. scOnion learns a 2*r*-dimensional cell embedding **W** by concatenating the paired optimizers **U** and **V**, which yields a shared cell *k*-nearest-neighbor graph for visualization and clustering. Per-cell concordance scores (Fig. 2) prioritize clusters and neighborhoods for modality-specific subtyping, after which cluster markers are identified by differential expression and accessibility analysis. **b**, Simulation comparing seven joint-embedding methods (columns) across five representative regimes of cross-modality signal sharing (rows), with rows 3 and 4 the asymmetric cases in which a bridging factor is added in only one direction. Two latent factors, lineage and maturity, dominate opposite modalities. The full 5×5 design, its adjusted Rand index, and the four-group ground truth are given in Extended Data Fig. 2a,b and Methods. Each panel is a UMAP with identical parameters. In each row label the upper number is the ATAC signal ratio and the lower the RNA ratio. **c**, Modality-specific coordinates **U** (RNA) and **V** (ATAC) show similar coarse organization of the major immune populations in the sorted PBMC 10K sample, overlaid on a shared UMAP of the stacked U and V coordinates. **d**, Per-cell concordance on the scOnion joint embedding. Low-concordance cells (red and orange) localize to discrete regions of the embedding. **e**, Subtype annotation within coarse compartments using the joint embedding, resolving fine immune states across myeloid and lymphoid lineages. **f**, Two example compartments shown in the scOnion joint, RNA-only, and ATAC-only embeddings, annotated with within-modality silhouette scores. Low-concordance CD4 naive shows uneven support, separating more strongly in ATAC (0.29) than RNA (0.07), whereas high-concordance CD8 memory is jointly supported in both modalities (0.31 and 0.34). The RNA-only and ATAC-only panels are the Seurat principal-component and Signac LSI reductions of the same object, with UMAPs recomputed on the compartment’s cells and the depth-tracking first LSI component discarded, whereas the scOnion panel is the corresponding region of the joint UMAP computed once over all cells with the released package (Methods). Silhouette scores are descriptive, computed in each modality’s own reduction (dimension ranges in Methods). Panels c to f use the granulocyte-sorted PBMC 10K sample (*n* = 10,280 cells). Statistical tests and sample sizes are reported in Methods. Source data are provided with this paper.

### 2.4 scOnion provides an uncertainty-aware gene–peak coupling readout supported by orthogonal evidence

Unlike cells, which largely recur as corresponding cell types across modalities, features lack a clear one-to-one cross-modal correspondence, and genes and peaks are related only through indirect priors such as genomic proximity. As a result, aligning features across modalities is substantially harder than integrating cells (Supplementary Fig. 4a). scOnion addresses this challenge by leveraging the cross-modal correspondence provided by cell pairing. Because the same cells anchor both modalities in a common embedding, features from different layers can be compared through their patterns across this shared cell space.

Projecting each modality’s features onto the joint cell embedding places genes and peaks in a shared low-dimensional space, where genes and peaks with similar variation across cells have similar coordinates (Methods). We score each candidate pair by a weighted inner product of their coordinates, which measures how strongly they align along the axes shared by the two modalities. Each feature coordinate is a sum of per-cell contributions, so restricting it to a selected population yields a context-specific coupling map without refitting the embedding. The score is defined for any gene–peak pair and can be applied to pairs on different chromosomes, although the analyses here focus on *cis* associations. Each score reduces to an inner product of low-dimensional feature coordinates, so all links follow from a single matrix multiplication. Because the score is not tied to a genomic window, the same fit can rank broad candidate sets without fitting or testing each pair separately (Fig. 4a). However, a raw covariation score is difficult to compare across pairs, both because nearby gene–peak pairs covary more strongly as a result of genomic proximity, and because a score estimated from few cells is less reliable than one supported by many. scOnion accounts for both by subtracting the mean score among pairs in the same genomic distance range and dividing by a per-link standard error (SE), yielding a distance-controlled, uncertainty-aware score (*z*_dev_, Methods). Because the linkage score is bilinear in per-cell contributions, each leave-one-out score follows from an algebraic update of the full score, so a per-link jackknife SE can be computed in closed form without re-fitting the embedding. Other methods that provide per-link significance, whether correlation methods such as Signac [11] and FigR [12] or the graph-embedding method scGLUE [14], obtain it from permutation or embedding-shuffling nulls, whereas scOnion provides a per-link jackknife SE in closed form at negligible additional cost. This jackknife conditions on the fitted low-rank embedding and quantifies how sensitive each link score is to individual cells. Measured by the absolute gene–peak Spearman correlation across cells, summarized as the median over genes, peaks ranked most highly by scOnion were more co-active with their target gene than distance-ranked or random peaks, with the gap largest for the top-ranked peaks (Supplementary Fig. 4b).

**Figure 4:**
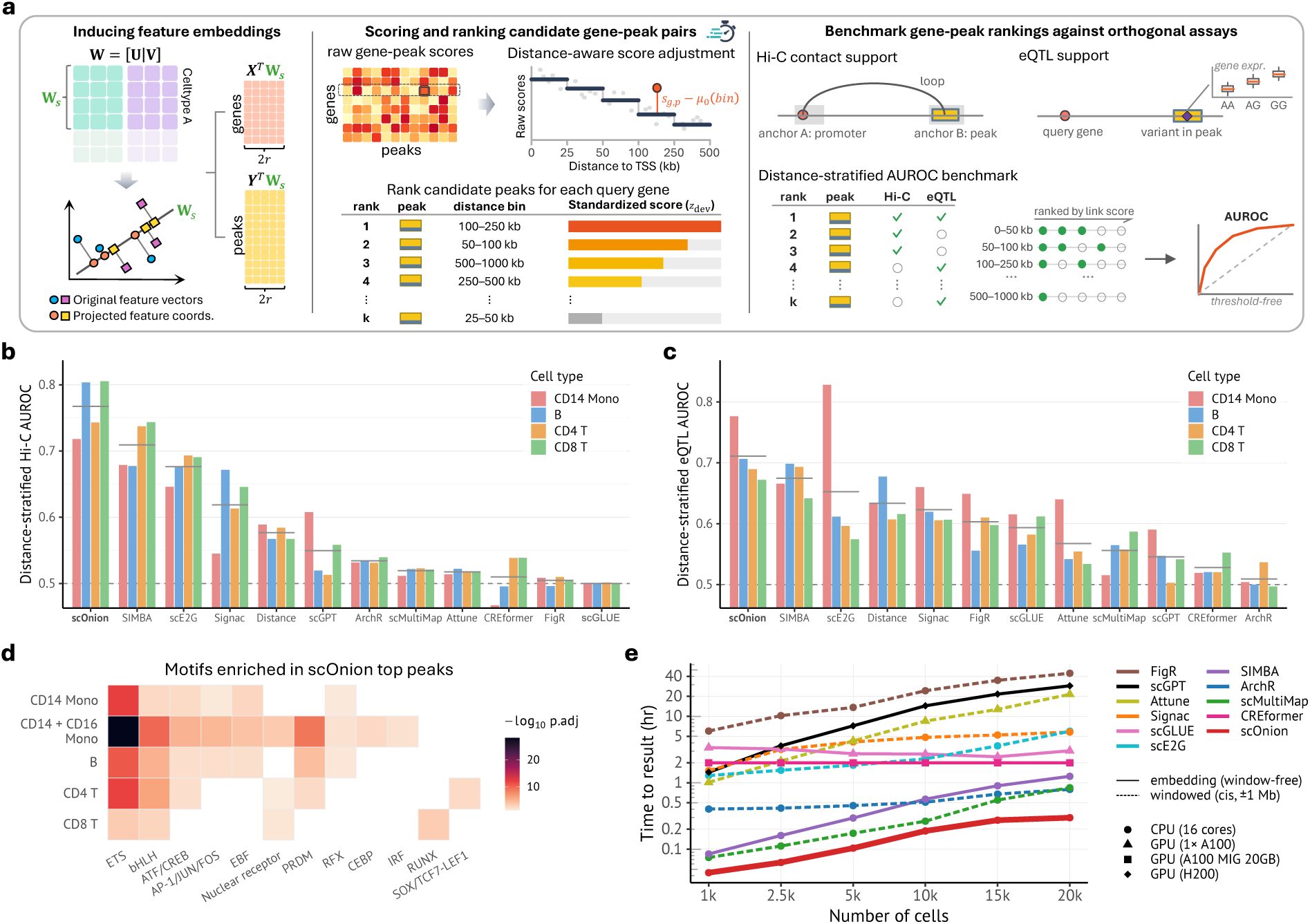
scOnion feature embedding recovers validated regulatory element–gene (rE2G) links. **a**, Overview of feature-level linkage. Modality-specific features are projected into the joint cell embedding to induce comparable gene and peak coordinates (Methods). A weighted inner product between gene and peak coordinates scores each candidate pair, and a distance-aware standardization (*z*_dev_) compares each score against a distance-matched background before ranking. Rankings are benchmarked against orthogonal assays, Hi-C contact and eQTL, by distance-stratified AUROC. **b**, Recovery of validated Hi-C promoter–peak contacts, measured as the threshold-free area under the receiver operating characteristic curve (AUROC) for scoring true contacts above non-contacts. AUROC is computed within gene and genomic-distance strata and pooled (distance-stratified, Methods), so proximity alone cannot inflate it. The dashed line at 0.5 is chance. For each method (x-axis) the four bars are the four PBMC cell types (B, CD4 T, CD8 T, CD14 monocyte; legend) and the gray tick marks their mean. *Distance* is a baseline that ranks peaks by distance to the TSS alone. **c**, Recovery of cell-type-matched fine-mapped eQTLs, scored as in panel b. A peak is a positive for a gene when it contains a fine-mapped eQTL variant of that gene (Methods). Cell types, per-method bars, mean ticks and the *Distance* baseline are as in panel b. **d**, Transcription-factor motif families enriched in the scOnion top-linked peaks of each cell type (− log_10_ adjusted *P*, one-sided hypergeometric test against a GC-matched background; Methods). Lineage-informative families are shown. Generic families, which carry the strongest raw enrichment, are in Supplementary Fig. 4c. **e**, Time to result versus number of cells for eleven methods, each producing a *cis* gene–peak linkage table (candidate gene–peak pairs within 1 Mb of the TSS), on CPU (16 cores) or a single GPU (Methods). Values are elapsed hours on a single hardware unit, and methods are marked as embedding methods, which need no window, or windowed *cis*-only methods. scGPT runtimes are extrapolated from a per-cell-type measurement and CREformer’s are independent of cell number (pseudobulk inference), whereas all other points are measured. scOnion had the shortest measured time to result at every cell number evaluated. Statistical tests and sample sizes are reported in Methods. Source data are provided with this paper.

To assess whether these scores prioritize gene–peak pairs supported by orthogonal evidence, we benchmarked scOnion against eight published gene–peak linkage methods, two large single-cell foundation models (scGPT [19] and CREformer [20]), and a trivial genomic-distance-to-TSS baseline in four main PBMC cell types (B, CD4 T, CD8 T and CD14 monocytes), using two sources of support. Against cell-type-matched promoter–peak contacts from intact Hi-C (ENCODE4 [21, 22]), scored by a distance-stratified, threshold-free AUROC, scOnion achieved the highest AUROC in all four cell types and in all six peak–TSS distance bins (mean AUROC 0.77 versus 0.71 for the next-best method; Fig. 4b and Extended Data Fig. 4a). Against a fully independent genetic source, fine-mapped cell-type eQTLs from OneK1K [23], scOnion was again among the strongest methods (highest mean AUROC, 0.71), with the supervised method scE2G [24] leading in CD14 monocytes (0.83 versus 0.78; Fig. 4c). Both foundation models export pretrained gene and peak embeddings, and we scored each candidate pair by the inner product of its two embeddings (Methods). Neither improved on the distance baseline on either benchmark, even though each is pretrained on far larger corpora than any dedicated linkage method, scGPT on over 33 million cells and CREformer as a three-billion-parameter model trained on bulk and single-cell multiomic data. In this benchmark, pretraining scale alone therefore did not yield stronger feature linkage than a linkage-specific score derived directly from the paired data. An ablation showed that the distance control and per-link SE underlie scOnion’s advantage, raising Hi-C AUROC over the raw score in all four cell types by 0.06 to 0.21 (Extended Data Fig. 4b).

As an independent validation, the peaks most strongly linked to each cell type’s marker genes were enriched for the binding motifs of that lineage’s transcription factors (Fig. 4d and Methods), recovering canonical regulators such as *TCF7* and *LEF1* in naive CD4 T cells and RUNX factors in memory CD8 T cells. Because the score is distance-controlled, this enrichment is carried by distal REs, not by promoter-proximal accessibility alone (Supplementary Fig. 4c).

scOnion was the fastest of the evaluated methods at every dataset size from 1,000 to 20,000 cells on the granulocyte-sorted PBMC multiome, taking 18 minutes at 20,000 cells compared with 47 minutes for the next-fastest method and 44 hours for the slowest (Fig. 4e and Methods). Unlike the two foundation models, it requires no pretraining. On the NeurIPS bone-marrow multiome, which has a comparable feature space of 116,490 peaks, scOnion scaled to all 69,249 cells in 32 minutes on the same 16 CPU cores (Extended Data Fig. 4c,d and Methods).

### 2.5 scOnion characterizes transcription-epigenome dynamics in B-cell fate bifurcation

As naive B cells activate and commit to one of two fates, antibody-secreting plasmablasts (PB) or germinal-center B-cell precursors (preGCBC), they remodel their transcriptome and chromatin in the differentiation process [25]. The PB fate engages the antibody-secretory program driven by *PRDM1*, *IRF4* and *XBP1* [26], whereas the preGCBC fate sustains B-cell identity and the germinal-center program under *BCL6*, *PAX5* and *BACH2* [27]. We therefore asked how cross-modal concordance and RE–G coupling change as cells commit to the two fates. To understand these cross-modality dynamics, we applied scOnion to an IGVF multiome time course of an *in vitro* system [25] that captures this bifurcation as a temporal continuum (Fig. 5a). When cells were ordered along each branch by pseudotime, concordance was high in the naive and terminal states but fell to a minimum near fate commitment along both the PB and preGCBC trajectories, reaching 0.81 along the PB branch and 0.78 along the preGCBC branch in donor 2, with both minima near pseudotime 0.5 (Fig. 5b, Extended Data Fig. 5a, and Methods). This decline is consistent with uneven RNA and ATAC remodeling during the transition [12]. The dip was reproducible across donors, with the minimum near pseudotime 0.5 in both (Extended Data Fig. 5a), and it remained after regressing per-cell concordance on sequencing depth, cell-cycle and immediate-early signature scores (Extended Data Fig. 5b). To test whether such concordance dynamics also arise in other differentiation programs, we applied scOnion independently to a human bone-marrow CD34+ hematopoietic progenitor multiome [28] and traced concordance along developmental pseudotime for four differentiation lineages. Concordance again varied along pseudotime in all four lineages, with similar trajectories across the two replicates (Extended Data Fig. 5c and Methods).

**Figure 5:**
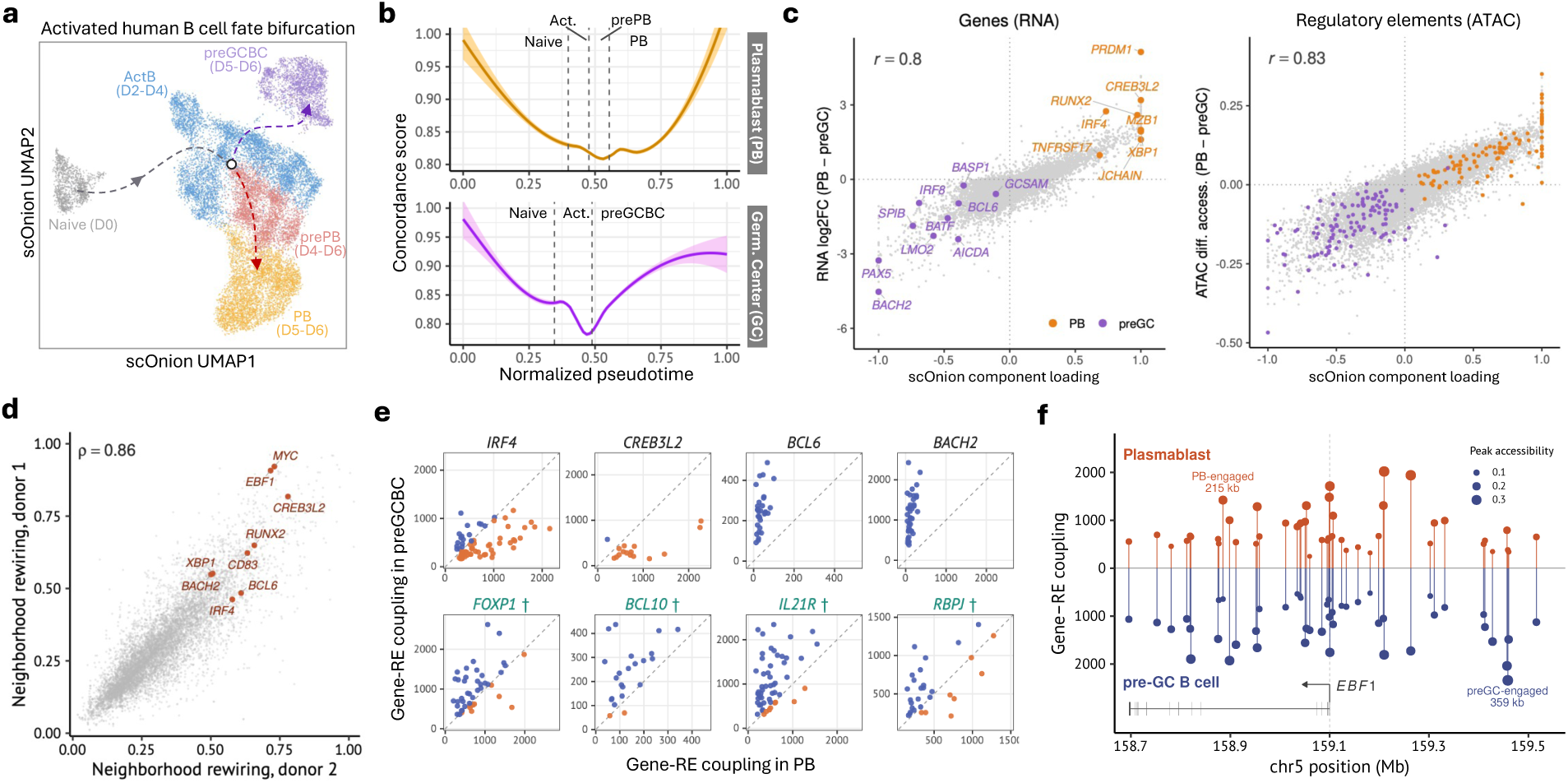
scOnion interprets an activated human B-cell fate bifurcation. **a**, scOnion joint-embedding UMAP of the activated human B-cell system, from naive cells through activated B cells (ActB) to the two divergent fates, plasmablast (PB, via prePB) and germinal center (GC, via preGCBC). **b**, Concordance along normalized pseudotime for the PB and GC lineages, shown as a loess fit with a 95% confidence band (Methods). Both lineages show a transient concordance decrease near fate commitment. **c**, scOnion fate-axis loading (component 2, x-axis) against RNA log_2_ fold change (left) and ATAC differential accessibility (right) between the PB and preGCBC fates. On the left, curated fate genes are colored by pole (PB, orange; preGCBC, purple) over a gray background of all genes. The 18 curated genes are eight plasmablast (*PRDM1*, *IRF4*, *XBP1*, *MZB1*, *TNFRSF17*, *CREB3L2*, *RUNX2* and *JCHAIN*) and ten pre-germinal-center (*BCL6*, *BATF*, *IRF8*, *SPIB*, *PAX5*, *BACH2*, *AICDA*, *LMO2*, *GCSAM* and *BASP1*). On the right, the twelve REs most strongly linked to each fate gene by *z*_dev_ are colored by the fate of their gene, over a random subsample of 40,000 peaks in gray. For display the loadings are divided by the 99.5th percentile of their absolute value, clipped to [−1, 1] and oriented toward the plasmablast pole, and the correlations are computed on the unscaled loadings. The loading tracks both differential signals (Pearson *r* = 0.80 across 15,352 genes and 0.83 across 167,404 peaks), so one scOnion axis orders both the fate genes and their linked REs consistently with the two fates. **d**, Per-gene neighborhood rewiring score in the two independent donors (two-sided Spearman *ρ* = 0.86 over 8,831 genes). Labeled genes are fate-associated genes that rewire strongly in both donors. Seven are highlighted in the original study, and *EBF1* and *RUNX2* are transcription factors outside that set. **e**, scOnion coupling of each candidate RE to its gene in PB (x-axis) and preGCBC (y-axis) for eight genes, the four canonical fate regulators *IRF4*, *CREB3L2*, *BCL6* and *BACH2*, which are differentially expressed between the fates, and four B-cell genes rewired without differential expression, *FOXP1*, *BCL10*, *IL21R* [35] and *RBPJ*. Each point is a candidate RE, colored by the fate in which it couples more strongly; the dashed line marks equal coupling in both fates. **f**, scOnion rE2G coupling across the *EBF1* locus (chr5) in PB (upward, red) and preGC (downward, blue); each candidate RE within 500 kb of the TSS is a lollipop at its genomic position, sized by accessibility and filled when it has an activated B-cell Hi-C loop to the *EBF1* promoter (open, no loop detected), with the *EBF1* gene model below. *EBF1* is expressed equally in both fates (log_2_ fold change −0.10) yet couples to largely different REs in each, its rewiring ranking at the 98th percentile in donor 2 and reproduced in donor 1. A PB-engaged element 215 kb from the TSS and a preGC-engaged element 279 kb away, both with Hi-C support, are labeled. Panels a to c, e and f use donor 2, and panel d compares the two donors. Statistical tests and sample sizes are reported in Methods. Source data are provided with this paper.

At the feature level, scOnion recovered the fate decision in the second component of the joint feature embedding. The first component explained the most variance (69.7% of the joint cell embedding spectrum) but mainly captured a technical axis of library size and detection depth (RNA library size *R*^2^ = 0.72, genes detected *R*^2^ = 0.71; Supplementary Fig. 5b,c). The second component (22.7%) instead separated the two fates, with gene loadings tracking the transcriptional difference between PB and preGCBC (RNA log2 fold change, *r* = 0.80 over *n* = 15,352 genes) and RE loadings tracking the corresponding difference in chromatin accessibility (*r* = 0.83 over *n* = 167,404 peaks), placing the PB regulators and their secretory effectors (*PRDM1*, *IRF4*, *XBP1*, *MZB1*, *JCHAIN*, *TNFRSF17*) opposite the germinal-center program (*PAX5*, *BACH2*, *BCL6*, *AICDA*, *LMO2*, *SPIB*; Fig. 5c). CREB3L2 also loaded toward the PB-associated end of this component and was reported by the original study as a central node of the plasmablast regulatory network [25], recovered here without fate labels. Together these two components accounted for 92% of the joint cell embedding spectrum (Supplementary Fig. 5b), so its structured variation reduced almost entirely to one technical axis and the fate axis.

We next used the same feature embedding to examine how each gene’s coupling to its REs is rewired between the two fates. Because scOnion scores every gene–element pair together with its uncertainty in closed form (Fig. 4), restricting the embedding to each fate’s cells regenerates the full coupling map at negligible cost, letting us scan rewiring genome-wide across both fates and both donors without refitting or permutation. We summarized each gene’s rewiring as a neighborhood rewiring score, the degree to which its ranking of nearby REs changes between the two fates (Methods). Per-gene rewiring was highly reproducible across two independent donors (Spearman *ρ* = 0.86, *n* = 8,831 genes, Fig. 5d). Canonical B-cell fate regulators tended to have higher rewiring scores than other genes (AUROC 0.77, *n* = 15 versus 8,816 background genes, one-sided Wilcoxon *P* = 1.7 × 10^−4^, Fig. 5d and Methods). Among these regulators, the strongest rewiring in both donors was at *MYC* and *EBF1*, followed by *CD83*, *XBP1*, *BACH2*, *BCL6* and *IRF4*, and the genome-wide screen additionally placed *CREB3L2* and *RUNX2*, two transcription factors outside the curated regulator set (Supplementary Table 6), among the most rewired genes (Fig. 5d). Rewiring was largely independent of differential expression (Spearman *ρ* = 0.05 against | log_2_ FC| across all 8,831 tested genes, Supplementary Fig. 5d), allowing scOnion to identify fate-associated genes with little change in expression. At each of the eight loci examined, a gene’s coupling to its genomically nearby REs varied in strength between the two fates, both for canonical fate drivers reported in the original study (such as *IRF4* and *BCL6*) and for genes that scOnion nominated but the study did not (such as *FOXP1* and *RBPJ*; Fig. 5e). *FOXP1* represses *PRDM1*, *IRF4* and *XBP1* and is downregulated in germinal-center B cells [29, 30], and *RBPJ* transduces the Notch2 signal that directs activated B cells away from the germinal center [31]. *EBF1* was expressed equally in PB and preGCBC yet coupled to distinct REs in each fate, a PB-engaged element 215 kb from the TSS and a preGCBC-engaged element 279 kb away, both supported by Hi-C loops, a switch invisible to differential expression (Fig. 5f). Together, concordance identified where RNA and chromatin coordination changed during lineage commitment, while RE–G rewiring identified fate-relevant genes not apparent from differential expression alone.

### 2.6 scOnion generalizes to CITE-seq protein–gene linkage

Existing methods that link REs to genes are mostly built around priors specific to the RNA–ATAC pairing, such as a candidate window of fixed genomic distance, a distance-decay weighting, or a transcription-factor motif model [5, 11, 12]. However, these priors might not have a counterpart for other modality pairs. A surface protein and a gene, for example, share no genomic coordinate, so neither a candidate window nor a motif model is defined. Beyond the known cognate relationship, linking surface proteins to other genes can reveal broader transcriptional programs associated with a surface phenotype. Most CITE-seq analysis methods integrate cells [8] or denoise abundances [32], and few link features [4, 33]. During feature linkage scOnion uses no prior or external information beyond the two modality matrices and their cell pairing, so the construction is not specific to a genomic coordinate and can in principle apply to other paired assays with matched cells. We applied scOnion to human PBMC CITE-seq, which measures RNA and surface-protein (ADT) abundance in the same cells [8] (Fig. 6a). The identical pipeline produced a per-cell concordance score and a joint embedding of cells, genes and proteins. Concordance varied systematically across cell types and was reproducible across four donors (Fig. 6b,c and Supplementary Fig. 6a), so the cross-modality agreement scOnion measures is not limited to the RNA–ATAC pairing. Permuting the RNA–ADT cell pairing collapsed concordance from its intact median near 0.72 to a floor near 0.53 in every donor (Extended Data Fig. 6a), showing that concordance captures the cross-modality information carried by the correct cell pairing, as in the RNA–ATAC setting. We then used the joint feature embedding and its linkage score to relate surface proteins to genes. We first used cognate protein–gene pairs as a positive control. Each surface protein is translated from exactly one gene, its cognate, so recovering that gene is a positive control. For every protein we ranked all genes by their coupling score to it and recorded the cognate’s rank, defining cognate recovery as the fraction of proteins whose cognate falls among the top-ranked genes (Methods). Cognate recovery is not expected to be perfect because post-transcriptional regulation and trafficking can weaken correspondence between a surface protein and its transcript. Nevertheless, cognate genes should rank above chance overall. scOnion recovered cognates at higher rates than raw RNA–protein correlation and correlation on totalVI-denoised expression, at the top-1, top-5 and top-20 cutoffs and in all four donors (top-5 recovery 24% versus 14% and 10% averaged over donors, Fig. 6d and Extended Data Fig. 6b). Denoising with totalVI lowered recovery in this dataset, suggesting that this denoising approach did not preserve the RNA–protein relationships the analysis aims to recover.

**Figure 6:**
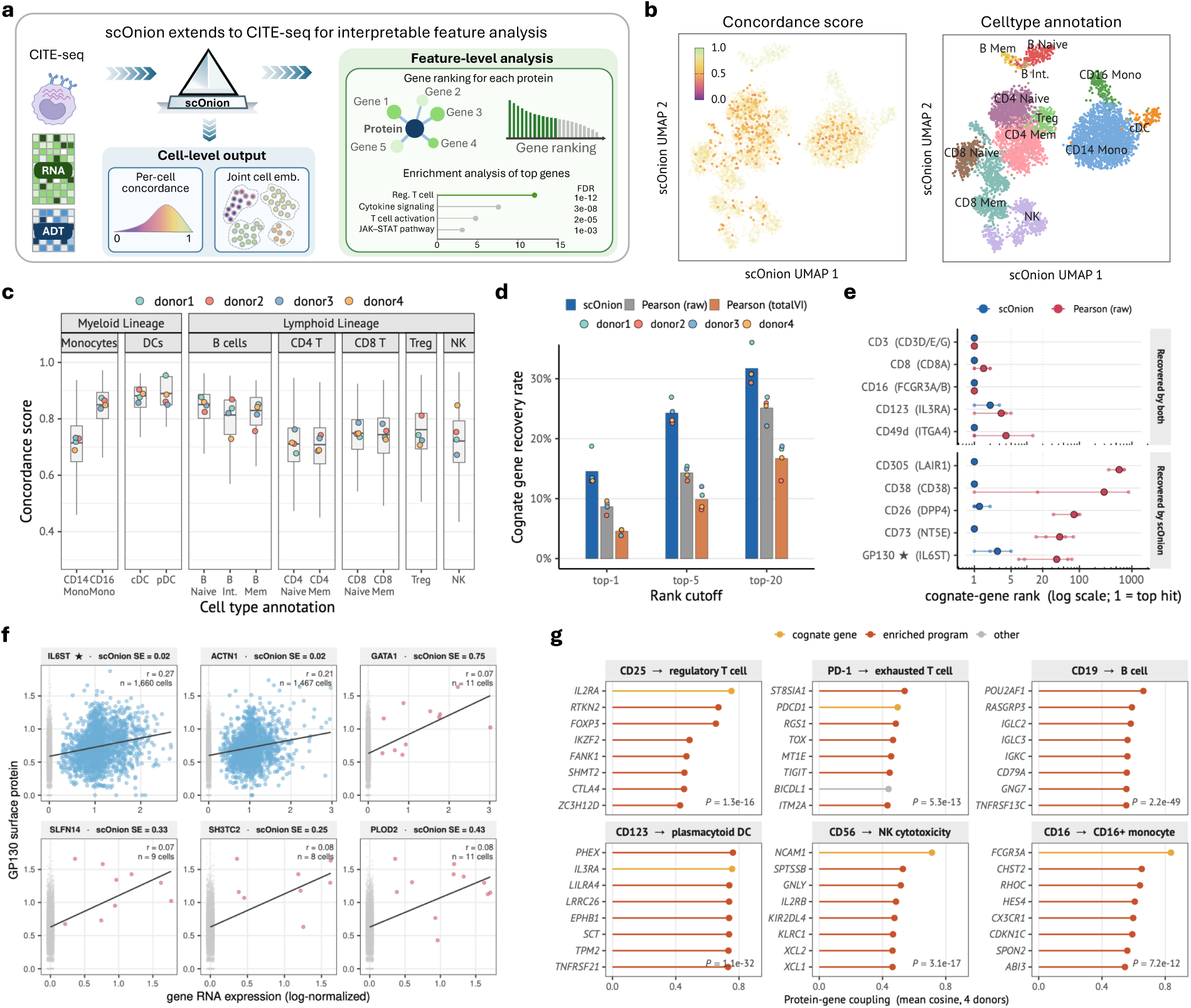
scOnion generalizes to CITE-seq for interpretable feature analysis. **a**, scOnion applies to CITE-seq (RNA and ADT) without modification, producing per-cell concordance and a joint cell embedding together with feature-level analysis of protein–gene coupling and an uncertainty-adjusted linkage score. **b**, scOnion joint-embedding UMAP colored by concordance (left) and cell-type annotation (right) for one donor of the human PBMC CITE-seq dataset. **c**, Per-cell-type concordance across four donors in myeloid and lymphoid lineages. **d**, Cognate-gene recovery rate, the fraction of 208 scorable proteins whose cognate gene ranks within the cutoff, at top-1, top-5, and top-20 for scOnion, Pearson correlation on raw counts, and Pearson correlation on totalVI-denoised values, across four donors. **e**, Cognate-gene rank for scOnion and raw Pearson correlation across ten representative proteins. For each protein every gene is ordered by its association with that protein and the position of the protein’s own transcript is plotted, rank 1 being best out of 15,237 to 15,648 genes per donor. Points are the mean over the four donors and the line spans the donor range. The ten are grouped into five whose cognate both methods place near the top and five whose cognate scOnion places near the top while raw correlation does not. The dotted guides mark the top-1, top-5 and top-20 cutoffs of Fig. 6d. CD26 denotes the CD26-2 antibody, one of two DPP4 clones in the panel. **f**, Per-cell GP130 surface protein versus gene RNA for the cognate *IL6ST*, a co-expressed gene (*ACTN1*), and four few-cell decoys (*GATA1*, *SLFN14*, *SH3TC2* and *PLOD2*), annotated with cell count and per-link standard error. Links supported by many cells, both the cognate and the co-expressed gene, have low standard error and are retained by the two-standard-error lower score (blue), whereas few-cell links carry large standard error and are deprioritized (red). **g**, Genes most strongly coupled to six surface proteins spanning the major PBMC lineages (CD25, PD-1, CD19, CD123, CD56, CD16) in the joint feature embedding (mean cosine over four donors, top eight shown), with the cognate gene in gold, genes in a significantly enriched program term in rust, and other coupled genes in grey. Over-representation of each protein’s top 50 coupled genes recovered its expected program (regulatory T cell, exhausted T cell, B cell, plasmacytoid dendritic cell, NK cytotoxicity, CD16-positive monocyte; Enrichr, Benjamini–Hochberg-adjusted *P* shown, significant in every donor; Methods). For CD25 and PD-1 the recovered genes include the state-defining transcription factors (*FOXP3*, *TOX*), not only co-markers. Panels b–g are human PBMC CITE-seq from four donors analyzed as biological replicates. Panel c is a box plot of per-cell concordance grouped by cell type. The center line is the median, the box limits are the first and third quartiles (interquartile range, IQR), and the whiskers extend to the most extreme value within 1.5× the IQR. Outliers beyond the whiskers are plotted as individual points. Statistical tests and sample sizes are reported in Methods. Source data are provided with this paper.

We next examined two properties that may contribute to this improved recovery, robustness to transcript dropout and per-link uncertainty estimation. The advantage over correlation was concentrated in abundant surface proteins whose cognate transcript is only sparsely detected. For CD305 (*LAIR1*), CD38 and CD73 (*NT5E*), raw correlation ranked the cognate gene well below the top in most donors, as low as rank 724, 867 and 77, whereas scOnion placed it first in all four donors (Fig. 6e). Across all four donors, scOnion recovered cognates over a much wider range of transcript sparsity than correlation, which required a transcript detected in many cells (Supplementary Fig. 6b). The low-dimensional cell-state representation stabilized cognate links involving sparse transcripts, which were often missed by raw correlation. scOnion also assigns each coupling a closed-form jackknife SE, an estimate of uncertainty that correlation- and denoising-based methods do not provide (Methods). For example, the cognate GP130–*IL6ST* coupling and other well-supported links were estimated with low uncertainty across roughly 1,660 cells, whereas sev-eral genes that outranked the cognate by the point estimate were supported by only 8 to 11 cells, had much larger per-link jackknife SE, and were deprioritized once the ranking accounted for this uncertainty (Fig. 6f). Across the four donors this moved the *IL6ST* cognate from ranks of 23 to 167 under the point estimate to ranks of 1 to 5. Across all proteins, cognate links carried systemati-cally lower SE than the high-magnitude few-cell links that topped the raw coupling, a median ratio of 3.5 (Extended Data Fig. 6c), so the SE distinguishes couplings supported broadly across cells from estimates sensitive to a few cells, as it does in multiome data. Ranking by the uncertainty-adjusted score gave a small but consistent improvement in cognate recovery over the point estimate, raising top-5 recovery from 20% to 24% in all four donors (Extended Data Fig. 6b and Supplementary Table 4). Finally, because scOnion ranks every gene against each protein, we asked whether the genes most strongly coupled to a protein recover nontrivial associations beyond the cognate. Across proteins spanning the major PBMC lineages, they formed coherent programs (Fig. 6g). For CD25 and PD-1 the top-coupled genes included the transcription factors and checkpoints that define the regulatory-T-cell state (*FOXP3*, *IKZF2* and *CTLA4*) and the exhausted-T-cell state (*TOX* and *TIGIT*), not only co-expressed surface markers, while CD19, CD123, CD56 and CD16 recovered their B-cell, plasmacytoid-dendritic-cell, natural-killer and CD16-positive monocyte lin-eage programs. Enrichment analysis returned the matching program as the leading term for each protein’s top-ranked genes (adjusted *P* between 2 × 10^−49^ and 7 × 10^−12^), reproduced in each of the four donors analyzed separately (Methods). These programs were recovered without cell-type labels, showing that the coupling ranking extends beyond cognate protein–gene pairs to broader cell-state-associated transcriptional programs.

## 3 Discussion

Here we present scOnion, a concordance-aware framework for paired single-cell multiomic data. From a single joint spectral decomposition of the two modalities, scOnion returns three connected outputs, a per-cell concordance score, a joint cell embedding, and a feature-to-feature coupling with a closed-form uncertainty estimate. Across immune, neural, and developmental datasets, its joint cell embedding retained cell-type and state separations that comparison methods represented less completely, and its feature coupling recovered links that agree with orthogonal chromatin-contact, genetic, and lineage motif evidence. scOnion additionally reports per-cell concordance, a measure of how strongly the two modalities agree in each cell, which varied with cell state and decreased transiently as cells committed to a fate.

Cell-paired multiomic assays directly establish cell correspondence across modalities. scOnion uses this correspondence to relate features that lack a natural cross-modal pairing by projecting each feature onto the joint cell embedding and comparing their coordinates in this shared space. The score is therefore not intrinsically tied to a predefined genomic window, though the *cis* analyses here use a ±1 Mb candidate set, and it makes no assumption specific to the assay pair. Because a feature’s coordinates in this space aggregate contributions across cells, the coupling for any chosen set of cells is recomputed directly from the global fit by restricting the sum to those cells, which gives context-specific couplings without re-estimating the embedding. The linear, low-rank construction of scOnion enables both interpretability and efficiency. Its objective has a closed-form solution, and its embedding axes carry a defined meaning, capturing shared and modality-specific structure. The same closed-form solution also yields derived quantities such as the per-link standard error at little additional cost. Correlation-based methods are interpretable but score pairs within a predefined genomic window that is adopted mainly for computational feasibility. Graph-embedding methods search across the full feature set for linkage discovery, but they depend on a trained model and are less directly interpretable. scOnion retains the interpretability of the former and the broad candidate scoring of the latter, and it estimates the uncertainty in closed form rather than through permutation-based null distributions. Two large pretrained foundation models scored near a genomic-distance baseline on the same benchmarks, suggesting that under the linkage scoring strategy evaluated here, their much larger pretraining corpora did not translate into more accurate feature linkage.

scOnion has several limitations. First, scOnion is currently designed to analyze a single multi-omic dataset or samples generated within the same experimental run. We have not examined its behavior when jointly analyzing multiple samples with batch effects. Second, the fit relies on several matrix operations of potentially large cell-by-feature matrices, so its computational cost grows with the number of cells. Our benchmarks reached 69,249 cells in 32 minutes and 455 GB of memory on 16 CPU cores, with wall-clock time growing approximately linearly with cell number and memory usage increasing more slowly. In practice, applying scOnion across several samples requires their ATAC peaks to be called jointly, so that every cell is represented in the same feature space. Third, validating rE2G links is intrinsically hard. No external source is an absolute gold standard, and each available proxy is informative but carries an assay-specific bias. As a complementary vali-dation, the protein–gene setting provides a known correspondence between each protein and its cognate transcript, although post-transcriptional regulation makes this relationship imperfect.

Several directions could strengthen the model and broaden its applications. scOnion maximizes a linear objective, and a kernel generalization could capture non-linear dependencies between features, but at some cost to the closed-form solution and its efficiency. The core construction also connects to the broader families of reduced-rank regression and generalized canonical correlation analysis, and developing this connection theoretically could provide formal tests of concordance and feature coupling beyond the descriptive statistics used here. The same construction extends to further paired assays and to trimodal assays such as DOGMA-seq [34], a generalization that will grow more valuable as trimodal profiling becomes more widely adopted. Extending the framework to jointly model multiple samples is another direction for future work.

We designed scOnion as a lightweight R package that runs on standard CPU hardware without pretraining or specialized accelerators. For exploratory analysis, its joint cell embedding yields well-separated cell types and states, and its per-cell concordance score identifies cells with greater disagreement between modalities, including cells passing through state transitions. Its feature linkage procedure scores large candidate sets without fitting a separate model for each pair and provides a cell-population-specific score and uncertainty estimate for each link. These rankings can prioritize RE-–G and protein–gene pairs for experimental follow-up, including perturbation studies and analyses linking noncoding variants to candidate target genes. Together, these components provide a computationally accessible framework for identifying where modalities agree across cells and prioritizing cross-modal feature relationships for further investigation.

## 5 Extended Data Figures

**Extended Data Figure 1:**
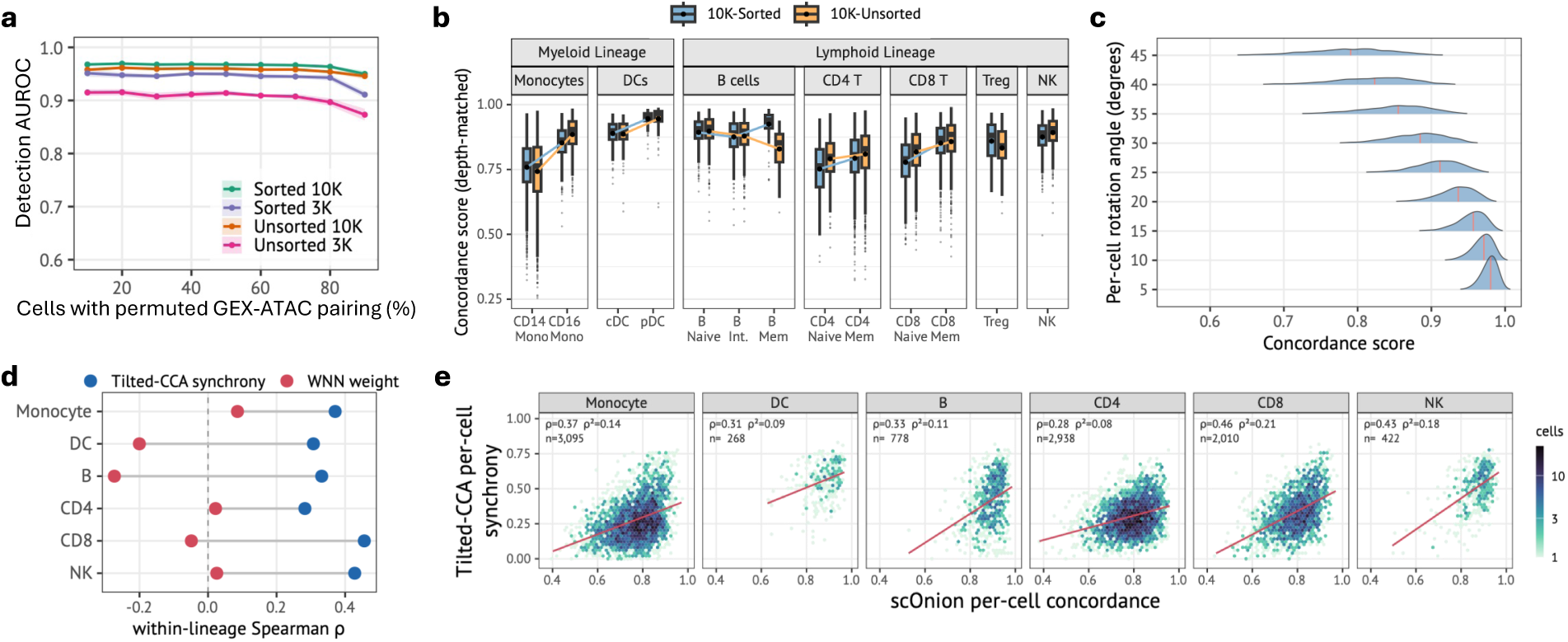
Per-cell concordance detects disrupted pairing, is insensitive to sequencing depth, and is distinct from existing per-cell measures. **a**, Detection curve for the permutation control of Fig. 2b. In each of four PBMC multiome samples the RNA and ATAC pairing was scrambled in an increasing fraction of cells (10 to 90%), and per-cell concordance alone was used to classify scrambled against intact cells, scored by AUROC (0.87 to 0.97 across samples and fractions; Methods). **b**, Per-cell-type concordance after down-sampling RNA and ATAC to a common per-cell depth and refitting scOnion, for the 10K sorted and unsorted samples. The Fig. 2d ordering is preserved at equalized depth (CD14 below CD16 monocytes, naive below memory T cells, pDC highest) and reproduces across both samples (Methods). **c**, Controlled simulation with no measurement noise, complementing the permutation control of panel a. Paired measurements were generated for 3,024 cells in six balanced groups, with the two modality representations of each cell separated by a known rotation angle assigned independently of group membership at nine levels from 5^◦^ to 45^◦^, each spread over a ±1^◦^ window (Methods). Ridges show the distribution of esti-mated concordance at each angle, pooled over ten simulations (*n* = 30,240 cells). Concordance falls monotonically as the angle increases, from a median of 0.98 at 5^◦^ to 0.79 at 45^◦^, and tracks the true per-cell angle with Spearman *ρ* = 0.90, the same value within groups as across them. **d**, scOnion concordance compared with two other per-cell cross-modality measures on the sorted PBMC 10K sample, the per-cell synchrony score of Tilted-CCA [6] and the per-cell RNA modality weight of weighted-nearest-neighbor (WNN) integration [8]. Each point is the within-lineage Spearman correlation of a measure with concordance across six immune lineages (Methods). Concordance is positively correlated with Tilted-CCA synchrony in every lineage (*ρ* = 0.28 to 0.46) but weakly and inconsistently correlated with the WNN weight (*ρ* = −0.27 to +0.09). **e**, Per-cell relationship behind the Tilted-CCA comparison in panel d, for the same six immune lineages (*n* = 9,511 cells). Hexagonal bins give the number of cells and the red line is an ordinary least-squares fit, with the within-lineage Spearman *ρ*, its square and the number of cells shown in each panel. Concordance and Tilted-CCA synchrony are only weakly related within every lineage, and within finer cell types the correlation is indistinguishable from zero in naive B cells (*ρ* = 0.01) and conventional dendritic cells (*ρ* = 0.14, 95% bootstrap intervals including zero; Methods). Panel b shows box plots and panel c shows density ridges; for the box plots the center line is the median, the box limits are the first and third quartiles (interquartile range, IQR), and the whiskers extend to the most extreme value within 1.5× the IQR. Outliers beyond the whiskers are plotted as individual points. Statistical tests and sample sizes are reported in Methods. Source data are provided with this paper.

**Extended Data Figure 2:**
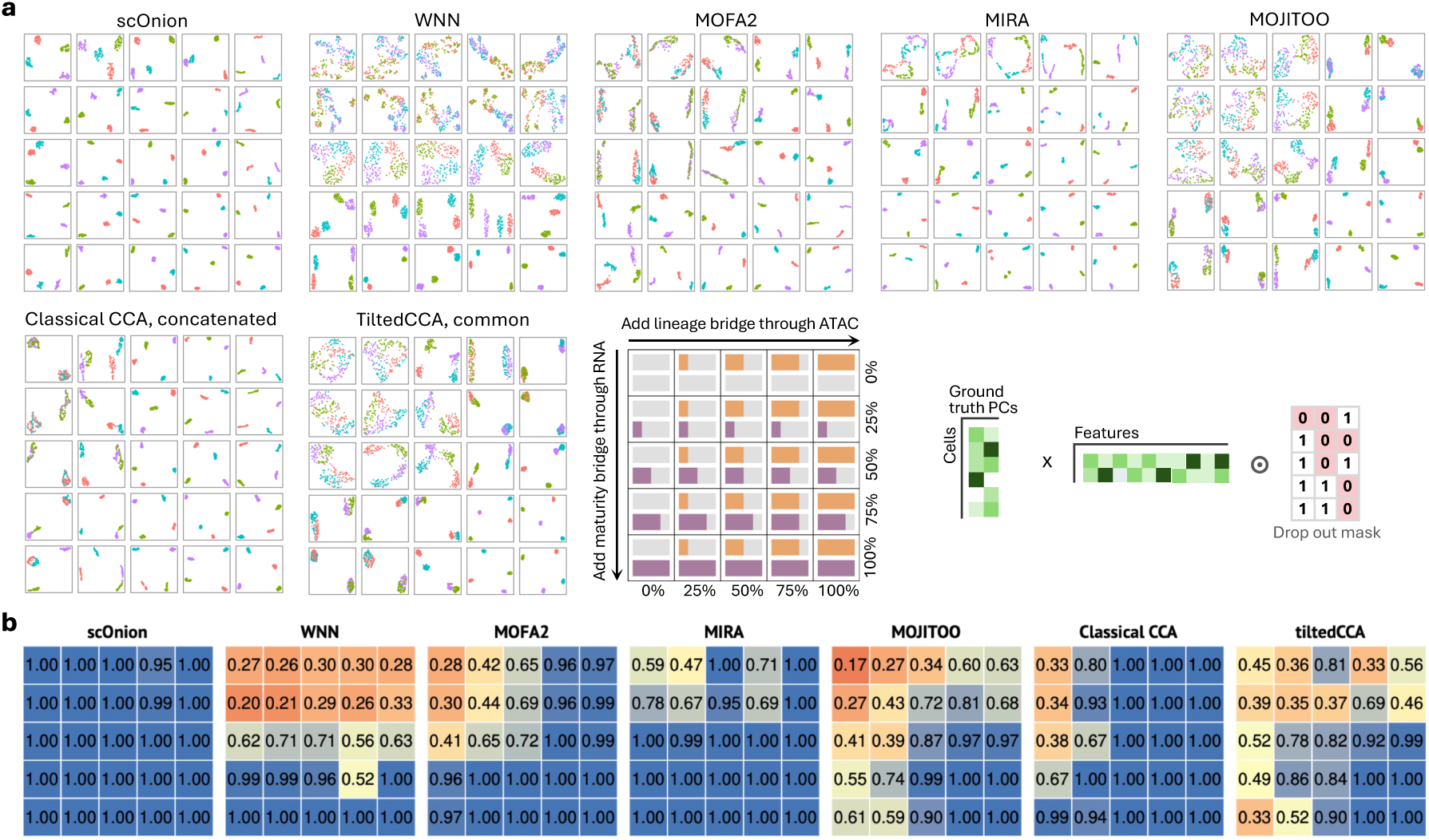
Simulation benchmark of joint cell embeddings under asymmetric cross-modality signal. **a**, Full 5×5 simulation grid for seven integration methods (Fig. 3b and Methods). Two orthogonal latent factors, lineage and maturity, generate paired RNA and ATAC measurements, with lineage dominant in RNA and maturity dominant in ATAC. The dominant factor of each modality is bridged into the other from 0% to 100% in 25% increments (schematic, lower right). Ground truth contains four groups (A1, A2, B1, B2). Each panel is a UMAP with identical parameters. scOnion and MIRA retain all four groups even when shared signal is weak, whereas intersection-oriented methods such as MOJITOO and the Tilted-CCA common embedding mainly recover the remaining shared axis. **b**, Quantification of the simulation grid in panel a. Adjusted Rand index (ARI) against the four-group ground truth for each of the seven methods across the same 5×5 bridge conditions (Methods). ARI is shown per condition because separability depends on the sharing regime. All methods reach ARI near 1 when both axes are strongly shared, but under weak or one-sided sharing the intersection-oriented methods decline while scOnion remains near 1 throughout (condition-averaged ARI compared with scOnion by two-sided paired Wilcoxon signed-rank test over the 25 conditions, Supplementary Table 4). Statistical tests and sample sizes are reported in Methods. Source data are provided with this paper.

**Extended Data Figure 3:**
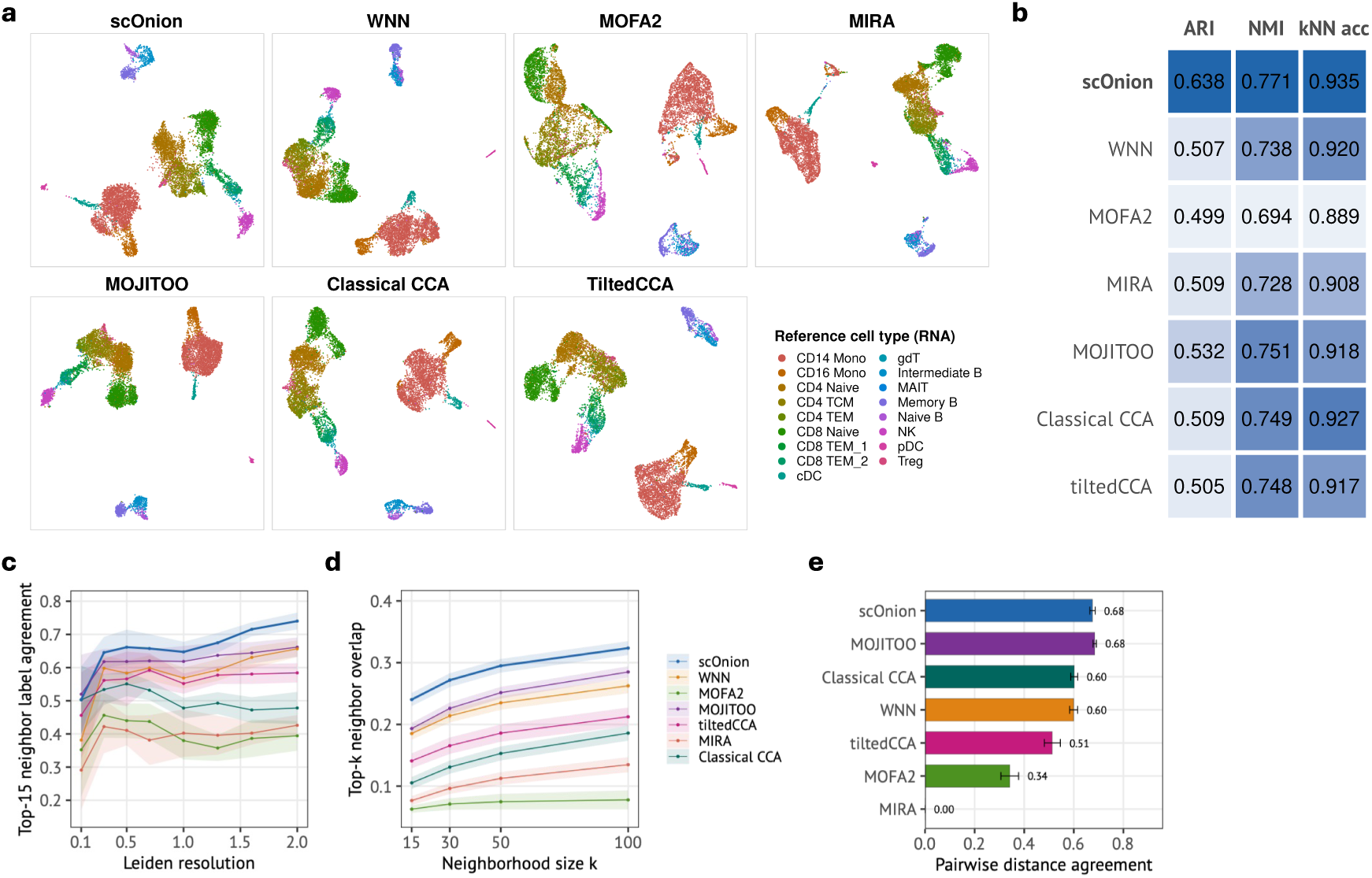
Real-data joint-embedding benchmarks, with and without cell-type an-notation. **a**, Joint-embedding UMAPs for seven integration methods (scOnion, WNN, MOFA2, MOJITOO, Classical CCA, Tilted-CCA and MIRA) on the granulocyte-sorted PBMC 10K multiome (9,611 cells), each colored by the 17 RNA-reference-transferred cell types (Methods). All methods recover the major immune populations (quantified in panel b). **b**, Real-data integration benchmark on the same dataset (17 reference cell types). Adjusted Rand index (ARI), normalized mutual information (NMI) and *k*-nearest-neighbor label-transfer accuracy (*k* = 15) for the joint embeddings of the seven methods, evaluated against the Azimuth level-2 cell types transferred by RNA (Methods). scOnion recovers the reference cell types best on all three metrics (ARI 0.638 against at most 0.53 for the other methods). **c–e**, Annotation-free comparison on the NeurIPS 2021 bone-marrow multiome (24,997 cells), using no cell-type labels. Each modality’s own structure sup-plies the reference: RNA principal components and ATAC latent semantic indexing components are Leiden-clustered separately, and every embedding is scored against both references. Scores are rescaled per modality to the range between the cross-modal floor and the unimodal ceiling, then combined as their geometric mean, so an embedding that discards one modality scores zero (Methods). Bands and error bars are 95% confidence intervals across the 13 batches. Seven methods are shown. **c**, Agreement between a cell’s 15 nearest neighbors in the embedding and its own reference cluster, across Leiden resolutions 0.1 to 2.0. scOnion is highest at every resolution, and its advantage widens as the reference becomes finer. **d**, Fraction of a cell’s *k* nearest neighbors in the embedding that are also its *k* nearest neighbors in that modality’s own space, for *k* = 15 to 100. This measure uses no clustering on either side. scOnion is highest at every *k*. **e**, Spearman agreement between all pairwise cell-cell distances in the embedding and in each modality’s space, the one measure that reflects global geometry beyond local neighborhoods. scOnion and MOJI-TOO are indistinguishable (both 0.68). MIRA reaches the cross-modal floor and therefore scores zero. Source data are provided with this paper.

**Extended Data Figure 4:**
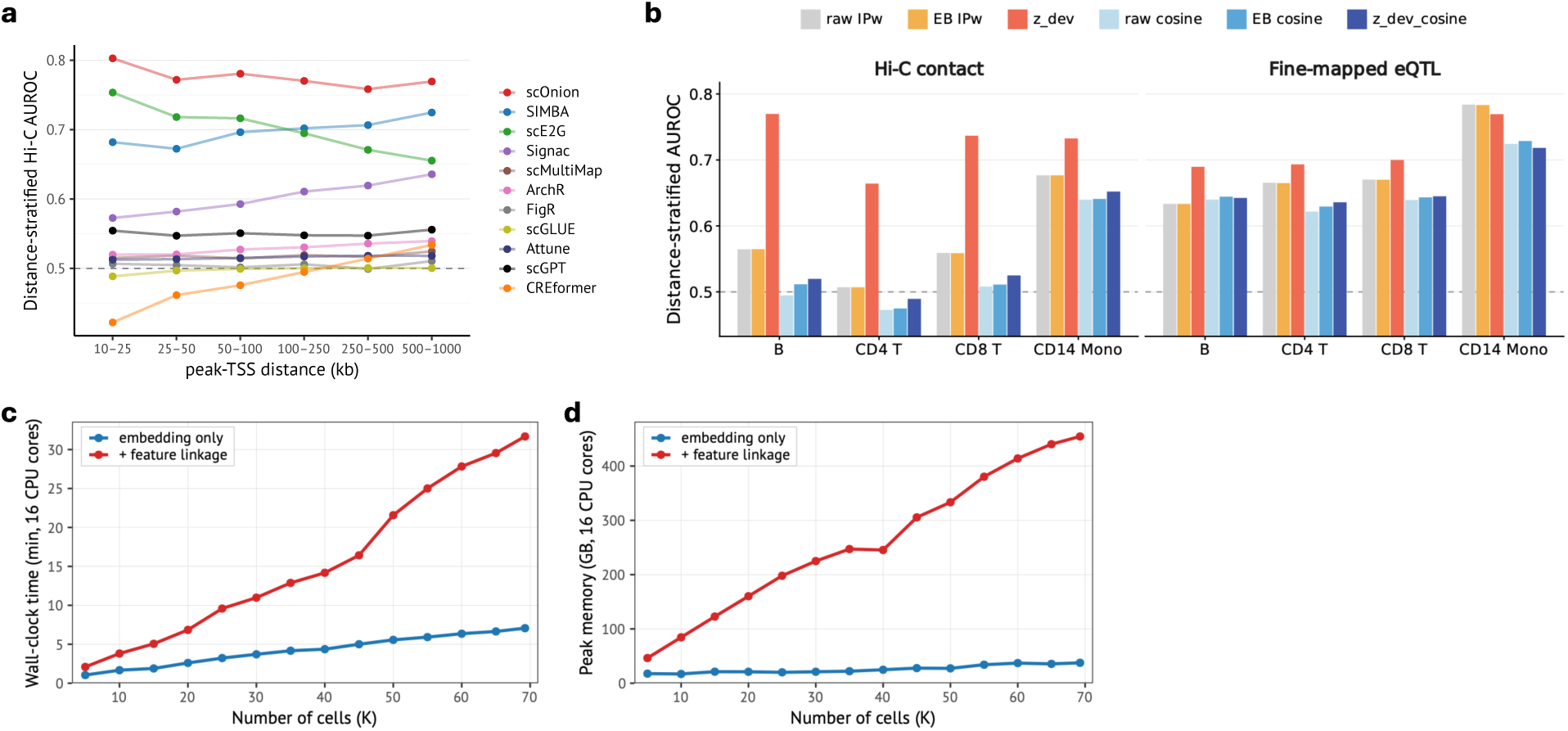
Distance-resolved linkage validation, linkage-score ablation and scaling to 69,249 cells. **a**, Distance-stratified Hi-C AUROC across six peak-to-TSS distance bins spanning 10 kb to 1 Mb (10–25, 25–50, 50–100, 100–250, 250–500 and 500–1000 kb) for eleven methods, including the scGPT and CREformer foundation models. scOnion maintains high AUROC across bins, including the distal ranges where windowed methods decline, whereas the foundation-model embeddings remain near chance throughout. **b**, Ablation of the gene–peak linkage score, showing distance-stratified AUROC against Hi-C contact (*n* = 39,970, 19,087, 42,547 and 16,760 positive pairs in B, CD4 T, CD8 T and CD14 monocytes) and fine-mapped eQTL (*n* = 198, 1,016, 418 and 74 positives) for six score variants, three adjustments (raw, empirical-Bayes shrinkage and *z*_dev_) applied to two bases (the weighted inner product and its distance-weighted cosine normalization), across four cell types (Methods). *z*_dev_ on the magnitude-carrying inner product is highest in every cell type against Hi-C, exceeding the unadjusted inner product by 0.06 to 0.21 AUROC, and is highest in three of four cell types against eQTL. Cosine variants, which divide out the magnitude, are lower on average on both truth sets. Empirical-Bayes shrinkage reproduces the unadjusted score, since it reweights monotonically in the standard error and rarely reorders pairs within a stratum. **c**, scOnion wall-clock time against number of cells on the NeurIPS 2021 bone-marrow multiome (69,249 cells, 13,431 genes and 116,490 peaks), from 5,000 cells to the full dataset on 16 CPU cores. Curves show the joint cell embedding alone and the embedding plus the *cis* gene–peak linkage table (every peak within ±1 Mb of a TSS, including the per-link jackknife standard error). The full dataset takes 7.1 minutes to embed and 31.7 minutes end to end (Methods). Shared preprocessing and object load and save, a flat 2.5 to 3.2 minutes per job, are excluded. **d**, Peak memory for the same runs, rising from 46 GB at 5,000 cells to 455 GB at 69,249 cells. Memory is the whole-job peak reported by the scheduler and includes the parallel worker processes of the linkage stage. For the end-to-end series the larger of the two stage values is taken. It grows more slowly than cell number. Source data are provided with this paper.

**Extended Data Figure 5:**
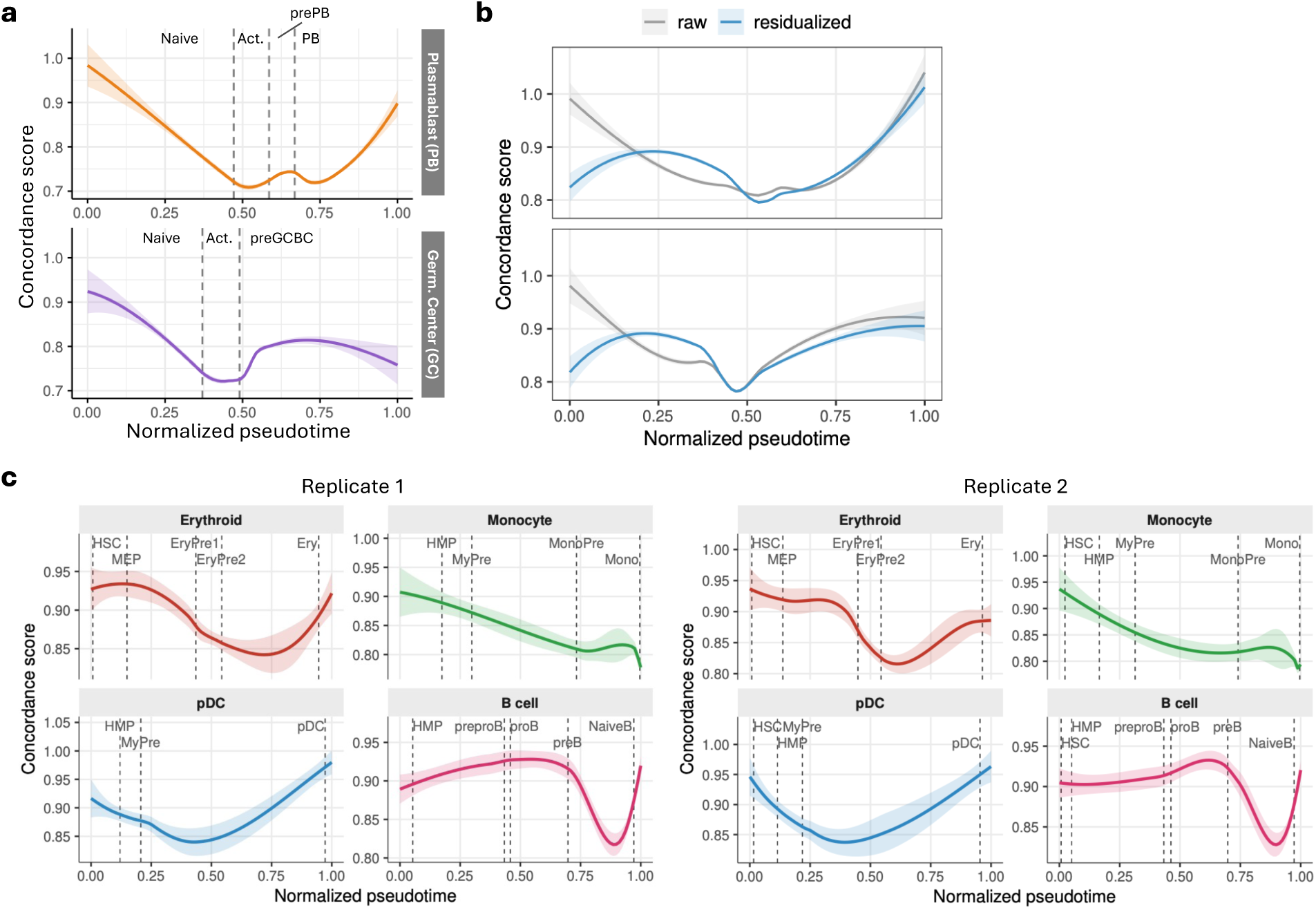
Concordance dynamics along differentiation reproduce in a second donor, persist after adjustment for technical and cell-state covariates, and generalize to an independent system. **a**, Concordance along normalized pseudotime for the plasmablast (PB) and germinal-center (GC) lineages in donor 1, shown as a loess fit with a 95% confidence band. Both lineages show a transient concordance decrease near fate commitment, reproducing the pattern in the primary donor (Fig. 5b). **b**, Cross-modal concordance along normalized pseudotime for the plasmablast (PB, top) and germinal-center (GC, bottom) lineages of the primary donor, shown before (raw) and after residualizing per-cell concordance on technical and cell-state covariates (the log_10_ per-cell sum of log-normalized RNA expression as a depth covariate, an immediate-early-gene stress score and S-phase and G2M scores, which jointly explain *R*^2^ = 0.126; Methods). The commitment dip near mid-pseudotime persists after residualization (cells split into ten equal-sized groups along pseudotime; two-sided Wilcoxon rank-sum test comparing the group with the lowest mean concordance with the last group, *P* ≤ 1.8 × 10^−100^ in both lineages). **c**, scOnion applied independently to the human bone-marrow CD34+ progenitor multiome of Otto et al. [28]. Concordance along normalized Palantir pseudotime for four differentiation lineages (erythroid, monocyte, pDC and B cell), shown as loess fits with 95% confidence bands and dashed lines marking the median pseudotime of each cell state, for two technical replicates. Concordance is modulated along pseudotime in a lineage-specific pattern, with no single uniform dip, reproducibly across replicates (Methods). Statistical tests and sample sizes are reported in Methods. Source data are provided with this paper.

**Extended Data Figure 6:**
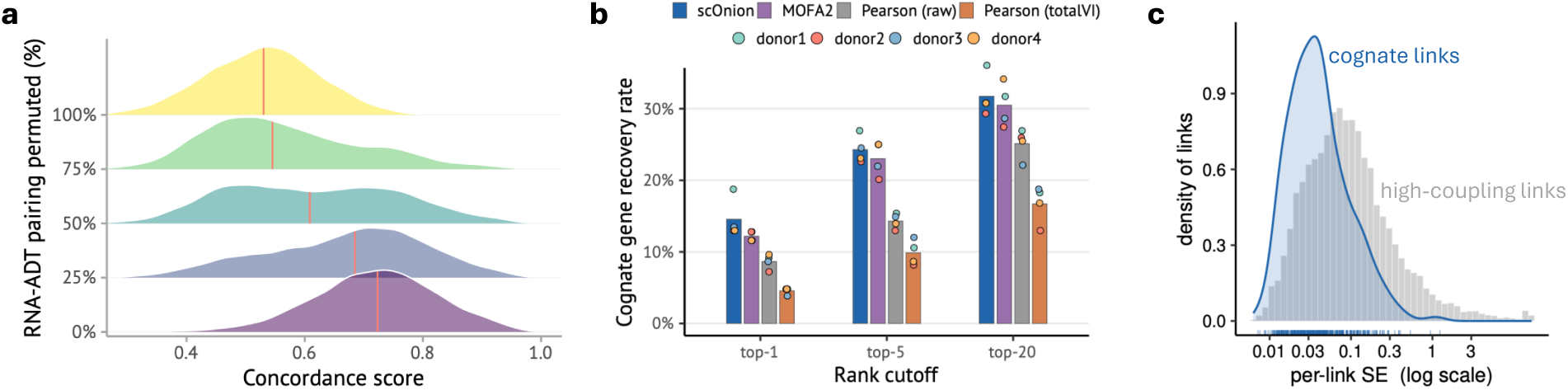
CITE-seq controls and competitive cognate-gene recovery. **a**, Concordance score distributions as an increasing fraction of RNA–ADT cell pairings is permuted (0% to 100% in 25% increments, each level an independent draw in which the selected cells’ ADT pro-files are permuted among themselves, so intermediate levels are bimodal), for a representative Hao PBMC day-0 donor, the CITE-seq analogue of the scMultiome permutation control (Fig. 2b and Methods). As the pairing is destroyed the distribution shifts from the intact median (0.72 to 0.75 across donors) toward a pairing-destroyed floor (median 0.53), consistently across all four donors. **b**, Cognate-gene recovery rate at top-1, top-5, and top-20 rank cutoffs for scOnion (uncertainty-aware score cos −2 SE), MOFA2, Pearson correlation on raw counts, and Pearson correlation on totalVI-denoised values, across four donors. scOnion recovers the most cognate genes at all three cutoffs, significantly more than either Pearson baseline at every cutoff (McNemar *P* ≤ 8.6 × 10^−5^) and more than MOFA2 at top-1 (*P* = 0.019), with the top-5 and top-20 margins over MOFA2 within sampling variation (*P* = 0.11 and 0.55). MOFA2 is next, above both Pearson baselines, and the totalVI-denoised correlation is lowest. scOnion additionally supplies a closed-form per-link standard error, which MOFA2 does not provide. **c**, Distribution of per-link standard error for cognate links and for the high-coupling links that top the raw score, across all proteins. Cognate links carry systematically lower standard error, roughly 3.5-fold (paired one-sided Wilcoxon test over 725 protein–donor cases, *P* = 5.7 × 10^−89^). Panel c overlays a box plot on each violin; the center line is the median, the box limits are the first and third quartiles (interquartile range, IQR), and the whiskers extend to the most extreme value within 1.5× the IQR. Outliers beyond the whiskers are plotted as individual points. Statistical tests and sample sizes are reported in Methods. Source data are provided with this paper.

## 6 Methods

### 6.1 The scOnion model

A single fit returns three quantities from one closed-form decomposition (Fig. 1a). The first is a per-cell cross-modality concordance score (Fig. 2), the cosine similarity cossim 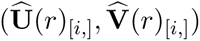 of a cell’s paired coordinates, rescaled to [0, 1] and defined in Per-cell concordance score. The second is a joint cell embedding **W** (Fig. 3). The third is a joint feature embedding, which carries the gene–peak linkage score (the rE2G coupling plotted in Fig. 5e,f) with its distance-aware z-statistic and per-link standard error, and its protein–gene form (Fig. 6).

#### Notation

**X** ∈ ℝ*^n^*^×^*^g^* and **Y** ∈ ℝ*^n^*^×^*^p^* are the normalized data matrices of the two paired modalities, with *n* cells, *g* genes and *p* features of the second modality (Construction of the input matrices from counts). 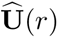 and 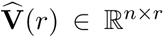 are the paired cell embeddings at rank *r*, 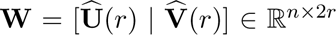 is the joint cell embedding, and 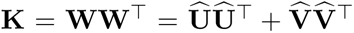 is the cell kernel of the joint embedding. *δ* is the genomic distance from a peak to a transcription start site (TSS). SE(*g, p*) is the per-link jackknife standard error, *s*_(−_*_i_*_)_ the linkage score of a feature pair with cell *i* omitted, *s̅* the mean of those *n* values, and *ω* the empirical-Bayes shrinkage weight.

#### Optimization objective

**XX**^T^ and **YY**^T^ are the cell-by-cell Gram matrices of the two modalities. The method applies to any two paired assays with matched cells and possibly distinct feature sets. scOnion first denoises each modality by truncating its singular values at a data-driven ratio threshold (3). For a fixed integer *r*, 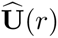 and 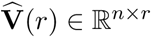 are the optimizers, over the set **O**(*n, r*) of *n* × *r* orthonormal matrices, of

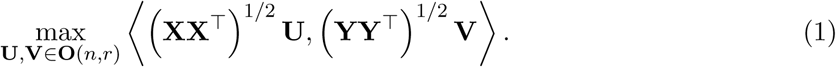

This can be read as joint principal component analysis in cell space, because the cell-space form 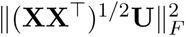 and the feature-space form 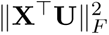 share their maximizers (Proposition S1).

#### Closed-form solution and rotation invariance

Let **UΣV**^T^ be the singular value decomposition (SVD) of (**XX**^T^)^1*/*2^(**YY**^T^)^1*/*2^. The first *r* columns of **U** and **V** solve 1 (Theorem S2), uniquely up to a joint rotation. The per-cell similarity, the pairwise cell distances in **W** that define neighborgraphs and UMAPs, and the cross-modal loading product are each invariant to that rotation (proofs in Supplementary Note 1).

#### Feature embedding and projection

**W** serves as a common low-dimensional representation for the features of both modalities. The projections

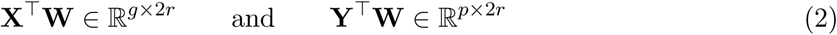

are the inferred feature loadings for genes and peaks. Diagonalizing the cross-modal loading product **X**^T^**WW**^T^**Y** by SVD gives the shared component loadings **P** (genes) and **Q** (peaks, or proteins in the CITE-seq setting) with singular-value weights *d*, which define the linkage score of Joint feature embedding and gene–peak linkage score. The number of retained feature components *r*^′^ can differ from *r* and is selected by the same ratio rule 3 at its own threshold *T*_3_.

#### Rank selection

The rank is chosen from the data using a consecutive singular value ratio thresholding. For each modality we take the SVD of its scaled matrix with singular values *σ*_1_ ≥ · · · ≥ *σ_n_* and retain, at a threshold *T*,

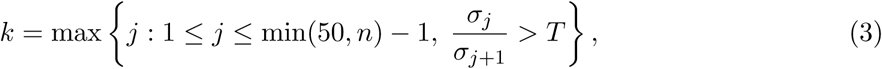

so components are dropped once the spectrum flattens (Supplementary Fig. 1). The two selected ranks *r*_1_ and *r*_2_ may be collapsed to their common minimum before the joint decomposition (`reconcile.ranks = TRUE`) or each modality may keep its own rank for modality-specific down-stream analysis. The joint cell embedding **W** has min(*r*_1_*, r*_2_) paired components either way. The defaults *T*_1_ = *T*_2_ = 1.01 and, for the feature-embedding rank *r*^′^, *T*_3_ = 1.1 were used for every real dataset. The selected *r* and *r*^′^ and the rank convention for each analysis are listed in Supplementary Table 3.

#### Implementation

scOnion is implemented in R. It denoises each modality by a truncated SVD of up to 50 components, using `irlba::irlba` on the dense scaled expression matrix and `RSpectra:: svds` [36] on the sparse term-frequency-inverse-document-frequency accessibility matrix, and multiplies the rank-truncated operators with a custom Rcpp/Eigen routine. A complete fit is one call to `Onion()`, the entry point for every result except the simulation grid (Joint cell embedding benchmark), and gene–peak linkage tables come from `OnionLinkageTable()`. Versions of every dependency and tool are listed in Supplementary Table 2.

### 6.2 Ethics statement and data processing

#### Ethics statement

All datasets are previously published and publicly available, generated under their original studies’ ethical approvals and participant consent. No new human or animal data were collected, and all analyses used de-identified secondary data.

#### 10x PBMC multiome

We imported Cell Ranger ARC barcode-filtered matrices for four 10x Genomics PBMC multiome samples, 3K and 10K, granulocyte-sorted and unsorted, from the 10x Datasets portal (https://www.10xgenomics.com/datasets) and retained cells with 1,500–20,000 RNA counts, 2,000–50,000 ATAC counts, nucleosome signal below 1.5 and TSS enrichment above 4, with no bound on the mitochondrial fraction. Cell types are Azimuth level-2 labels transferred from Hao et al. [8], via `pbmcMultiome.SeuratData` for the sorted 10K sample. The quality-controlled sorted 10K object (36,601 genes, 143,883 peaks) underlies Fig. 2 and Fig. 4b–e, and the per-cell measure comparison drops rare and low-confidence types (gamma-delta T, MAIT).

#### IGVF activated B-cell multiome

We obtained the raw FASTQ files and per-barcode anno-tation of the IGVF activated B-cell multiome [25], released on the IGVF Data Portal (https://data.igvf.org, Supplementary Table 1), through the consortium Synapse project (`syn52948700`) on 8 September 2025. Reads were processed with Cell Ranger ARC [37]. The annotation gives each of the 60,086 barcodes one of 18 cell states and a MULTI-seq sample (donor and collection day). We kept annotated barcodes with RNA and ATAC counts between the 2.5th and 97.5th percentiles, mitochondrial fraction and nucleosome signal below the 97.5th and TSS enrichment above the 2.5th, over the aggregated dataset. Donor 1 was profiled at days 0, 5 and 6, donor 2 at days 0 to 6.

#### NeurIPS 2021 bone-marrow multiome

We downloaded the NeurIPS 2021 bone-marrow multiome [38] from GEO (`GSE194122`, https://www.ncbi.nlm.nih.gov/geo/query/acc.cgi?acc=GSE194122) on 24 August 2026 and used its raw counts. A batch-stratified subsample spanning all 13 batches entered the annotation-free benchmark (Supplementary Table 1).

#### Bone-marrow CD34+ progenitor multiome

The T-cell-depleted CD34+ multiome of Otto et al. [28] was downloaded from GEO (https://www.ncbi.nlm.nih.gov/geo/, accessions in Supplementary Table 1) on 7 April 2026 as two technical replicates and fit independently, keeping annotated barcodes with at least 200 detected genes and peaks on the standard chromosomes. Cell states, Palantir [39] pseudotime and terminal lineages were taken as published.

#### SEA-AD middle temporal gyrus multiome

We downloaded the per-donor final-nuclei objects of two SEA-AD donors [17] from its registry on Amazon S3 (https://sea-ad-single-cell-profiling.s3.amazonaws.com) on 24 February 2026, kept 10xMulti nuclei with at least 3,000 ATAC features and more than 12,500 ATAC counts and used the Subclass and Class annotations.

#### Hao PBMC CITE-seq

Raw RNA counts of the Hao et al. PBMC CITE-seq dataset [8] came from GEO `GSE164378` (sample `GSM5008737`, https://www.ncbi.nlm.nih.gov/geo/query/acc.cgi?acc=GSE164378), and counts of its 228-antibody TotalSeq-A panel (ADT), metadata and level-2 labels from the study’s reference object (Zenodo record 7779017). We used the four donors profiled in one batch and demultiplexed by cell hashing [40], at pre-vaccination day 0, as biological replicates.

#### Orthogonal validation resources

Gene–peak links were validated against cell-type-matched ENCODE4 intact Hi-C loop calls [21, 22] (HiCCUPS [41, 42], https://www.encodeproject.org, downloaded 29 May 2026), OneK1K fine-mapped eQTLs [23] from the eQTL Catalogue (https://ftp.ebi.ac.uk/pub/databases/spot/eQTL/susie/QTS000038/, downloaded 10 June 2026) and JASPAR2024 motifs [43] (https://jaspar.elixir.no, TFBSTools [44], BSgenome.Hsapiens.UCSC.hg38 [45]), with accessions in Supplementary Table 1.

#### Construction of the input matrices from counts

RNA counts were log-normalized (`LogNormalize`, scale factor 10^6^) and centered and scaled over all genes to form **X**. ATAC counts were normalized by TF-IDF (Signac `RunTFIDF` [11]) to form **Y**. For CITE-seq, ADT counts were CLR-normalized (`NormalizeData`, `method="CLR"`, `margin=2`) to form **Y**. Inside the fit scOnion drops constant features and retains genes whose column standard deviation in **X** exceeds `x.sds`= 0.95.

### 6.3 Per-cell concordance score

#### Definition

The columns of **Û**(*r*) and **V̂**(*r*) are paired one to one by the SVD, so a cell’s coordinates in the two embeddings are directly comparable. For cell *i* we compute the cosine similarity of its paired coordinates,

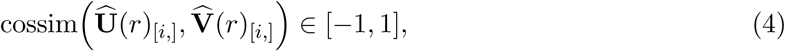

and report it rescaled to the unit interval,

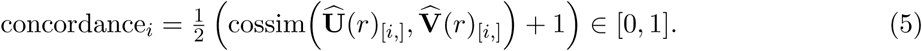

Values near 1 mark a cell whose local structure is supported similarly by both modalities (Fig. 2a). We compute the score with `scOnion::OnionConcordance`.

For per-cell distributions and cell-type contrasts the unit of replication is the cell and the statistics are descriptive of one sample, so every contrast was repeated in independent samples and donors. For the linkage benchmarks the unit is the gene–peak or protein–gene pair. Biological replicates are independent donors, and the two bone-marrow CD34+ datasets are technical replicates (Supplementary Table 1).

#### Graded-divergence simulation for concordance

We generated 3,024 cells in six balanced groups of 504, with group means drawn in random directions of norm 10 in a 23-dimensional latent space **Z** and isotropic within-group scatter of standard deviation 1. A second, independent set of group means with independently drawn labels gave a latent **Z**^⊥^, orthogonalized against **Z** within each cell and rescaled to matching norm. Each cell *i* was assigned an angle *θ_i_* at one of nine levels from 5^◦^ to 45^◦^ in steps of 5^◦^, drawn on a stratified uniform grid of width ±1^◦^ about each level, with 56 cells at every combination of group and level so that every group has the same true mean concordance. The first modality received **Z** and the second 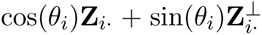, so the angle between a cell’s two representations is exactly *θ_i_*and its true concordance is (1 + cos *θ_i_*)*/*2 by 5. Latent factors were projected into 1,000 gene and 5,000 peak features by orthonormalized loading rows, with no dropout, clamping or additive noise. scOnion was fit with the rank fixed at the latent dimension and the internal feature filter disabled. Over ten simulations, estimated and true per-cell concordance were compared by two-sided Spearman correlation, overall and within each group (Extended Data Fig. 1c).

#### Synthetic cell mismatching as null

The 10,489 quality-control-filtered cells of the unsorted PBMC 10K sample were partitioned into ten folds stratified by Azimuth level-2 cell type, and cell identities within the RNA modality were shuffled one fold at a time, in 10% increments from 0% to 100%, each level nesting the cells already shuffled, and concordance was recomputed at every level (Fig. 2b). In each of the four PBMC samples, concordance alone was used to classify scrambled against intact cells at each scrambled fraction from 10% to 90%, scored by AUROC (Extended Data Fig. 1a).

#### Association with quality control and neighborhood agreement

Concordance was com-pared with six per-cell quality-control metrics (Fig. 2e) by two-sided Spearman correlation on the granulocyte-sorted PBMC 10K sample (*n* = 10,280). Cross-modality neighborhood agreement was scored from a 25-nearest-neighbor graph on RNA (PC=1:30) and one on ATAC LSI (PC=2:30), by the normalized-rank RMSE between the two neighbor rankings (Fig. 2f) and by the Jaccard overlap of the two neighbor sets (Supplementary Fig. 2c), with parameters in Supplementary Table 3. A third measure was each cell’s Spearman correlation between its RNA profile and its gene-activity profile from `Signac::GeneActivity` [11] over the genes present in both (settings in Supplementary Table 2). Each measure was correlated with concordance on the same cells by two-sided Spearman correlation.

#### Cell-type contrasts and sensitivity

Cell-type and cell-state contrasts (e.g. CD14 vs CD16) were tested by one-sided Wilcoxon rank-sum test on per-cell concordance in the sorted and unsorted PBMC 10K samples (Fig. 2d) and in the two 3K samples (Supplementary Fig. 2b). To control sequencing depth, every cell of each modality in both PBMC 10K samples was thinned by binomial subsampling of its raw counts to a common target, the tenth percentile of observed depth, equivalent to `DropletUtils::downsampleMatrix` [46], and the thinned matrices were re-preprocessed. The depth-matched concordance was correlated with each cell’s original `nCount_ATAC` and the Fig. 2d contrasts were re-tested (Extended Data Fig. 1b). Sensitivity to the denoising threshold was assessed on the sorted PBMC 10K sample by refitting across the usable range *T*_1_ = *T*_2_ ∈ [1.0025, 1.02] with *r* auto-selected and comparing per-cell concordance and the Fig. 2d contrasts with the default fit (Supplementary Fig. 2d and Supplementary Table 3). Concordance was also compared with the Tilted-CCA synchrony score and the WNN modality weight within six immune lineages (Extended Data Fig. 1d,e and Supplementary Note 6). The test, *n* and exact *P* value behind every panel are in Supplementary Table 4.

### 6.4 Joint cell embedding

#### Definition

Solving 1 gives two unscaled *r*-dimensional cell embeddings, and the joint cell embedding is their concatenation,

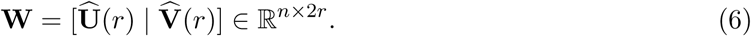

Using unscaled principal directions lets every retained direction contribute equally.

#### Initial coarse cell-type label assignment

Labels for the sorted PBMC 10K sample start from the Azimuth level-2 annotation in `pbmcMultiome.SeuratData`. Cells with closely related level-2 types were grouped under coarse labels (Supplementary Table 3) as initial assignment. Two joint subclusters whose markers contradicted their transferred lineage were removed before their compartments were scored, T cells among the CD14 monocytes (220 of 3,028 cells) and effector cells among the naive CD8 T cells (17 of 1,485).

#### Clustering and subtyping

**W** was imported into Seurat object as a dimensional reduction. A shared-nearest-neighbor graph was built over all 2*r* dimensions with `FindNeighbors`, clusters were called with `FindClusters`, fine subtypes were resolved by iterative `FindSubCluster` on the same graph, and display coordinates came from `RunUMAP` on **W** [47] (settings in Supplementary Table 3).

#### Within-modality separation of joint subtypes

A compartment is the set of cells carrying both a given joint cluster and the matching transferred label (Fig. 3f and Supplementary Fig. 3d). Seven compartments holding 21 subtypes over 6,835 cells were scored, and the naive CD8 T subclusters and the NK cluster were not. Within each compartment a silhouette width was computed per cell with `cluster::silhouette`, using the scOnion subcluster as the grouping and Euclidean distance in one modality’s own reduction, RNA principal components 1–25 or ATAC LSI components 2–30, and a subtype’s within-modality silhouette is the mean over its cells (Fig. 3f and Supplementary Fig. 3a,b). For Fig. 3f the joint embedding and its UMAP were recomputed with the released package at the default settings (`reconcile.ranks = TRUE`, *r* = 29) on the same 10,280 cells, keeping the subtype labels of the original clustering, which the recomputed embedding recovers (adjusted Rand index 0.83 against the joint clusters; the two compartments shown remain 99.4% and 98.3% intact). The RNA and ATAC silhouettes do not depend on the joint fit. Supplementary Fig. 3a relates the mean of a subtype’s two silhouettes to its mean concordance by two-sided Spearman correlation over the 21 subtypes (Supplementary Table 4).

### 6.5 Joint cell embedding benchmark

All compared methods, here and in the later benchmarks, were run on the same paired matrices as scOnion.

#### Compared methods

Seurat weighted nearest neighbor (WNN, `FindMultiModalNeighbors`) [8] embeds a weighted multimodal neighbor graph built from an RNA principal-component and an ATAC LSI reduction. MOFA2 [9] was fit separately on each dataset, with the factor number set for each. MOJITOO [7] integrates the leading singular vectors of each modality by canonical correlation. MIRA [10] fits a topic model per modality at default settings, and its joint topic representation was embedded by UMAP. Tilted-CCA [6] was run with its standard pipeline of `create_multiSVD`, and its compared embedding is the common score. Classical canonical correlation (CCA) analysis [48] took the singular vectors of the cross-modality inner product of the two modality bases at a common rank.

#### Construction of the simulation grid

For Fig. 3b and Extended Data Fig. 2a,b, two latent coordinates drawn at ±1 give *n* = 200 cells in four ground-truth groups, one per corner of the lineage × maturity plane. Each modality was assigned one factor as its dominant axis, lineage for RNA and maturity for ATAC, and its second axis was a convex combination of the complementary factor and independent noise, mixed by a signal ratio varied over five levels from 0 to 1. Crossing the RNA and ATAC levels gave a 5 × 5 grid of 25 paired datasets. Each latent was projected into 1000 genes and 5000 peaks by loadings carrying designated marker features against a weak Gaussian background, with second-axis loadings Gram–Schmidt orthonormalized against the first. Measurement sparsity was emulated by modality-specific Bernoulli dropout, higher for ATAC, with negative values set to zero. Numeric parameters are in Supplementary Table 3.

#### Comparison on the simulation grid

scOnion and the six methods were run on all 25 datasets. On this grid we solved 1 directly, with the grid-specific scaling and rank rules of Supplementary Table 3. Recovery was scored per condition on each method’s two-dimensional UMAP embedding (settings in Supplementary Table 2) by standardizing the coordinates, running *k*-means with *k* = 4 (25 restarts, at most 50 iterations) and comparing the partition with the four ground-truth groups by adjusted Rand index (ARI, `mclust::adjustedRandIndex`), averaged over ten *k*-means seeds. Condition-averaged ARI was compared between methods by two-sided paired Wilcoxon signed-rank test over the 25 conditions, and the average silhouette width of the four ground-truth groups (`cluster::silhouette`) was computed in the same embeddings (Supplementary Table 4).

#### Real PBMC data against transferred labels

For Extended Data Fig. 3a,b, the comparison was repeated on the granulocyte-sorted PBMC 10K multiome against the Azimuth level-2 taxonomy transferred by RNA alone (`pbmcMultiome.SeuratData`, Supplementary Table 1). For each of the seven methods, a *k*-nearest-neighbor graph on the method’s embedding (`FNN::get.knn`) was Leiden-clustered (`igraph::cluster_leiden`) and scored against the reference labels by ARI and normalized mutual information, and *k*-nearest-neighbor label-transfer accuracy was computed by leave-one-out plurality vote (*k* = 15).

#### Annotation-free benchmark

For Extended Data Fig. 3c–e, each modality’s own structure supplied the reference on a batch-stratified subsample of the NeurIPS bone-marrow multiome [38] (GEO `GSE194122`, Supplementary Table 1). RNA principal components and ATAC LSI compo-nents were each Leiden-clustered with the same functions at a range of resolutions (Supplementary Table 3). Each embedding was scored against the RNA-derived and the ATAC-derived reference by three measures. *k*-nearest-neighbor label transfer (*k* = 15) predicts each cell’s reference cluster by plurality vote and is corrected as 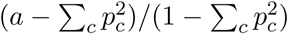, where *a* is the accuracy and *p_c_* the proportion of cells in reference cluster *c*. Neighborhood overlap is the fraction of a cell’s *k* nearest neighbors in the embedding that are also among its *k* nearest neighbors in the modality’s own space, at four values of *k* (Supplementary Table 3). Pairwise distance agreement is the Spearman correlation between all cell–cell distances in the embedding and in the modality’s space, computed within batch on up to 2,000 cells. To combine modalities, each score was rescaled to the range attainable on that modality, (*s* − floor)*/*(ceiling − floor), with the ceiling the score of that modality’s own unimodal embedding and the floor the score of the opposite unimodal embedding, clipped to [0, 1] and combined as the geometric mean, so an embedding that discards one modality scores zero. All quantities were computed per batch, with 95% confidence intervals across batches.

### 6.6 Joint feature embedding and gene–peak linkage score

In this study, for scMultiome data, **P**, **Q** and *d* are computed per-chromosome, so every gene–peak linkage score is a within-chromosome (*cis*) score. But one may generate **P**, **Q** without such restriction. A candidate peak is also called a candidate regulatory element (RE), and *δ* is measured from it to the strand-aware TSS of the gene it is scored against. Feature maps are built for the autosomes and chromosome X. Chromosome Y is absent in a subset of donors and is not scored.

#### Raw gene–peak linkage score

For gene *g* and candidate peak *p* the raw linkage score is the singular-value-weighted inner product

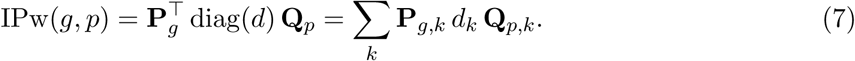

Because **X**^T^**WW**^T^**Y** = **P** diag(*d*)**Q**^T^, this equals the embedding-space inner product (**X**^T^**W**)*_g_* · (**Y**^T^**W**)*_p_*. `OnionLinkageTable` scores every *cis* pair whose peak midpoint lies within ±1 Mb of the TSS.

#### Population-specific feature embedding

Restricting the rows of **X**, **Y** and **W** to a population *S* gives 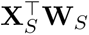 and 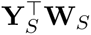, which accumulate only over the cells in *S*, and passing them through 7 yields population-specific linkage scores from a single global fit without resolving 1.

#### Distance stratified linkage score and the z-statistic

Raw linkage scores decay with genomic distance. *Cis* pairs were binned by *δ* at fixed breakpoints (Supplementary Table 3), and pairs with *δ <* 10 kb were excluded from every distance-stratified analysais because the loop caller has an effective floor near that distance. Each score was standardized within its bin by the bin mean *µ*_0_[bin(*δ*)],

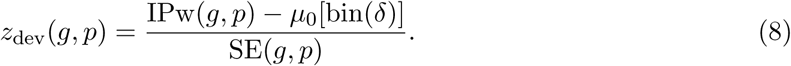

#### Per-link jackknife standard error

SE(*g, p*) is the leave-one-cell-out jackknife standard error of the raw score at the fixed embedding, computed in closed form without refitting. The score is bilinear in the cell embedding, so all *n* omitted-cell scores follow from two matrix-vector products (Supplementary Note 3).

#### Alternative linkage adjustment via Empirical-Bayes shrinkage

The raw score can be shrunk toward zero by its uncertainty as *ω* ·IPw with *ω* = *τ* ^2^*/*(*τ* ^2^ +SE^2^) and *τ* ^2^ = max(0, var(IPw)−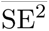).

#### Motif enrichment

For each of five PBMC populations (CD14 monocytes, CD14 plus CD16 monocytes, B, CD4 T and CD8 T cells) we took the top 25 marker genes per constituent cluster from Seurat `FindAllMarkers` ordered by average log-fold change, after removing long non-coding, microRNA, antisense and clone-identifier names, selected each marker’s top 50 scOnion-linked peaks by *z*_dev_ (8) and pooled them within the population (Fig. 4d and Supplementary Fig. 4c, counts in Supplementary Table 4). Peaks were ranked on the population-specific feature embed-ding of the whole-PBMC fit used in Fig. 4b,c, with no concordance-based cell filtering. En-richment was computed with Signac `FindMotifs` on JASPAR2024 [43] matrices assembled with `TFBSTools` [44] on `BSgenome.Hsapiens.UCSC.hg38` [45] sequences, against the default background that `MatchRegionStats` samples from the whole-PBMC peak universe to match the query GC content, with query peaks excluded from the pool (Supplementary Table 2). Motifs at adjusted *P <* 0.05 were collapsed into transcription-factor families by a curated mapping.

#### Co-activity

Each gene’s *cis* candidates were ranked by raw IPw, and for five neighborhood sizes (Supplementary Table 3) the mean absolute Spearman correlation across cells between the gene’s expression and each peak’s accessibility was compared across the scOnion ranking, the genomic-distance ranking and a random ordering of the same candidates (Supplementary Fig. 4b).

### 6.7 Gene–peak linkage benchmark against Hi-C and eQTL

#### Design

We evaluated *cis* gene–peak rankings against cell-type-matched ENCODE intact Hi-C loops [21, 22] and OneK1K fine-mapped eQTLs [23] in B, CD4 T, CD8 T and CD14 monocyte cells. scOnion was scored by *z*_dev_ on the population-specific feature embedding of the whole-PBMC fit, with no concordance-based cell filtering. Eight published linkage methods, SIMBA, scE2G, Signac, scMultiMap, ArchR, FigR, scGLUE and Attune, were scored by their native linkage score joined to the identical ±1 Mb candidate grid, a distance baseline was ranked by negative distance, and the foundation models scGPT and CREformer were scored from their pretrained embeddings (versions and settings in Supplementary Table 2).

#### Compared methods

Signac `LinkPeaks()` [11] and FigR `runGenePeakcorr()` [12] were run with default parameters except for a ±5 Mb window and no significance cutoff, so that every pair could be ranked. SIMBA [13] was run as a PyTorch-BigGraph co-embedding of cells, genes and peaks on all features. scGLUE [14] was trained as a variational model with regulatory inference and Attune [49] as a cross-attention transformer. scMultiMap [50] and ArchR `addPeak2GeneLinks()` [51] were run at a 1 Mb maximum distance, and scE2G [24] as the ENCODE-rE2G pipeline in inference-only mode with the pretrained power-law model and no Hi-C.

#### Foundation-model embeddings

Two pretrained single-cell foundation models were scored on the same candidate pairs, assessed only on their exported embeddings. For scGPT [19] we used embeddings in which genes and peaks share one space, from a model fine-tuned on the PBMC 10K multiome per cell type, and scored each pair by the dot product of its gene and peak embeddings. For CREformer [20] we used its gene embeddings from the Regulatory encoder, contextualized over peaks within ±300 kb of the TSS, with peak embeddings from the Elementary encoder, both computed per cell type from pseudobulk accessibility, and the same dot product.

#### Positive sets

We downloaded the ENCODE4 intact Hi-C loop calls for the four cell types from the ENCODE portal (https://www.encodeproject.org, accessions in Supplementary Table 1) on 29 May 2026. We retained loops at donut-corrected FDR ≤ 0.01 with span between 10 kb and 1 Mb on chromosomes 1–22 and X, reduced anchors to unique bins and built an anchor-adjacency graph. A peak was Hi-C-supported for a gene if it overlapped a bin that is a loop partner of a bin overlapping that gene’s promoter, defined with GenomicRanges `promoters()` as 2000 bp upstream to 200 bp downstream of the TSS, excluding the gene’s own promoter bins. We downloaded the OneK1K SuSiE 95% credible sets from the eQTL Catalogue (https://ftp.ebi.ac.uk/pub/databases/spot/eQTL/susie/QTS000038/) on 10 June 2026. Lineage truths aggregated the matched OneK1K subtypes, with CD14 monocytes matched to `CD14_Mono` only, by the maximum posterior inclusion probability (PIP) per variant, and a pair was eQTL-supported if the peak contained a credible-set variant of that gene at PIP *>* 0.1. Positive counts per cell type are in Supplementary Table 1.

#### Distance-controlled AUROC

Within each stratum, a gene × distance-bin cell for Hi-C and, because eQTL positives are sparse, a cell type × distance-bin cell for eQTL, all candidate peaks were ranked by the method’s score, unscored peaks were assigned the lowest rank, and the Mann–Whitney statistic *U* was computed. The pooled AUROC is 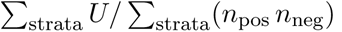 over strata containing both positives and negatives. Ranking performance throughout is this area under the receiver operating characteristic curve, computed from the Mann–Whitney statistic, threshold-free and without a *P* value. The linkage-score ablation of Extended Data Fig. 4b is described in Supplementary Note 6.

### 6.8 Runtime and scalability

#### Time to result across methods

For Fig. 4e we measured the wall-clock time for each of eleven methods to go from processed matrices to a *cis* gene–peak linkage table over every peak within ±1 Mb of a TSS. Shared preprocessing was performed once and excluded, and the matrices-to-links step was timed. The source object was the sorted PBMC 10K multiome with all cell types (36,601 genes and 143,883 peaks), with the feature set held fixed (Supplementary Table 3). Sizes up to 10,000 were random down-samples of the object, and 15,000 and 20,000 were built by tiling duplicated cells and fragments under unique barcodes. All methods were timed on the same set of matrices. CPU methods ran on 16 cores. GPU methods report single-GPU wall time, scGLUE and Attune on one NVIDIA A100, scGPT on an H200 and CREformer on an A100 MIG partition. Each method was timed on its production entry point (Supplementary Table 2), scOnion on `Onion()` followed by `OnionLinkageTable()`. scE2G’s time includes pipeline orchestration, scGPT was timed per cell type and extrapolated in proportion to cell number, and CREformer runs pretrained inference on pseudobulk profiles.

#### Joint cell embedding runtime against MIRA

scOnion and MIRA were timed on the same 10,333 cells of the granulocyte-sorted PBMC multiome. For scOnion the step from the processed matrices to the joint cell embedding is two truncated SVDs and a low-rank product, which run single-threaded, and it took a median of 4.5 minutes over three runs (242.6, 271.3 and 271.9 seconds) on one CPU core with no GPU. Timed end to end from raw counts under the same instrument, scOnion took a median of 6.0 minutes. MIRA’s complete native pipeline, from raw counts through normalization, feature selection, topic-number selection and topic-model fitting, took 2 hours 6 minutes (7,562 seconds) in one run on an NVIDIA A100 GPU with 8 CPU cores (Supplementary Table 2), about 21 times the scOnion end-to-end time.

#### Scaling sweep for scOnion

For Extended Data Fig. 4c,d we timed scOnion alone on the NeurIPS bone-marrow multiome at fourteen cell numbers (Supplementary Table 3), each an independent random sample of cells drawn without replacement and seeded by its target size. Normalization was performed once on the full object and excluded. The embedding stage and the linkage stage at ±1 Mb on the saved embedding that scores every candidate pair by 7 with its jackknife standard error were timed separately on 16 CPU cores with parallelism over chromosomes, with the rank selected inside each run by 3 at the default thresholds.

### 6.9 B-cell differentiation and fate bifurcation

#### Cell states and pseudotime

We fit the two donors independently at the default thresholds, donor 2 as the primary analysis and donor 1 as a replicate (ranks in Supplementary Table 3). Pseudotime came from Slingshot [52] (`slingshot::slingshot`) on each donor’s joint cell embedding, rooted at Naive with two lineages ending in PB and preGCBC (settings in Supplementary Table 2, lineage sizes in Supplementary Table 1), and was min-max normalized within lineage (Fig. 5a).

#### Concordance along pseudotime

Per-cell concordance was ordered along each lineage and summarized against normalized pseudotime by a loess fit (`geom_smooth` in ggplot2, span 0.75, degree 2) with 95% confidence bands, in both donors (Fig. 5b and Extended Data Fig. 5a). For the bone-marrow CD34+ progenitor multiome, the cells of each of its four lineages were ordered by its published Palantir [39] pseudotime (Supplementary Table 1), min-max normalized within lineage, and summarized by the same fit (Extended Data Fig. 5c). The residualization of Extended Data Fig. 5b and the covariate check of Supplementary Fig. 5c are described in Supplementary Note 6.

#### Fate axis

Concordance is an inner product of unit-normalized per-cell coordinates, so it de-composes additively across the *r* components. The commitment-window dip was attributed to components by contrasting the mean per-component contribution in the decision window (ActB and prePB) against resting and committed cells (Naive, PB and preGCBC), which localized it to component 2. The fate axis of Fig. 5c is component 2 of the feature embedding, whose gene loadings **P** and peak loadings **Q** were correlated (Pearson) with PB-versus-preGCBC differential expression and differential accessibility (Supplementary Table 4).

#### Differential gene expression and accessibility

Genes were called differentially expressed (DE) at |log2FC| ≥ 1, and differential accessibility was the difference in mean log-normalized ATAC signal per peak.

#### Gene-to-RE rewiring

Candidate REs were the peaks within 500 kb of the TSS in the donor 2 linkage table, keeping peaks accessible and genes expressed in both fates and requiring at least 10 candidates (filters in Supplementary Table 3). After equalizing the two fates in cell number, the feature projections were recomputed on each fate’s cells (Population-specific feature embedding), and every gene–RE pair was scored by 7 with its jackknife standard error and empirical-Bayes shrinkage *ω* (Fig. 5e,f). A gene’s rewiring score is one minus the Spearman correlation between its shrunken RE-linkage profiles in the two fates, over GENCODE v32 protein-coding genes [53], and its mean over the two donors was related to |log2FC| by two-sided Spearman correlation (Supplementary Fig. 5d and Supplementary Table 4).

#### Cross-donor reproducibility

The rewiring program was recomputed genome-wide in donor 1 with the identical algorithm, and per-gene rewiring was compared across donors by two-sided Spearman correlation (Fig. 5d). Recovery of the canonical B-cell fate regulators of Supplementary Table 6 among the most-rewired genes was scored in each donor by AUROC and by one-sided Wilcoxon rank-sum test of their rewiring scores against all other scored genes (Supplementary Table 4).

### 6.10 CITE-seq protein–gene coupling

#### Data and fit

The four day-0 donors of the Hao PBMC CITE-seq dataset [8] were each fit as a biological replicate. Input RNA matrix was constructed as stated previously, except that the log-normalized scale factor was 10^4^. The CLR-normalized ADT matrix entered unscaled as **Y**, and the variance filter was disabled (`x.sds`= 0). scOnion was fit per donor at the gene–peak default thresholds (Fig. 6a, ranks in Supplementary Table 3). Surface protein has no genomic coordinate, so links were scored from the global feature embeddings without distance stratified adjustment.

#### Cognate gene mapping

Each of the 228 antibodies was mapped to its cognate gene or genes by matching its name, clone suffix stripped, against the official, alias and previous symbols of the HGNC complete set (accessed 19 June 2026) [54]. Antibodies against complexes were split into gene sets, ambiguous cases were resolved by manual curation, and the isotype controls have no cognate (Supplementary Table 5). The ranking universe was GENCODE release 32 protein-coding, immunoglobulin and T-cell-receptor genes [53], intersected with each donor’s embedded genes (sizes in Supplementary Table 1). The 208 antibodies with a cognate in every donor’s embedding are scorable, and all reported rates use this intersection. For multi-gene cognates the rank is the minimum over the set.

#### Cognate retrieval and baselines

For every antibody, all genes were ranked by the cosine similarity between **P***_g_* and **Q***_p_*, and the recovery rate at *k* ∈ {1, 5, 20} is the fraction of the 208 scorable proteins whose cognate ranks within *k*, averaged over donors (Fig. 6d,e). The scOnion series is the uncertainty-aware score cos −2 SE_cos_ defined below. Baselines were run under the identical task, scoring and universe (Supplementary Table 2). Raw correlation is the Pearson correlation across cells between log-normalized expression and CLR-normalized abundance. totalVI [32] was trained on the raw RNA and ADT counts and scored by the same correlation on its denoised values. MOFA2 [9] was fit on the two normalized views and scored by the cosine similarity between its RNA and ADT factor loadings, and compared with scOnion at each *k* by McNemar paired test over protein–donor cases pooled across donors on a matched gene universe (Extended Data Fig. 6b and Supplementary Table 4). The permuted-pairing control (Extended Data Fig. 6a), the sparsity-stratified recovery analysis (Supplementary Fig. 6b) and the decoy-versus-cognate standard-error comparison (Extended Data Fig. 6c) are described in Supplementary Note 6.

#### Uncertainty-aware ranking

The closed-form jackknife of Per-link jackknife standard error carries over, giving a standard error for every gene–protein pair without refitting, mapped from the inner-product to the cosine scale (Supplementary Table 3). The uncertainty-aware score is cos −2 SE_cos_ (Fig. 6f).

#### Protein-anchored gene programs

The genes present in all four donors were ranked by cosine linkage to each of six proteins, averaging over donors, and each protein’s top 50 genes were tested for over-representation with Enrichr [55], with the same genes as background and Benjamini–Hochberg-adjusted *P* values (Fig. 6g, libraries in Supplementary Table 2). The enrichment was repeated on each donor’s own top 50.

### 6.11 Reporting summary

Further information on research design is available in the Nature Portfolio Reporting Summary linked to this article. All analyses are secondary analyses of published data. No statistical method was used to predetermine sample size. Data were excluded only by the cell-level quality-control filters, the 10 kb floor on gene–peak distance and the analysis-specific filters listed in Supplementary Table 3. There was no group allocation, so no randomization was performed, and investigators were not blinded to the labels assigned by the original studies. The statistical test behind each panel, its sidedness, *n* and exact *P* value are listed in Supplementary Table 4, and every random seed in Supplementary Table 3.

## Data availability

The four 10x Genomics PBMC Single Cell Multiome ATAC + Gene Expression datasets used here are available from the 10x Genomics Datasets portal, each from a single healthy donor and processed with Cell Ranger ARC: “PBMC from a Healthy Donor - Granulocytes Removed Through Cell Sorting (10k)” (https://www.10xgenomics.com/datasets/pbmc-from-a-healthy-donor-granulocytes-removed-through-cell-sorting-10-k-1-standard-2-0-0); “PBMC from a Healthy Donor - No Cell Sorting (10k)” (https://www.10xgenomics.com/datasets/pbmc-from-a-healthy-donor-no-cell-sorting-10-k-1-standard-2-0-0); “PBMC from a Healthy Donor - Granulocytes Removed Through Cell Sorting (3k)” (https://www.10xgenomics.com/datasets/pbmc-from-a-healthy-donor-granulocytes-removed-through-cell-sorting-3-k-1-standard-2-0-0); and “PBMC from a Healthy Donor - No Cell Sorting (3k)” (https://www.10xgenomics.com/datasets/pbmc-from-a-healthy-donor-no-cell-sorting-3-k-1-standard-2-0-0). The cell-type-matched intact Hi-C loop calls are available from the ENCODE4 data portal as four experiments, `ENCSR847RHU` (file `ENCFF281CTW`, B cell), `ENCSR335JYP` (`ENCFF749LFC`, CD4-positive alpha-beta T cell), `ENCSR321BHC` (`ENCFF403GHY`, CD8-positive alpha-beta T cell) and `ENCSR236EYO` (`ENCFF214LRJ`, CD14-positive monocyte). No ENCODE scMultiome sample is used in this study. The activated B-cell multiome is available from the IGVF Data Portal (https://data.igvf.org) as seven released analysis sets from the Satpathy laboratory, one per 10x channel: `IGVFDS0793KZKS`, `IGVFDS4931IGGD`, `IGVFDS3005LSBM`, `IGVFDS4731QGJR`, `IGVFDS7016ZFVC`, `IGVFDS6443DIXD` and `IGVFDS5437FJHN`. The multiplexed sample of the first channel is `IGVFSM3641EGVH`, and its component measurement sets are `IGVFDS1570UJMU` (single-nucleus RNA), `IGVFDS8103FALC` (single-nucleus ATAC) and `IGVFDS7563XAXA` (lipid-conjugated oligo sequencing for MULTI-seq demultiplexing). The deposit spans three donors, `IGVFDO0938OYMI`, `IGVFDO1264SGCS` and `IGVFDO1718HCES`, multiplexed together in the same libraries and demultiplexed by MULTI-seq. The two analyzed here are those labelled `donor1` and `donor2` in the sample aliases. The SEA-AD middle temporal gyrus data are available from the SEA-AD open registry on Amazon S3 (Sage Bionetworks `syn26223298`) [17]. The CITE-seq data are available from GEO under accession GSE164378 [8]. The bone-marrow CD34+ hematopoietic progenitor multiome [28] is available from GEO (replicate 1, `GSM6005306` and `GSM6005307`; replicate 2, `GSM6005308` and `GSM6005309`; each a paired RNA and ATAC multiome). The NeurIPS 2021 bone-marrow mononuclear cell multiome [38] is available from GEO under accession GSE194122. The OneK1K single-cell eQTL fine-mapping [23] is available from the eQTL Catalogue (study `QTS000038`) and from Zenodo (DOI 10.5281/zenodo.7619795).

## Code availability

scOnion is an R package, released as version 0.99.2 under the MIT license at https://github.com/ziqistats/scOnion. It installs with `BiocManager::install("ziqistats/scOnion")`, which resolves its CRAN and Bioconductor dependencies together, and ships the hg38 TSS annotation, a vignette and a citation record. The entry points used for every result here except the sim-ulation grid are `Onion()` and `OnionLinkageTable()`, with the rank convention of each analysis set by `reconcile.ranks` (Rank selection). The version used here is archived on Zenodo (DOI 10.5281/zenodo.XXXXXXX).

## 7 Supplementary Notes

### Supplementary Note 1 The scOnion solution, its relation to classical PCA, and uniqueness of the optimizers

#### Relation to classical PCA

Applying PCA to each modality separately maximizes the variance captured within that modality, but the two coordinate systems generally do not correspond across modalities

The objective in 1 instead couples them. Each of **Û** and **V̂** captures the dominant cell-level variance of its modality, while maximizing the inner product forces the retained directions to align across modalities. This construction relies on an equivalence between expressing the PCA objective through the original data matrix and through the cell Gram matrix, formalized in the following proposition.

#### Proposition S1

*The data matrix form and Gram matrix form of PCA,* 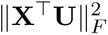 and 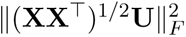, *have the same maximizers (and likewise for* **Y**).

*Proof.* It follows easily from the fact that

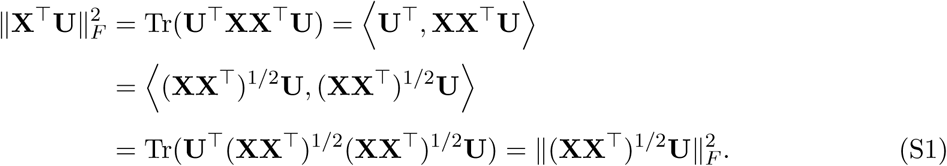

#### Theorem S2

*Let* (**XX**^T^)^1*/*2^(**YY**^T^)^1*/*2^ = **UΣV**^T^ *be its singular value decomposition, and* **Û**, **V̂** *contain the first r left and right singular vectors of* (**XX**^T^)^1*/*2^(**YY**^T^)^1*/*2^*, respectively. Then* **Û**, **V̂** *solve the optimization in 1*.

*Proof.* By definition of SVD, **Σ** is a diagonal matrix consisting of the singular values of (**XX**^T^)^1*/*2^(**YY**^T^)^1*/*2^ arranged in a non-increasing fashion.

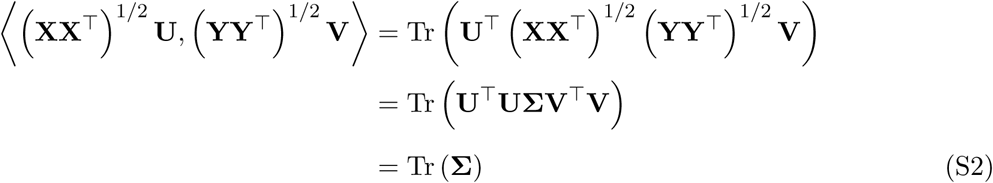

By the monotonicity of the singular values arranged in **Σ**, the trace and consequently the inner product are guaranteed to be maximized.□

#### Uniqueness of U and V

Similar to the conventional PCA, **U** and **V** are unique up to a joint rotation. For any orthonormal rotation matrix **R**, if **U**, **V** are optimizers, then **UR**, **VR** are also optimizers due to the invariance of trace under cyclic permutation of matrices. Here, we show that this rotation-induced non-uniqueness does not influence the construction of the similarity scores, joint cell embedding, and the joint feature embedding.

#### Invariance of the per-cell similarity score

Let **u***_i_,* **v***_i_* denote the *i*-th cell’s coordinates in the respective embedding. Let *d*(**u***_i_,* **v***_i_*) be a metric that defines the similarity score at the *i*-th cell. For any rotation matrix **R**, the rotated coordinates are **u***_i_***R**, **v***_i_***R**. Some metrics that satisfy

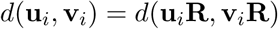

are Euclidean distance, dot product, and cosine similarity, where the measurement is preserved under orthogonal rotation.

- Euclidean distance:

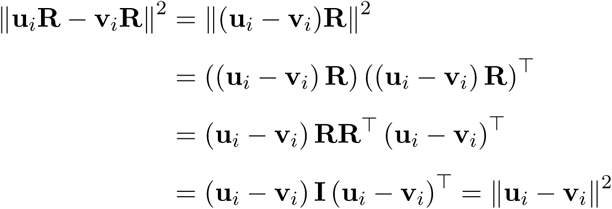

- Dot product: 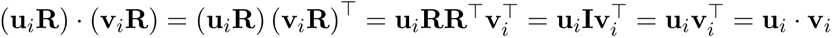
- Cosine similarity:

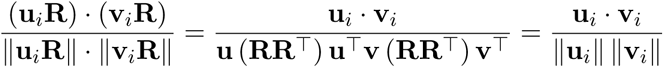

#### Invariance of the joint cell embedding

We show that the pairwise Euclidean distances between all cells are preserved under a joint rotation **R**. Let

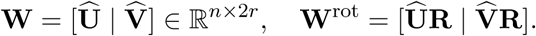

For any *i, j*, we want to show

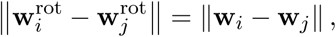

where **w***_i_* = [**u***_i_*|**v***_i_*] is the *i*-th row of **W**, or the original embedding coordinates for the *i*-th cell. Let us define

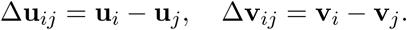

Then,

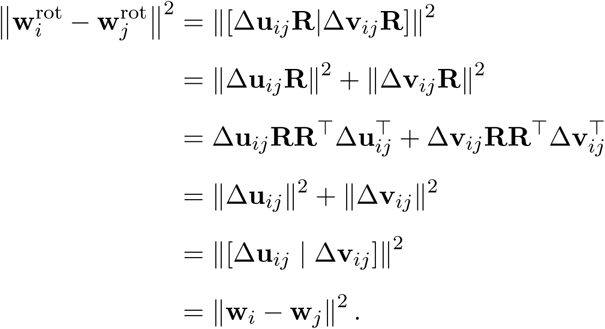

Since the pairwise Euclidean distance is preserved for all cells, it follows that UMAP is also invariant under such joint rotation, though one might not want to directly use the UMAP coordinates for downstream analyses.

#### Invariance of the joint feature embedding

The gene and peak embeddings are the projections **X**^T^**W** and **Y**^T^**W** onto the joint cell embedding. Under a joint rotation the cell embedding becomes 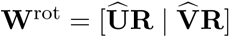, so

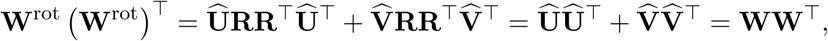

and the cross-modal loading product **X**^T^**WW**^T^**Y** is therefore unchanged. Because the shared component loadings **P** and **Q** and the singular-value weights *d* used by 7 are the singular value decomposition of that fixed product, the linkage score is unaffected as well, up to the sign and rotation ambiguity carried by any singular value decomposition.

### Supplementary Note 2 The hard-thresholding denoiser

For any matrix **X** of rank *n*, let **UΣV**^T^ denote its singular value decomposition, where **U** and **V** are the respective left and right singular vectors, and **Σ** is the diagonal matrix of singular values *σ*_1_ ≥ · · · ≥ *σ_n_*. Using this decomposition, **X** can also be written as 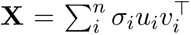, where *u_i_, v_i_* are the respective *i*-th left and right singular vectors, and *σ_i_* is the *i*-th singular value. To denoise **X** and its corresponding covariance matrix, we choose a threshold *T* that determines the number of retained components such that

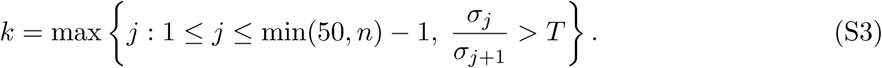

The resulting reconstructed, denoised matrix 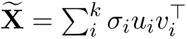 and 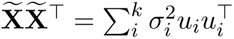 give the best rank-*k* approximation of **X** and **XX**^T^, respectively. The threshold *T* is a user-defined parameter balancing the trade-off between signal retention and noise reduction. In practice the usable range for single-cell multiomic data is 1.0025 ≤ *T* ≤ 1.02 (Supplementary Fig. 2d), and we use *T*_1_ = *T*_2_ = 1.01 by default, which selected 12 to 31 components in the whole-sample scMultiome fits reported here and 8 to 13 in cell-type subsets, matching Methods (Rank selection). The per-fit ranks are in Supplementary Table 3.

### Supplementary Note 3 Closed-form leave-one-cell-out standard error for the linkage score

The per-link standard error used in *z*_dev_ (8) is a fixed-embedding jackknife that does not refit scOnion. We write the raw linkage score of a gene *g* and a peak *p* as a bilinear form in the joint cell embedding 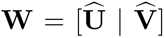. With the cross-modality cell kernel 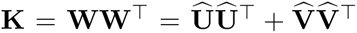 and the rank-*r* denoised gene and peak vectors 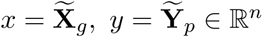,

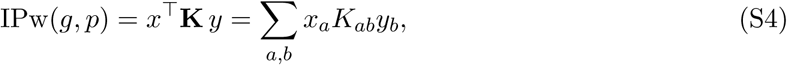

which equals the feature-space form 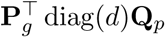 diag(*d*)**Q***_p_* because **X**^T^**WW**^T^**Y** = **P d**iag(*d*)**Q**^T^. The singular-value weights *d* enter only that feature-space form, not the cell kernel **K**. Removing cell *i* from the fitted score, without relearning **Û** or **V̂**, deletes every term in which cell *i* appears on either side,

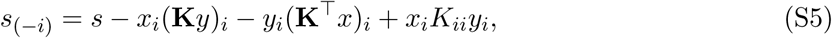

where the final term restores the (*i, i*) contribution that is otherwise subtracted twice. All *n* omitted-cell scores are obtained at once,

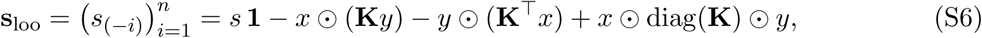

where ○ denotes element-wise multiplication, so the jackknife requires one full score and two matrix-vector products rather than *n* refits of the model. The pair-specific standard error is

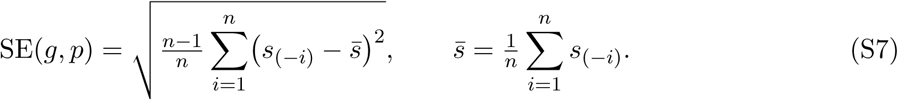

### Supplementary Note 4 Scope of the two simulation regimes

Two main simulations are used in this study. The construction of each is given in Methods and the grid settings in Supplementary Table 3.

#### Modality-uneven cell-state structure (**Fig. 3b** and Extended Data Fig. 2a,b)

The four ground-truth groups are the corners of a plane spanned by two factors, lineage and maturity. RNA carries lineage and ATAC carries maturity, and each factor is also added to the other modality at a signal ratio from 0 to 1, which defines the 5 × 5 grid. Its two extreme corners are the regimes the comparison separates. At ratios of 1 both factors are fully present in both modalities, so every method sees the same structure twice. At ratios of 0 each modality carries one factor, so the four groups can be recovered only by combining the modalities.

#### Graded cross-modality divergence (Extended Data Fig. 1c)

This simulation makes the divergence between the two modalities a known per-cell quantity and removes measurement noise entirely, so that the estimated concordance can be compared against a value that is exact by construction. Because the rotation angle is assigned independently of group membership, with the same number of cells at every combination of group and angle level, the true mean concordance of every group is identical by construction, and any difference between groups in the estimate is estimation error. The estimation spread of concordance in this simulation grows with the angle, and the growth has two multiplicative sources. This is because the estimation error in the angle itself is proportional to the angle, so the spread of the estimated angle widens from 3.9^◦^ at a true angle of 5^◦^ to 8.4^◦^ at 45^◦^.

### Supplementary Note 5 Guidance on parameter choice, score selection and application

#### Choice of rank and of the denoising threshold

The rank is selected by the ratio rule 3, so the user may set different thresholds. The sensitivity analysis is described in Methods and its parameters are given in Supplementary Table 3. The threshold enters the fit only through the rank that 3 returns, so two thresholds that select the same rank give the same fit.

#### Sequencing depth

Per-cell concordance carries a positive association with ATAC library size, so depth must be treated as a covariate. Where the question is whether a concordance difference between populations survives depth, the appropriate control is to equalize depth and refit rather than to regress depth out of the fitted score. The thinning procedure, its target depth and its outcome are given in Methods (Depth and hyperparameter sensitivity) and Extended Data Fig. 1b.

#### Scope of the linkage readouts

The distance-aware statistic *z*_dev_ of 8 is a ranking statistic rather than a calibrated probability or false-discovery rate, and it applies to feature pairs that carry genomic coordinates. For pairs without one, such as protein–gene pairs, the cosine readout and its standard error are used (Methods, Per-link jackknife standard error). The feature embedding draws its linkage signal from the cell-state variation captured by the leading joint directions, so the readouts are designed for populations that span several cell states, the whole sample or a compartment of related states, as in every analysis reported here.

#### Linkage-score readouts

scOnion reads the linkage between two features from their loadings in the joint embedding. This readout combines a raw similarity with three independent adjustments, so a small family of scores is available from the same feature embedding and the same closed-form standard error. The raw similarity is either the singular-value-weighted inner product IPw of 7, which preserves linkage magnitude, or its cosine normalization, which keeps only the direction of linkage and removes between-feature abundance differences. Either can be shrunk toward zero by its uncertainty through empirical-Bayes shrinkage with the weight *ω*, giving a variance-stabilized companion. Where a genomic-distance baseline is defined, as for gene–peak pairs, the raw score is standardized within distance strata as *z*_dev_. Any score can also be reported as an uncertainty-aware two-standard-error lower score score − 2 SE, which deprioritizes links supported by few cells.

### Supplementary Note 6 Extended methods

#### Per-cell concordance score

##### Comparison with Tilted-CCA synchrony and WNN modality weight

Concordance was compared with two existing per-cell measures on the sorted PBMC 10K sample (Extended Data Fig. 1d,e), the RNA modality weight of WNN [8] and the synchrony score of Tilted-CCA [6], both computed on the same cells from the RNA principal-component and ATAC LSI embeddings excluding the first LSI component. The WNN weight is the per-cell RNA modality weight re-turned by `FindMultiModalNeighbors`, and the Tilted-CCA synchrony score is the coefficient of determination from regressing each cell’s common-component profile in the anchor modality on the other modality’s profile rotated into that space (`compute synchrony`), from the Tilted-CCA fit described in Joint cell embedding benchmark. Because all three quantities partly reflect cell type, the 9,511 annotated cells were grouped into six immune lineages (monocyte, dendritic, B, CD4 T with regulatory T cells, CD8 T and NK) and the Spearman correlation of concordance with each measure was computed within lineage. The same correlation was recomputed after centering within the finer Azimuth labels (Extended Data Fig. 1e, bootstrap intervals in Supplementary Table 4). Tilted-CCA synchrony is defined relative to an anchor modality, and we report the RNA-anchored default because the Azimuth labels that form its metacells and the six lineages are transferred through RNA.

#### Gene–peak linkage benchmark against Hi-C and eQTL

##### Linkage-score ablation (Extended Data Fig. 4b)

The distance-controlled AUROC (Methods, Gene–peak linkage benchmark against Hi-C and eQTL) was applied to six score variants, crossing three adjustments (none, empirical-Bayes *ω*, and *z*_dev_) with two bases, the magnitude-carrying IPw and its distance-weighted cosine IPw*/*(‖*g‖_d_*‖*p*‖*_d_*) with 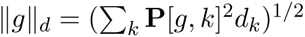, on both truth sets in each cell type. Because the jackknife is conditional on the fixed embedding, the norms carry no jackknife variance and SE_cos_ = SE*/*(‖*g‖_d_*‖*p‖_d_*) exactly. The ablation isolates whether the gain of *z*_dev_ over the unadjusted IPw comes from the distance-matched centering or from the standard error alone (per-cell-type values in Supplementary Table 4).

#### B-cell fate bifurcation

##### Residualization of concordance (Extended Data Fig. 5b)

To ask whether the decrease in the commitment window is technical, concordance was regressed on a depth covariate, the log_10_ per-cell sum of log-normalized RNA expression, an immediate-early-gene stress score [56] and S-phase and G2M scores, each the mean expression over the Seurat cell-cycle gene set, by ordinary least squares, and the residual was re-examined along pseudotime. For each lineage the cells were split into ten equal-sized groups along pseudotime, and dip depth was tested by two-sided Wilcoxon rank-sum test comparing the group with the lowest mean residualized concordance with the last group (Supplementary Table 4).

##### Technical covariates of the leading components (Supplementary Fig. 5c)

To ask whether a component tracks technical variation, each donor 2 cell was scored on components 1 to 3 by the mean of its RNA-side and ATAC-side coordinate and related by univariate *R*^2^ to five covariates computed from the log-normalized matrices, total RNA and ATAC signal, genes detected, percent mitochondrial signal and a cell-cycle score from the pooled Seurat S and G2M gene sets [57].

#### CITE-seq protein–gene coupling

##### Permuted-pairing control (Extended Data Fig. 6a)

As the CITE-seq analogue of Syn-thetic cell mismatching, an increasing fraction of RNA–ADT pairings was permuted at five levels from 0% to 100% in 25% steps, each level an independent draw under its own seed in which the selected cells’ ADT profiles were permuted among themselves. scOnion was refit at each level and concordance recomputed. The destroyed-pairing floor was established in each donor from 20 full permutations (null medians 0.527 to 0.530 against intact medians 0.72 to 0.75 across the four donors).

##### Recovery against transcript sparsity (Supplementary Fig. 6b)

Cognate-transcript detection was measured per donor as the fraction of that donor’s cells across all three time points with a nonzero raw count of the cognate most strongly linked to the protein. Proteins were assigned to quartiles of this measure once over the pooled protein-by-donor set (quartile breaks at 0.008, 0.052 and 0.203). Within each quartile, restricted to well-marking proteins with mean CLR abundance above the donor median, the top-5 recovery was computed by the unweighted cosine for scOnion and by raw correlation for the baseline.

##### Standard error of decoy and cognate links (Extended Data Fig. 6c)

Across the protein–donor cases whose top-cosine gene is not the cognate, the decoy’s standard error was compared with the cognate’s by paired one-sided Wilcoxon test (Supplementary Table 4).

## 8 Supplementary Tables

Six tables accompany this study. Supplementary Table 1 lists the datasets and the orthogonal vali-dation resources, Supplementary Table 2 the software and every non-default setting, Supplementary Table 3 the scOnion parameters used in each analysis, Supplementary Table 4 the statistical test behind each figure panel, Supplementary Table 5 the antibody-to-gene map, and Supplementary Table 6 the fate-regulator gene set.

**Supplementary Table 1:** Datasets and validation resources. Counts are the cells and features of the object each analysis was run on; analysis-specific subsets are in Supplementary Table 3. Full URLs are in Data availability.

| Dataset | Accession or source | Donors or replicates | Cells and features | Annotation source |
|---|---|---|---|---|
| 10x PBMC 10K, granulocyte-sorted, multiome | 10x Genomics Datasets portal | One healthy donor | 10,280 cells, 36,601 genes and 143,883 peaks | Azimuth level-2 labels [8] via pbmcMultiome.SeuratData |
| 10x PBMC 10K, unsorted, multiome | 10x Genomics Datasets portal | One healthy donor | 10,489 cells | As above |
| 10x PBMC 3K, granulocyte-sorted, multiome | 10x Genomics Datasets portal | One healthy donor | 2,490 cells | As above |
| 10x PBMC 3K, unsorted, multiome | 10x Genomics Datasets portal | One healthy donor | 2,554 cells | As above |
| SEA-AD middle temporal gyrus snMultiome | SEA-AD registry (Amazon S3); Synapse syn26223298; per-donor final-nuclei objects, RNA 2024-02-13, ATAC 2024-12-06 | Two donors, H20.33.015, H20.33.013 | 4,742 and 4,785 nuclei, 22,837 and 23,181 genes, 218,874 and 218,879 peaks | SEA-AD metadata, Subclass and Class; EnsDb.Hsapiens.v86, UCSC style, hg38 |
| IGVF activated B-cell multiome | IGVF Data Portal; analysis sets IGVFDS0793KZKS, IGVFDS4931IGGD, IGVFDS3005LSBM, IGVFDS4731QGJR, IGVFDS7016ZFVC, IGVFDS6443DIXD, IGVFDS5437FJHN | Two donors, donor1, donor2 | 10,058 and 17,941 cells (donor 2 Slingshot lineages 15,836 and 12,181 cells), 15,352 genes and 167,404 peaks | Cell states Naive, ActB, prePB, PB, preGCBC; lab-annotation available upon request |
| Hao PBMC CITE-seq, day 0, single batch | GEO GSE164378 | Four donors, biological replicates | 6,443, 5,978, 4,698 and 5,307 cells, 14,270 genes and 228 antibodies; ranking universes 15,648, 15,420, 15,237 and 15,238 genes | Cognate map, Supplementary Table 5; cell-type labels as published |
| Bone-marrow CD34+ progenitor multiome [28] | GEO; replicate 1 GSM6005306, GSM6005307; replicate 2 GSM6005308, GSM6005309 | Two technical replicates | 4,451 and 4,176 cells, 13,106 and 13,181 genes, 125,610 and 123,742 peaks | States and Palantir [39] pseudotime, four lineages, as published |
| NeurIPS 2021 bone-marrow mononuclear cell multiome [38] | GEO GSE194122 | 13 batches | 69,249 cells (24,997 batch-stratified for ED Fig. 3c–e), 13,431 genes and 116,490 peaks | None (annotation-free) |
| Validation resource | Accession (experiment / file) | Scope | Positive-set rule |  |
| ENCODE4 intact Hi-C loops [21, 22] | ENCSR847RHU / ENCFF281CTW | B cell, resting primary-cell reference | HiCCUPS loops; donut FDR ≤ 0.01; span 10 kb–1 Mb; 39,970 positives |  |
| ENCODE4 intact Hi-C loops | ENCSR335JYP / ENCFF749LFC | CD4-positive alpha-beta T cell | HiCCUPS loops; donut FDR ≤ 0.01; span 10 kb–1 Mb; 19,087 positives |  |
| ENCODE4 intact Hi-C loops | ENCSR321BHC / ENCFF403GHY | CD8-positive alpha-beta T cell | HiCCUPS loops; donut FDR ≤ 0.01; span 10 kb–1 Mb; 42,547 positives |  |
| ENCODE4 intact Hi-C loops | ENCSR236EYO / ENCFF214LRJ | CD14-positive monocyte | HiCCUPS loops; donut FDR ≤ 0.01; span 10 kb–1 Mb; 16,760 positives |  |
| OneK1K single-cell eQTL fine-mapping [23] | eQTL Catalogue QTS000038 (GRCh38); Zenodo 10.5281/zenodo.7619795 | PBMC, 982 donors | SuSiE 95% credible sets; PIP > 0.1; max PIP over subtypes; positives 198 (B), 1,016 (CD4 T), 418 (CD8 T), 74 (CD14) |  |
| JASPAR2024 position frequency matrices [43] | JASPAR2024 | Transcription-factor binding motifs | TFBSTools; BSgenome.Hsapiens.UCSC.hg38 |  |

**Supplementary Table 2:**
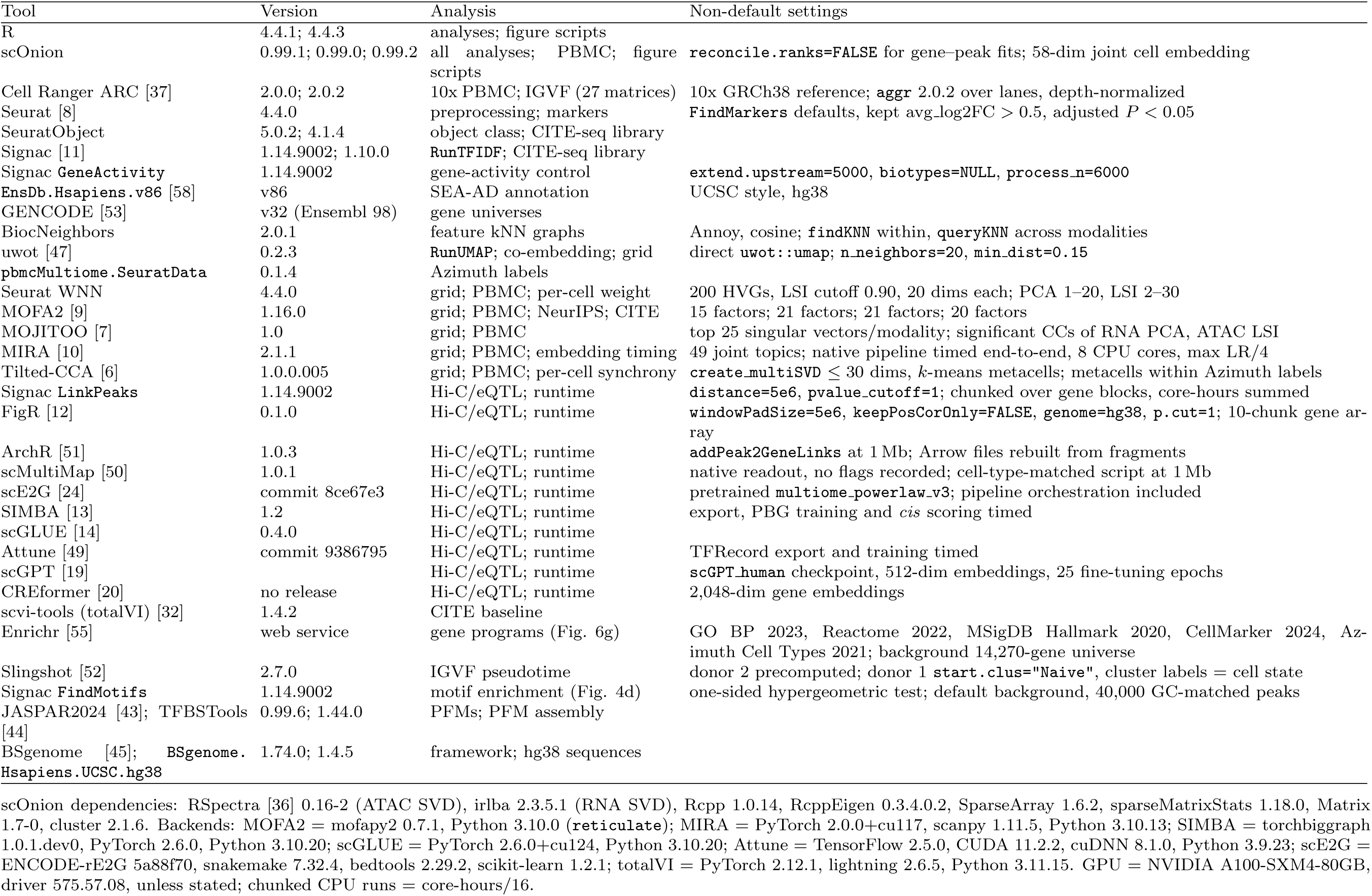
Software and non-default settings. Dependencies are listed below the table. Analyses: grid, Fig. 3b; PBMC, ED Fig. 3a,b; NeurIPS, ED Fig. 3c–e; per-cell, ED Fig. 1d,e; Hi-C/eQTL, Fig. 4b,c; runtime, Fig. 4e; CITE, Fig. 6. ED, Extended Data.

| Tool | Version | Analysis | Non-default settings |
|---|---|---|---|
| R | 4.4.1; 4.4.3 | analyses; figure scripts |  |
| scOnion | 0.99.1; 0.99.0; 0.99.2 | all analyses; PBMC; figure scripts | reconcile.ranks=FALSE for gene–peak fits; 58-dim joint cell embedding |
| Cell Ranger ARC [37] | 2.0.0; 2.0.2 | 10x PBMC; IGVF (27 matrices) | 10x GRCh38 reference; aggr 2.0.2 over lanes, depth-normalized |
| Seurat [8] | 4.4.0 | preprocessing; markers | FindMarkers defaults, kept avg_log2FC > 0.5, adjusted *P* < 0.05 |
| SeuratObject | 5.0.2; 4.1.4 | object class; CITE-seq library |  |
| Signac [11] | 1.14.9002; 1.10.0 | RunTFIDF; CITE-seq library |  |
| Signac GeneActivity | 1.14.9002 | gene-activity control | extend.upstream=5000, biotypes=NULL, process_n=6000 |
| EnsDb.Hsapiens.v86 [58] | v86 | SEA-AD annotation | UCSC style, hg38 |
| GENCODE [53] | v32 (Ensembl 98) | gene universes |  |
| BiocNeighbors | 2.0.1 | feature kNN graphs | Annoy, cosine; findKNN within, queryKNN across modalities |
| uwot [47] | 0.2.3 | RunUMAP; co-embedding; grid | direct uwot::umap; n_neighbors=20, min_dist=0.15 |
| pbmcMultiome.SeuratData | 0.1.4 | Azimuth labels |  |
| Seurat WNN | 4.4.0 | grid; PBMC; per-cell weight | 200 HVGs, LSI cutoff 0.90, 20 dims each; PCA 1–20, LSI 2–30 |
| MOFA2 [9] | 1.16.0 | grid; PBMC; NeurIPS; CITE | 15 factors; 21 factors; 21 factors; 20 factors |
| MOJITOO [7] | 1.0 | grid; PBMC | top 25 singular vectors/modality; significant CCs of RNA PCA, ATAC LSI |
| MIRA [10] | 2.1.1 | grid; PBMC; embedding timing | 49 joint topics; native pipeline timed end-to-end, 8 CPU cores, max LR/4 |
| Tilted-CCA [6] | 1.0.0.005 | grid; PBMC; per-cell synchrony | create_multiSVD ≤ 30 dims, *k*-means metacells; metacells within Azimuth labels |
| Signac LinkPeaks | 1.14.9002 | Hi-C/eQTL; runtime | distance=5e6, pvalue_cutoff=1; chunked over gene blocks, core-hours summed |
| FigR [12] | 0.1.0 | Hi-C/eQTL; runtime | windowPadSize=5e6, keepPosCorOnly=FALSE, genome=hg38, p.cut=1; 10-chunk gene ar-ray |
| ArchR [51] | 1.0.3 | Hi-C/eQTL; runtime | addPeak2GeneLinks at 1 Mb; Arrow files rebuilt from fragments |
| scMultiMap [50] | 1.0.1 | Hi-C/eQTL; runtime | native readout, no flags recorded; cell-type-matched script at 1 Mb |
| scE2G [24] | commit 8ce67e3 | Hi-C/eQTL; runtime | pretrained multiome_powerlaw_v3; pipeline orchestration included |
| SIMBA [13] | 1.2 | Hi-C/eQTL; runtime | export, PBG training and *cis* scoring timed |
| scGLUE [14] | 0.4.0 | Hi-C/eQTL; runtime |  |
| Attune [49] | commit 9386795 | Hi-C/eQTL; runtime | TFRecord export and training timed |
| scGPT [19] |  | Hi-C/eQTL; runtime | scGPT_human checkpoint, 512-dim embeddings, 25 fine-tuning epochs |
| CREformer [20] | no release | Hi-C/eQTL; runtime | 2,048-dim gene embeddings |
| scvi-tools (totalVI) [32] | 1.4.2 | CITE baseline |  |
| Enrichr [55] | web service | gene programs (Fig. 6g) | GO BP 2023, Reactome 2022, MSigDB Hallmark 2020, CellMarker 2024, Az-imuth Cell Types 2021; background 14,270-gene universe |
| Slingshot [52] | 2.7.0 | IGVF pseudotime | donor 2 precomputed; donor 1 start.clus="Naive", cluster labels = cell state |
| Signac FindMotifs | 1.14.9002 | motif enrichment (Fig. 4d) | one-sided hypergeometric test; default background, 40,000 GC-matched peaks |
| JASPAR2024 [43]; TFBSTools [44] | 0.99.6; 1.44.0 | PFMs; PFM assembly |  |
| BSgenome [45]; BSgenome.Hsapiens.UCSC.hg38 | 1.74.0; 1.4.5 | framework; hg38 sequences |  |

scOnion dependencies: RSpectra [36] 0.16-2 (ATAC SVD), irlba 2.3.5.1 (RNA SVD), Rcpp 1.0.14, RcppEigen 0.3.4.0.2, SparseArray 1.6.2, sparseMatrixStats 1.18.0, Matrix 1.7-0, cluster 2.1.6. Backends: MOFA2 = mofapy2 0.7.1, Python 3.10.0 (reticulate); MIRA = PyTorch 2.0.0+cu117, scanpy 1.11.5, Python 3.10.13; SIMBA = torchbiggraph 1.0.1.dev0, PyTorch 2.6.0, Python 3.10.20; scGLUE = PyTorch 2.6.0+cu124, Python 3.10.20; Attune = TensorFlow 2.5.0, CUDA 11.2.2, cuDNN 8.1.0, Python 3.9.23; scE2G = ENCODE-rE2G 5a88f70, snakemake 7.32.4, bedtools 2.29.2, scikit-learn 1.2.1; totalVI = PyTorch 2.12.1, lightning 2.6.5, Python 3.11.15. GPU = NVIDIA A100-SXM4-80GB, driver 575.57.08, unless stated; chunked CPU runs = core-hours/16.

**Supplementary Table 3:**
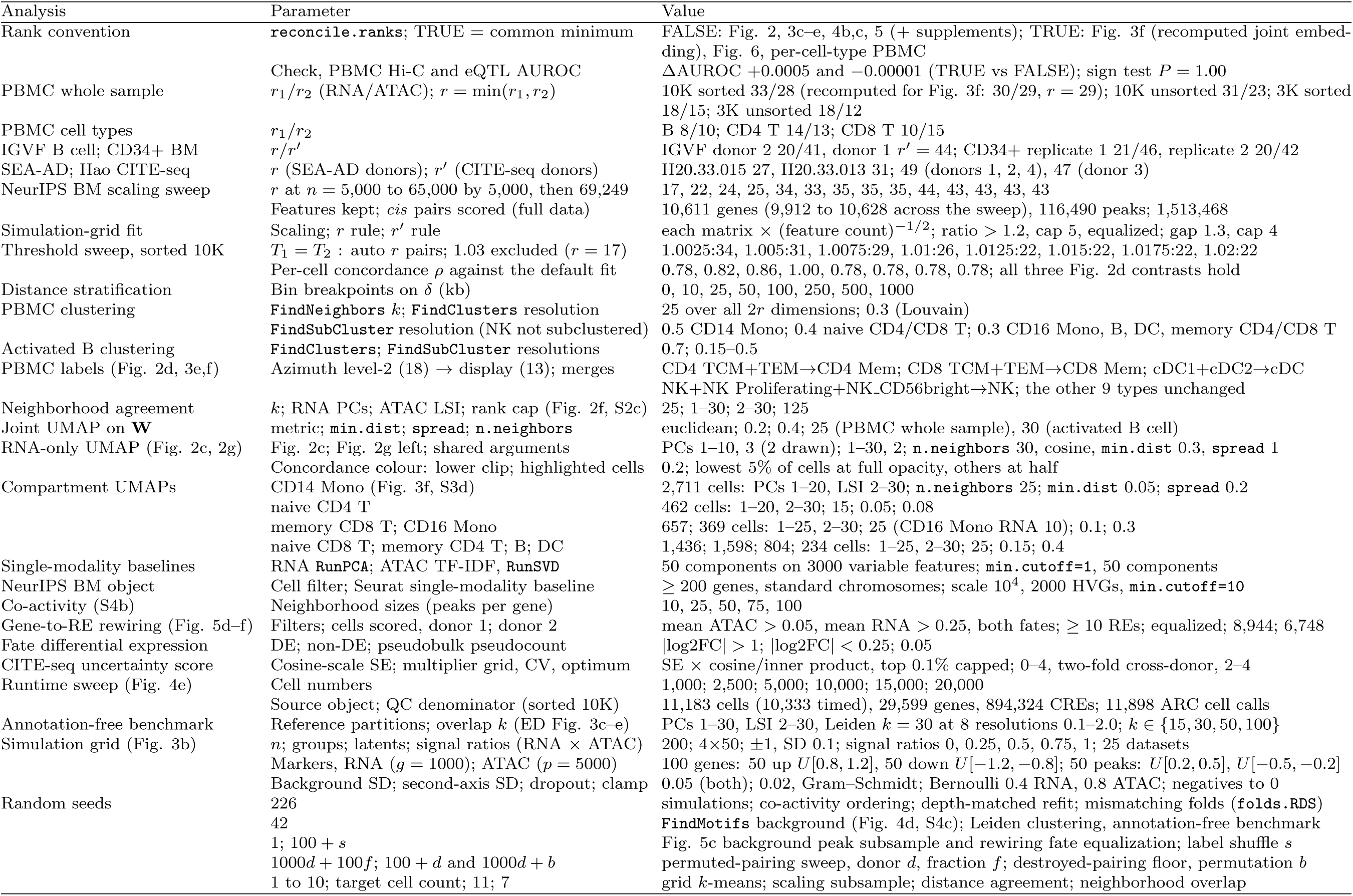
scOnion parameters by analysis. Defaults, all real data: *T*_1_ = *T*_2_ = 1.01, *T*_3_ = 1.1, ≤ 50 components per modality. *r*, *r^0^*: cell-and feature-embedding ranks; S, Supplementary Fig.; ED, Extended Data Fig. Settings given in Methods are not repeated.

| Analysis | Parameter | Value |
|---|---|---|
| Rank convention | reconcile.ranks; TRUE = common minimum | FALSE: Fig. 2, 3c–e, 4b,c, 5 (+ supplements); TRUE: Fig. 3f (recomputed joint embed-ding), Fig. 6, per-cell-type PBMC |
|  | Check, PBMC Hi-C and eQTL AUROC | ΔAUROC +0.0005 and −0.00001 (TRUE vs FALSE); sign test P = 1.00 |
| PBMC whole sample | r<sub>1</sub>/r<sub>2</sub> (RNA/ATAC); r = min(r<sub>1</sub>, r<sub>2</sub>) | 10K sorted 33/28 (recomputed for Fig. 3f: 30/29, r = 29); 10K unsorted 31/23; 3K sorted 18/15; 3K unsorted 18/12 |
| PBMC cell types | r<sub>1</sub>/r<sub>2</sub> | B 8/10; CD4 T 14/13; CD8 T 10/15 |
| IGVF B cell; CD34+ BM | r/r′ | IGVF donor 2 20/41, donor 1 r′ = 44; CD34+ replicate 1 21/46, replicate 2 20/42 |
| SEA-AD; Hao CITE-seq | r (SEA-AD donors); r′ (CITE-seq donors) | H20.33.015 27, H20.33.013 31; 49 (donors 1, 2, 4), 47 (donor 3) |
| NeurIPS BM scaling sweep | r at n = 5,000 to 65,000 by 5,000, then 69,249 | 17, 22, 24, 25, 34, 33, 35, 35, 35, 44, 43, 43, 43, 43 |
|  | Features kept; *cis* pairs scored (full data) | 10,611 genes (9,912 to 10,628 across the sweep), 116,490 peaks; 1,513,468 |
| Simulation-grid fit | Scaling; r rule; r′ rule | each matrix × (feature count)<sup>−1/2</sup>; ratio > 1.2, cap 5, equalized; gap 1.3, cap 4 |
| Threshold sweep, sorted 10K | T<sub>1</sub> = T<sub>2</sub>: auto r pairs; 1.03 excluded (r = 17) | 1.0025:34, 1.005:31, 1.0075:29, 1.01:26, 1.0125:22, 1.015:22, 1.0175:22, 1.02:22 |
|  | Per-cell concordance ρ against the default fit | 0.78, 0.82, 0.86, 1.00, 0.78, 0.78, 0.78, 0.78; all three Fig. 2d contrasts hold |
| Distance stratification | Bin breakpoints on δ (kb) | 0, 10, 25, 50, 100, 250, 500, 1000 |
| PBMC clustering | FindNeighbors k; FindClusters resolution | 25 over all 2r dimensions; 0.3 (Louvain) |
|  | FindSubCluster resolution (NK not subclustered) | 0.5 CD14 Mono; 0.4 naive CD4/CD8 T; 0.3 CD16 Mono, B, DC, memory CD4/CD8 T |
| Activated B clustering | FindClusters; FindSubCluster resolutions | 0.7; 0.15–0.5 |
| PBMC labels (Fig. 2d, 3e,f) | Azimuth level-2 (18) → display (13); merges | CD4 TCM+TEM→CD4 Mem; CD8 TCM+TEM→CD8 Mem; cDC1+cDC2→cDC<br>NK+NK Proliferating+NK_CD56bright→NK; the other 9 types unchanged |
| Neighborhood agreement | k; RNA PCs; ATAC LSI; rank cap (Fig. 2f, S2c) | 25; 1–30; 2–30; 125 |
| Joint UMAP on **W** | metric; min.dist; spread; n.neighbors | euclidean; 0.2; 0.4; 25 (PBMC whole sample), 30 (activated B cell) |
| RNA-only UMAP (Fig. 2c, 2g) | Fig. 2c; Fig. 2g left; shared arguments | PCs 1–10, 3 (2 drawn); 1–30, 2; n.neighbors 30, cosine, min.dist 0.3, spread 1 |
|  | Concordance colour: lower clip; highlighted cells | 0.2; lowest 5% of cells at full opacity, others at half |
| Compartment UMAPs | CD14 Mono (Fig. 3f, S3d) | 2,711 cells: PCs 1–20, LSI 2–30; n.neighbors 25; min.dist 0.05; spread 0.2 |
|  | naive CD4 T | 462 cells: 1–20, 2–30; 15; 0.05; 0.08 |
|  | memory CD8 T; CD16 Mono | 657; 369 cells: 1–25, 2–30; 25 (CD16 Mono RNA 10); 0.1; 0.3 |
|  | naive CD8 T; memory CD4 T; B; DC | 1,436; 1,598; 804; 234 cells: 1–25, 2–30; 25; 0.15; 0.4 |
| Single-modality baselines | RNA RunPCA; ATAC TF-IDF, RunSVD | 50 components on 3000 variable features; min.cutoff=1, 50 components |
| NeurIPS BM object | Cell filter; Seurat single-modality baseline | ≥ 200 genes, standard chromosomes; scale 10<sup>4</sup>, 2000 HVGs, min.cutoff=10 |
| Co-activity (S4b) | Neighborhood sizes (peaks per gene) | 10, 25, 50, 75, 100 |
| Gene-to-RE rewiring (Fig. 5d–f) | Filters; cells scored, donor 1; donor 2 | mean ATAC > 0.05, mean RNA > 0.25, both fates; ≥ 10 REs; equalized; 8,944; 6,748 |
| Fate differential expression | DE; non-DE; pseudobulk pseudocount | \|log2FC\| > 1; \|log2FC\| < 0.25; 0.05 |
| CITE-seq uncertainty score | Cosine-scale SE; multiplier grid, CV, optimum | SE × cosine/inner product, top 0.1% capped; 0–4, two-fold cross-donor, 2–4 |
| Runtime sweep (Fig. 4e) | Cell numbers | 1,000; 2,500; 5,000; 10,000; 15,000; 20,000 |
|  | Source object; QC denominator (sorted 10K) | 11,183 cells (10,333 timed), 29,599 genes, 894,324 CREs; 11,898 ARC cell calls |
| Annotation-free benchmark | Reference partitions; overlap k (ED Fig. 3c–e) | PCs 1–30, LSI 2–30, Leiden k = 30 at 8 resolutions 0.1–2.0; k ∈ {15, 30, 50, 100} |
| Simulation grid (Fig. 3b) | n; groups; latents; signal ratios (RNA × ATAC) | 200; 4×50; ±1, SD 0.1; signal ratios 0, 0.25, 0.5, 0.75, 1; 25 datasets |
|  | Markers, RNA (g = 1000); ATAC (p = 5000) | 100 genes: 50 up U[0.8, 1.2], 50 down U[−1.2, −0.8]; 50 peaks: U[0.2, 0.5], U[−0.5, −0.2] |
|  | Background SD; second-axis SD; dropout; clamp | 0.05 (both); 0.02, Gram–Schmidt; Bernoulli 0.4 RNA, 0.8 ATAC; negatives to 0 |
| Random seeds | 226 | simulations; co-activity ordering; depth-matched refit; mismatching folds (folds.RDS) |
|  | 42 | FindMotifs background (Fig. 4d, S4c); Leiden clustering, annotation-free benchmark |
|  | 1; 100 + s | Fig. 5c background peak subsample and rewiring fate equalization; label shuffle s |
|  | 1000d + 100f; 100 + d and 1000d + b | permuted-pairing sweep, donor d, fraction f; destroyed-pairing floor, permutation b |
|  | 1 to 10; target cell count; 11; 7 | grid k-means; scaling subsample; distance agreement; neighborhood overlap |

**Supplementary Table 4:**
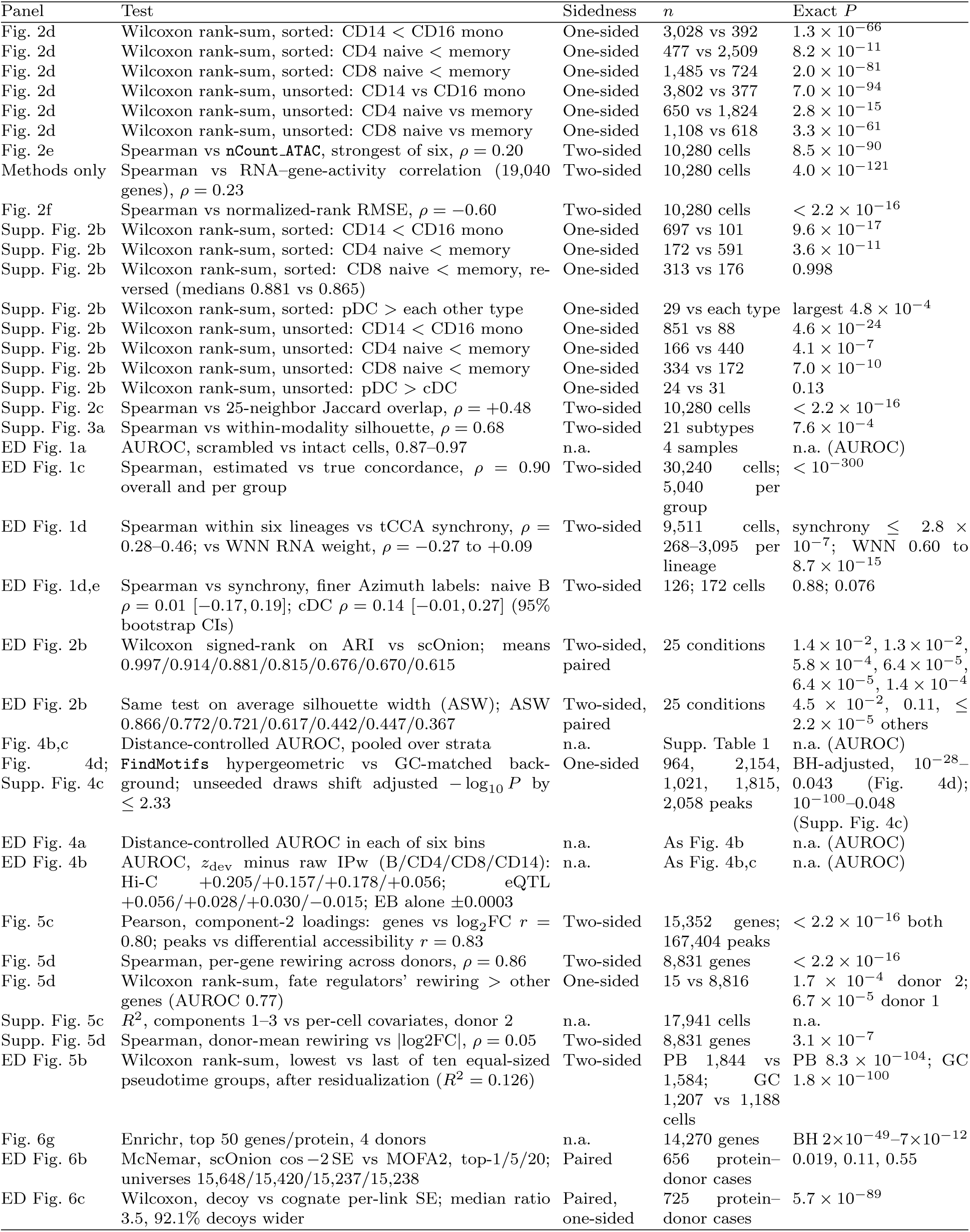
Statistical tests by figure panel. Panels reporting no test statistic (schematics, descriptive panels) are omitted; AUROC and *R*^2^ rows carry no *P*. ED Fig. 2b lists methods in the order scOnion, MIRA, CCA, MOFA2, MOJITOO, tCCA (Tilted-CCA), WNN. ED, Extended Data.

| Panel | Test | Sidedness | *n* | Exact *P* |
|---|---|---|---|---|
| Fig. 2d | Wilcoxon rank-sum, sorted: CD14 < CD16 mono | One-sided | 3,028 vs 392 | 1.3 × 10<sup>−66</sup> |
| Fig. 2d | Wilcoxon rank-sum, sorted: CD4 naive < memory | One-sided | 477 vs 2,509 | 8.2 × 10<sup>−11</sup> |
| Fig. 2d | Wilcoxon rank-sum, sorted: CD8 naive < memory | One-sided | 1,485 vs 724 | 2.0 × 10<sup>−81</sup> |
| Fig. 2d | Wilcoxon rank-sum, unsorted: CD14 vs CD16 mono | One-sided | 3,802 vs 377 | 7.0 × 10<sup>−94</sup> |
| Fig. 2d | Wilcoxon rank-sum, unsorted: CD4 naive vs memory | One-sided | 650 vs 1,824 | 2.8 × 10<sup>−15</sup> |
| Fig. 2d | Wilcoxon rank-sum, unsorted: CD8 naive vs memory | One-sided | 1,108 vs 618 | 3.3 × 10<sup>−61</sup> |
| Fig. 2e | Spearman vs nCount_ATAC, strongest of six, *ρ* = 0.20 | Two-sided | 10,280 cells | 8.5 × 10<sup>−90</sup> |
| Methods only | Spearman vs RNA–gene-activity correlation (19,040 genes), *ρ* = 0.23 | Two-sided | 10,280 cells | 4.0 × 10<sup>−121</sup> |
| Fig. 2f | Spearman vs normalized-rank RMSE, *ρ* = −0.60 | Two-sided | 10,280 cells | < 2.2 × 10<sup>−16</sup> |
| Supp. Fig. 2b | Wilcoxon rank-sum, sorted: CD14 < CD16 mono | One-sided | 697 vs 101 | 9.6 × 10<sup>−17</sup> |
| Supp. Fig. 2b | Wilcoxon rank-sum, sorted: CD4 naive < memory | One-sided | 172 vs 591 | 3.6 × 10<sup>−11</sup> |
| Supp. Fig. 2b | Wilcoxon rank-sum, sorted: CD8 naive < memory, re-versed (medians 0.881 vs 0.865) | One-sided | 313 vs 176 | 0.998 |
| Supp. Fig. 2b | Wilcoxon rank-sum, sorted: pDC > each other type | One-sided | 29 vs each type | largest 4.8 × 10<sup>−4</sup> |
| Supp. Fig. 2b | Wilcoxon rank-sum, unsorted: CD14 < CD16 mono | One-sided | 851 vs 88 | 4.6 × 10<sup>−24</sup> |
| Supp. Fig. 2b | Wilcoxon rank-sum, unsorted: CD4 naive < memory | One-sided | 166 vs 440 | 4.1 × 10<sup>−7</sup> |
| Supp. Fig. 2b | Wilcoxon rank-sum, unsorted: CD8 naive < memory | One-sided | 334 vs 172 | 7.0 × 10<sup>−10</sup> |
| Supp. Fig. 2b | Wilcoxon rank-sum, unsorted: pDC > cDC | One-sided | 24 vs 31 | 0.13 |
| Supp. Fig. 2c | Spearman vs 25-neighbor Jaccard overlap, *ρ* = +0.48 | Two-sided | 10,280 cells | < 2.2 × 10<sup>−16</sup> |
| Supp. Fig. 3a | Spearman vs within-modality silhouette, *ρ* = 0.68 | Two-sided | 21 subtypes | 7.6 × 10<sup>−4</sup> |
| ED Fig. 1a | AUROC, scrambled vs intact cells, 0.87–0.97 | n.a. | 4 samples | n.a. (AUROC) |
| ED Fig. 1c | Spearman, estimated vs true concordance, *ρ* = 0.90 overall and per group | Two-sided | 30,240 cells; 5,040 per group | < 10<sup>−300</sup> |
| ED Fig. 1d | Spearman within six lineages vs tCCA synchrony, *ρ* = 0.28–0.46; vs WNN RNA weight, *ρ* = −0.27 to +0.09 | Two-sided | 9,511 cells, 268–3,095 per lineage | synchrony ≤ 2.8 × 10<sup>−7</sup>; WNN 0.60 to 8.7 × 10<sup>−15</sup> |
| ED Fig. 1d,e | Spearman vs synchrony, finer Azimuth labels: naive B *ρ* = 0.01 [−0.17, 0.19]; cDC *ρ* = 0.14 [−0.01, 0.27] (95% bootstrap CIs) | Two-sided | 126; 172 cells | 0.88; 0.076 |
| ED Fig. 2b | Wilcoxon signed-rank on ARI vs scOnion; means 0.997/0.914/0.881/0.815/0.676/0.670/0.615 | Two-sided, paired | 25 conditions | 1.4 × 10<sup>−2</sup>, 1.3 × 10<sup>−2</sup>, 5.8 × 10<sup>−4</sup>, 6.4 × 10<sup>−5</sup>, 6.4 × 10<sup>−5</sup>, 1.4 × 10<sup>−4</sup> |
| ED Fig. 2b | Same test on average silhouette width (ASW); ASW 0.866/0.772/0.721/0.617/0.442/0.447/0.367 | Two-sided, paired | 25 conditions | 4.5 × 10<sup>−2</sup>, 0.11, ≤ 2.2 × 10<sup>−5</sup> others |
| Fig. 4b,c | Distance-controlled AUROC, pooled over strata | n.a. | Supp. Table 1 | n.a. (AUROC) |
| Fig. 4d; Supp. Fig. 4c | FindMotifs hypergeometric vs GC-matched back-ground; unseeded draws shift adjusted −log<sub>10</sub> *P* by ≤ 2.33 | One-sided | 964, 2,154, 1,021, 1,815, 2,058 peaks | BH-adjusted, 10<sup>−28</sup>–0.043 (Fig. 4d); 10<sup>−100</sup>–0.048 (Supp. Fig. 4c) |
| ED Fig. 4a | Distance-controlled AUROC in each of six bins | n.a. | As Fig. 4b | n.a. (AUROC) |
| ED Fig. 4b | AUROC, *z*<sub>dev</sub> minus raw IPw (B/CD4/CD8/CD14): Hi-C +0.205/+0.157/+0.178/+0.056; eQTL +0.056/+0.028/+0.030/−0.015; EB alone ±0.0003 | n.a. | As Fig. 4b,c | n.a. (AUROC) |
| Fig. 5c | Pearson, component-2 loadings: genes vs log<sub>2</sub>FC *r* = 0.80; peaks vs differential accessibility *r* = 0.83 | Two-sided | 15,352 genes; 167,404 peaks | < 2.2 × 10<sup>−16</sup> both |
| Fig. 5d | Spearman, per-gene rewiring across donors, *ρ* = 0.86 | Two-sided | 8,831 genes | < 2.2 × 10<sup>−16</sup> |
| Fig. 5d | Wilcoxon rank-sum, fate regulators’ rewiring > other genes (AUROC 0.77) | One-sided | 15 vs 8,816 | 1.7 × 10<sup>−4</sup> donor 2; 6.7 × 10<sup>−5</sup> donor 1 |
| Supp. Fig. 5c | *R*<sup>2</sup>, components 1–3 vs per-cell covariates, donor 2 | n.a. | 17,941 cells | n.a. |
| Supp. Fig. 5d | Spearman, donor-mean rewiring vs \|log2FC\|, *ρ* = 0.05 | Two-sided | 8,831 genes | 3.1 × 10<sup>−7</sup> |
| ED Fig. 5b | Wilcoxon rank-sum, lowest vs last of ten equal-sized pseudotime groups, after residualization (*R*<sup>2</sup> = 0.126) | Two-sided | PB 1,844 vs 1,584; GC 1,207 vs 1,188 cells | PB 8.3 × 10<sup>−104</sup>; GC 1.8 × 10<sup>−100</sup> |
| Fig. 6g | Enrichr, top 50 genes/protein, 4 donors | n.a. | 14,270 genes | BH 2×10<sup>−49</sup>–7×10<sup>−12</sup> |
| ED Fig. 6b | McNemar, scOnion cos −2 SE vs MOFA2, top-1/5/20; universes 15,648/15,420/15,237/15,238 | Paired | 656 protein–donor cases | 0.019, 0.11, 0.55 |
| ED Fig. 6c | Wilcoxon, decoy vs cognate per-link SE; median ratio 3.5, 92.1% decoys wider | Paired, one-sided | 725 protein–donor cases | 5.7 × 10<sup>−89</sup> |

**Supplementary Table 5:** Antibody-to-gene map for the 228-antibody TotalSeq-A panel, deposited as a file; columns below. Mapping against the HGNC complete set (genenames.org, accessed 19 June 2026) [54], with manual curation of the remainder.

| Column | Content |
|---|---|
| Antibody feature | ADT feature name (Hao object) with clone suffix; marker or isotype control |
| Cognate gene or genes | Encoding gene(s), all subunits of a complex; count |
| Mapping route | HGNC symbol, alias or previous-symbol match, or manual |
| Mapping provenance | Automated or curated; pre-curation hit set |
| HGNC validity | All cognate symbols current in HGNC |
| Per-donor scorability | Cognate genes in each donor universe; scorable yes/no |
| Genes absent | Cognate genes absent from the ranking universe |

**Supplementary Table 6:** Fate-regulator gene set. The 17 canonical B-cell fate genes used for the AUROC in Fig. 5d, and whether each carried a rewiring score in both donors. *SPIB* and *BCL2A1* did not and were excluded, giving *n* = 15 regulators scored against the remaining 8,816 genes. This curated set is not identical to the genes labelled in Fig. 5d, which the figure caption describes as the fate-associated genes highlighted in the original study.

| Gene | Rewiring score in both donors | Gene | Rewiring score in both donors |
|---|---|---|---|
| *PRDM1* | Yes | *MEF2B* | Yes |
| *IRF4* | Yes | *IRF8* | Yes |
| *XBP1* | Yes | *SPIB* | No, excluded |
| *MZB1* | Yes | *CD83* | Yes |
| *TNFRSF17* | Yes | *EBF1* | Yes |
| *BCL6* | Yes | *BCL2A1* | No, excluded |
| *BACH2* | Yes | *CCND2* | Yes |
| *PAX5* | Yes | *MYC* | Yes |
| *AICDA* | Yes |  |  |

## 9 Supplementary Figures

**Supplementary Figure 1:**
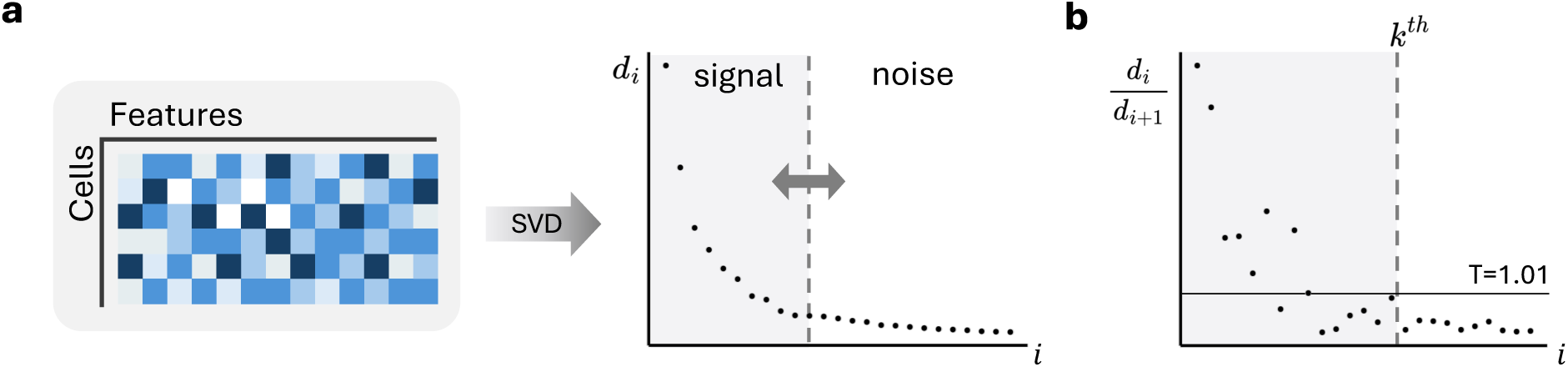
Data-adaptive singular-value thresholding for rank selection, applied at each singular value decomposition (SVD) truncation in scOnion (Methods). **a**, Denoising of a single modality. The centered and scaled cell-by-feature matrix is decomposed by SVD, and the ordered singular values are plotted against their index *i*. The spectrum decays steeply and then flattens, so the leading components (signal) are retained and the flat tail (noise) is truncated. **b**, The retained rank is read from the ratio of each singular value to the next. Components are kept while this ratio exceeds a threshold *T* and are dropped once the spectrum flattens, giving the largest index at which the ratio still exceeds *T* (3). The same rule sets both modality ranks at the default *T*_1_ = *T*_2_ = 1.01 (shown) and the feature-embedding rank *r*^′^ at *T*_3_ = 1.1 (Methods).

**Supplementary Figure 2:**
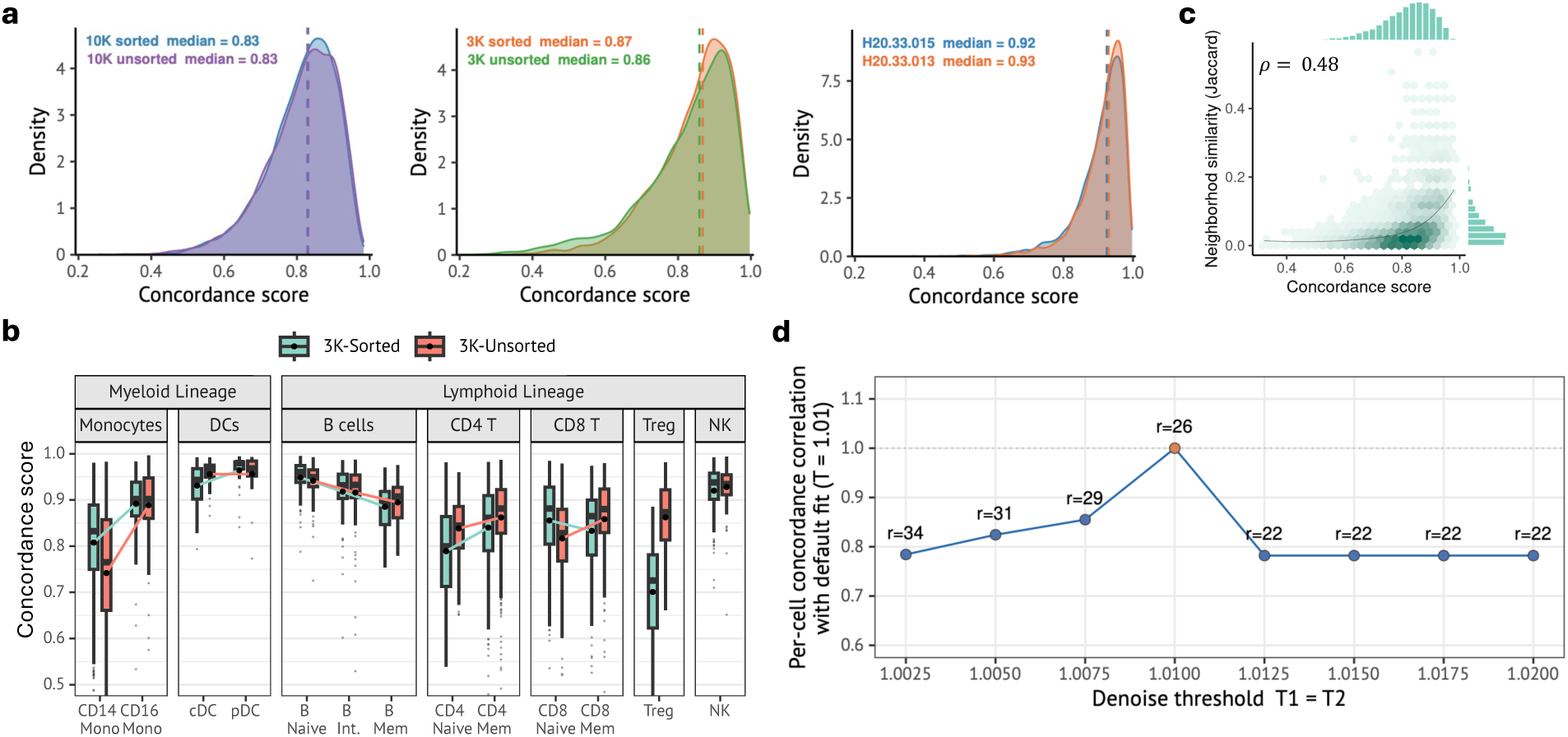
Concordance is reproducible across samples, tracks cross-modality neighborhood agreement, and is robust to the rank-selection threshold. **a**, Concordance score distributions for four 10x PBMC multiome samples (sorted and unsorted, 10K and 3K cells; median 0.83 to 0.87) and for a human middle temporal gyrus (MTG) multiome sample in two donors (H20.33.015 and H20.33.013; median 0.92 and 0.93). Median concordance is marked by the dashed line and is stable within a tissue and higher in MTG than in PBMC. **b**, Per-cell-type concordance for the 3K sorted and unsorted samples across myeloid and lymphoid populations. The CD14 versus CD16 monocyte and naive versus memory CD4 T cell contrasts of Fig. 2d are reproduced in both samples and pDCs rank highest in both, whereas the naive versus memory CD8 T cell contrast is reversed in the sorted 3K sample. **c**, Concordance versus cross-modality neighborhood similarity, the Jaccard overlap between each cell’s 25 nearest neighbors in the RNA embedding and in the ATAC embed-ding, with the cell itself excluded (two-sided Spearman *ρ* = +0.48, sorted PBMC 10K sample). Marginal histograms show the two distributions. **d**, Robustness of concordance to the scOnion de-noising threshold. Correlation of per-cell concordance with the default fit (*T*_1_ = *T*_2_ = 1.01) across the usable threshold range (1.0025 to 1.02), with the auto-selected rank *r* annotated. Concordance changes only at rank transitions, and every cell-type contrast is preserved at all thresholds (Methods). In all box plots the center line is the median, the box limits are the first and third quartiles (interquartile range, IQR), and the whiskers extend to the most extreme value within 1.5× the IQR. Outliers beyond the whiskers are plotted as individual points. Statistical tests and sample sizes are reported in Methods.

**Supplementary Figure 3:**
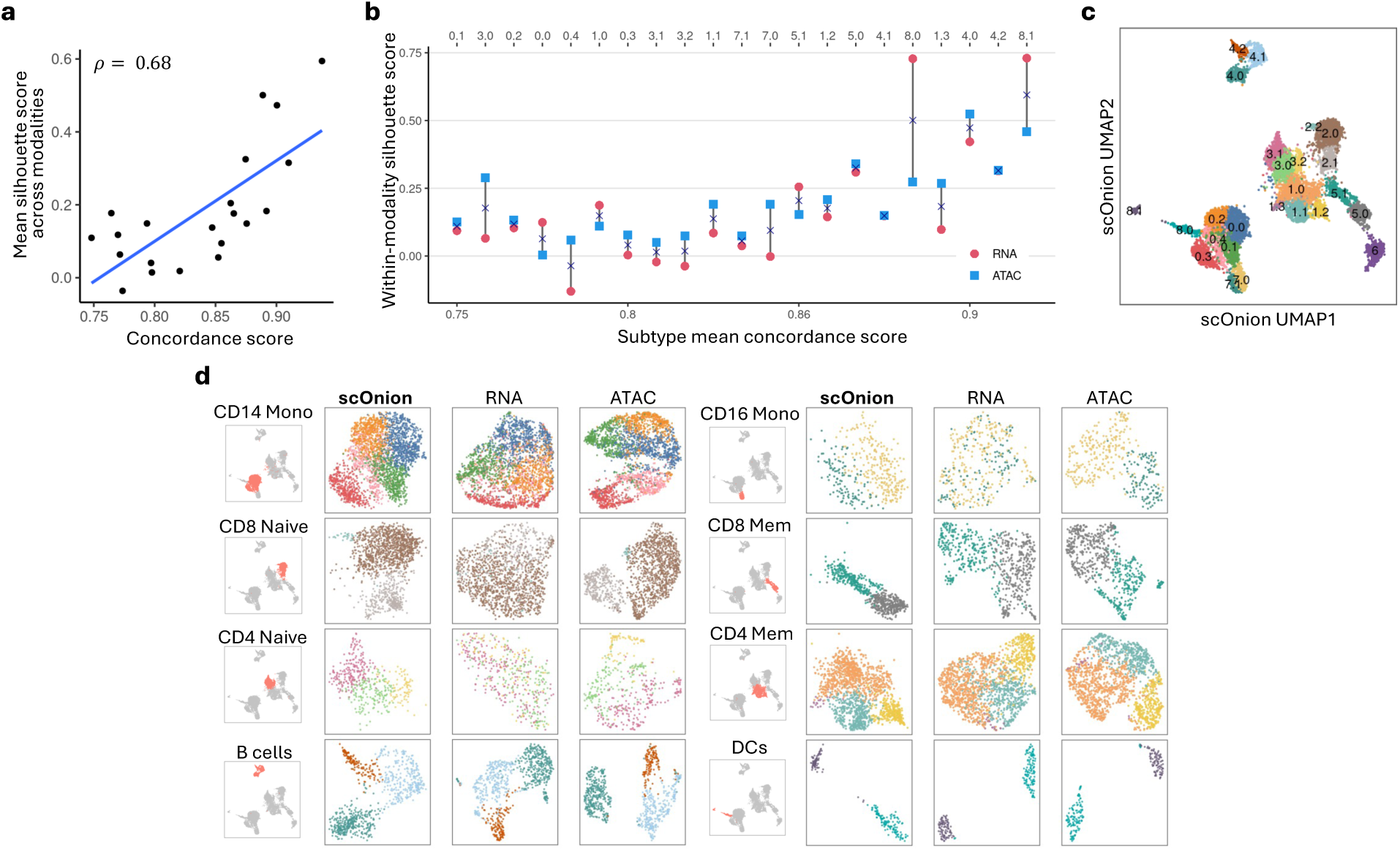
Concordance-guided subtyping within immune compartments. **a**, Mean within-modality silhouette score across RNA and ATAC versus concordance for scOnion subtypes in the sorted PBMC 10K sample (two-sided Spearman *ρ* = 0.68 over 21 subtypes; Methods). Higher-concordance populations tend to contain subtype separations supported by both modalities. **b**, Within-modality silhouette score in RNA and ATAC for each scOnion subtype, ordered by subtype mean concordance. Low-concordance subtypes tend to separate in one modality only. **c**, scOnion joint-embedding UMAP with fine subtype labels. **d**, Example compartments shown as scOnion joint, RNA-only, and ATAC-only UMAPs. Low-concordance compartments such as CD4 naive show separations clearer in one modality, whereas high-concordance compartments such as CD8 memory show separations supported by both (Fig. 3f). All eight compartments are shown; Fig. 3f shows two of them, with the joint embedding recomputed (Methods). Statistical tests and sample sizes are reported in Methods.

**Supplementary Figure 4:**
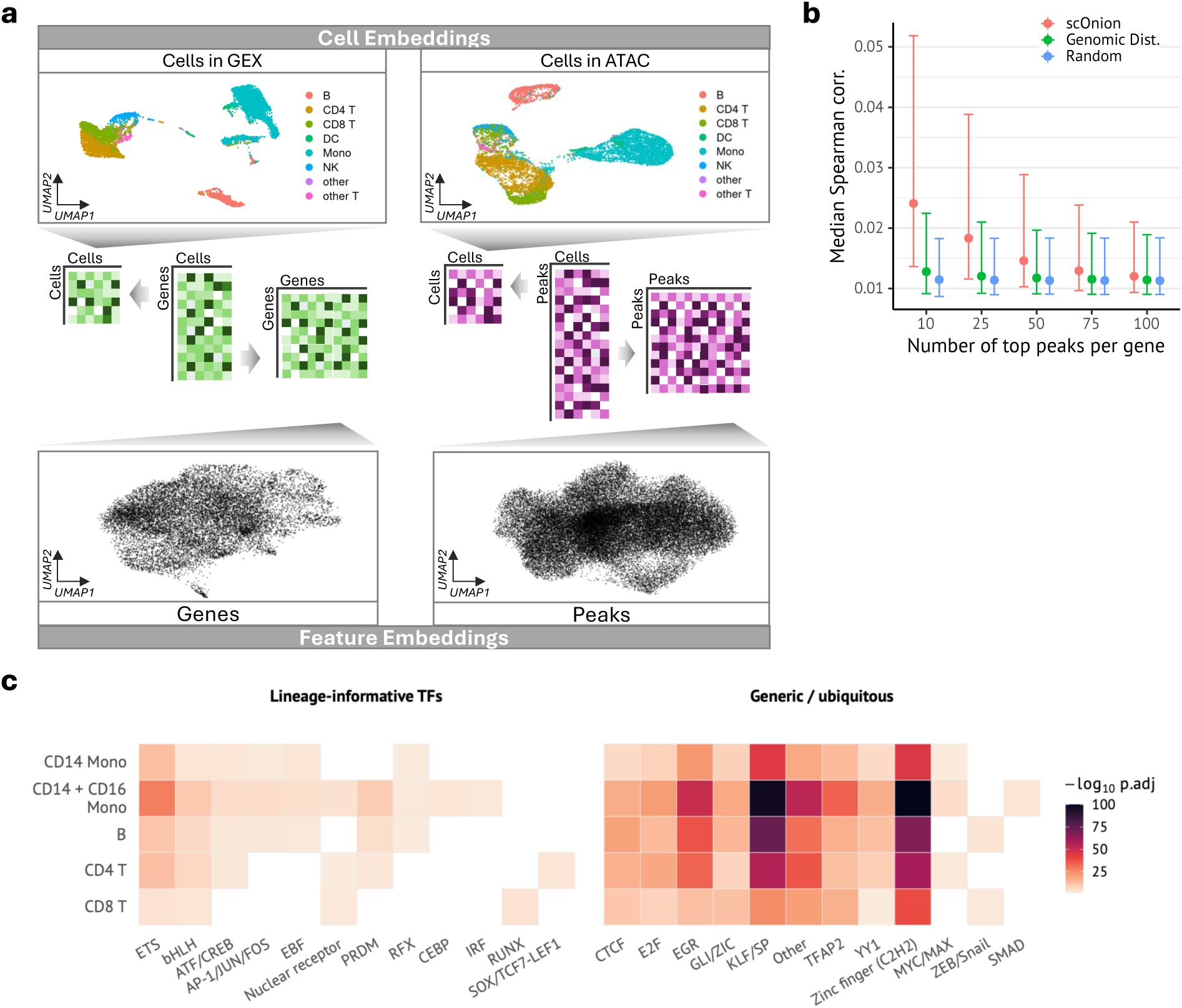
Joint feature embedding, linkage co-activity, and motif enrichment. **a**, Cell and feature embeddings. Top, UMAPs of the GEX and ATAC cell embeddings, which resolve discrete cell populations. Bottom, UMAPs of the gene and peak feature embeddings, obtained by projecting each modality’s features into the same shared coordinate system (Methods). The feature maps show no comparable clustering, so cross-modal feature relationships are read from coordinates. **b**, Co-activity of a gene with its top-ranked peaks, the mean absolute Spearman correlation across cells between the gene’s expression and the accessibility of each of its top *k* peaks, as a function of *k* (Methods). Points are the median across genes and error bars the interquartile range. scOnion-ranked peaks show higher co-activity than distance-ranked or random peaks, with the gap largest for the top few peaks. **c**, Motif enrichment of scOnion top-linked peaks per cell type, grouped into lineage-informative and generic or ubiquitous transcription-factor families (− log_10_ adjusted *P*). Lineage-informative families show cell-type-restricted enrichment, whereas generic families are broadly enriched.

**Supplementary Figure 5:**
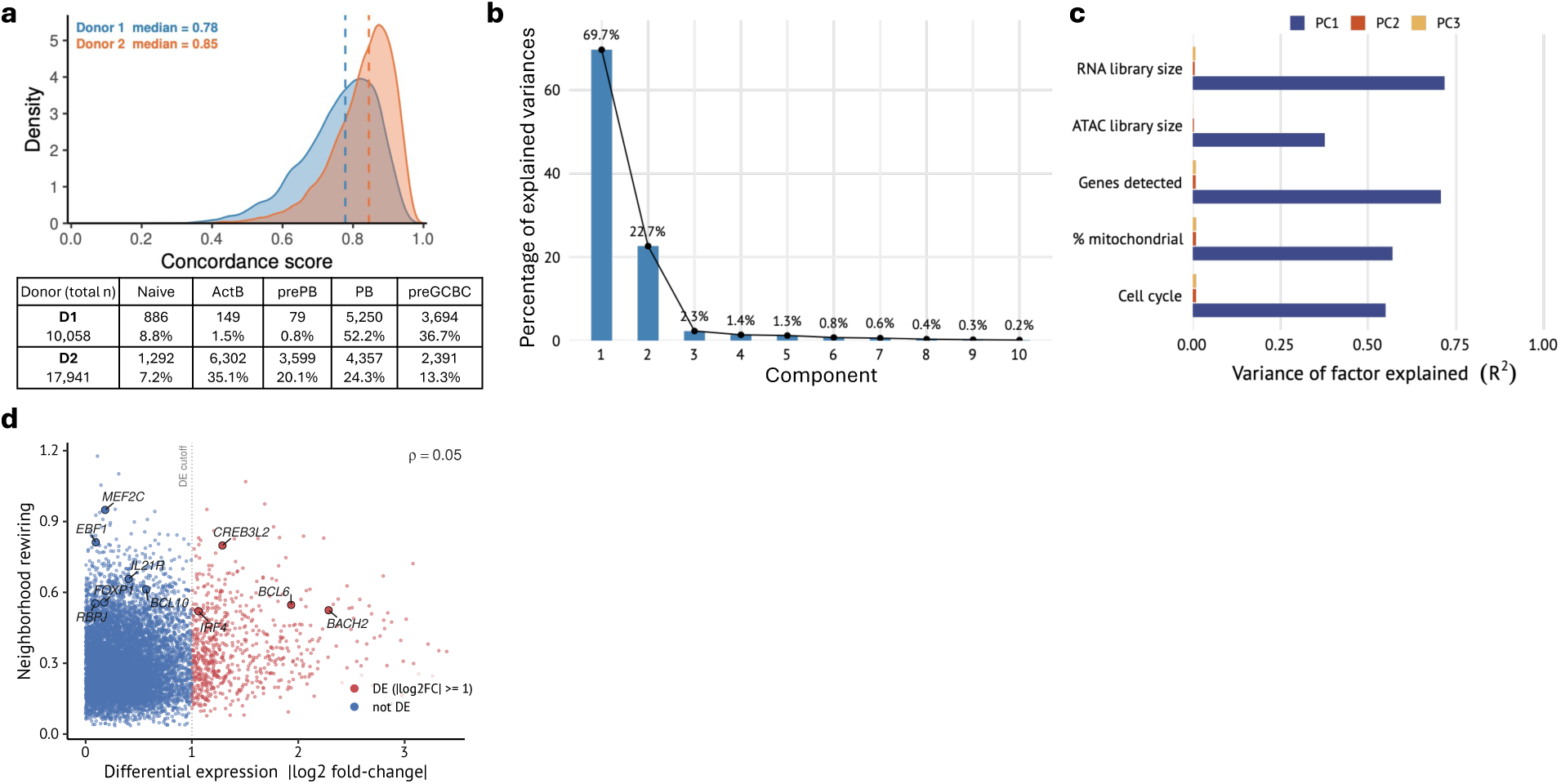
B-cell dataset composition, joint-embedding decomposition, and independence of rewiring from differential expression. **a**, Concordance score distributions for the two IGVF donors (median 0.78 and 0.85), with a table of per-state cell counts and proportions for each donor. The donors differ in cell-state composition, which may contribute to the difference in their overall concordance distributions. The neighborhood-rewiring analysis uses only the PB and preGCBC cells of each donor, which are abundant in both (table). **b**, Scree plot of the scOnion joint cell embedding for the B-cell system, giving the percentage of variance explained by each of the ten leading components. The first component dominates (69.7%) and the second carries 22.7%. Each share is the component’s squared singular value over the sum across the 20 retained components. **c**, Variance of each of the top three scOnion components explained (*R*^2^) by per-cell covariates (RNA and ATAC library size, genes detected, percent mitochondrial reads and cell-cycle score) in donor 2 (17,941 cells). The first component is explained almost entirely by library size and detection, a technical depth axis, whereas the fate component shown in Fig. 5c is largely independent of these covariates. **d**, Neighborhood rewiring score, averaged over the two donors, versus differential ex-pression (|log2FC|, PB versus preGCBC) for all 8,831 tested genes (two-sided Spearman *ρ* = 0.05). Genes above the DE threshold (|log2FC| ≥ 1) are colored, and the eight genes of Fig. 5e together with *EBF1* and *MEF2C* are labeled. High-rewiring non-DE genes such as *EBF1* and *MEF2C* are missed by a differential-expression analysis alone. Statistical tests and sample sizes are reported in Methods.

**Supplementary Figure 6:**
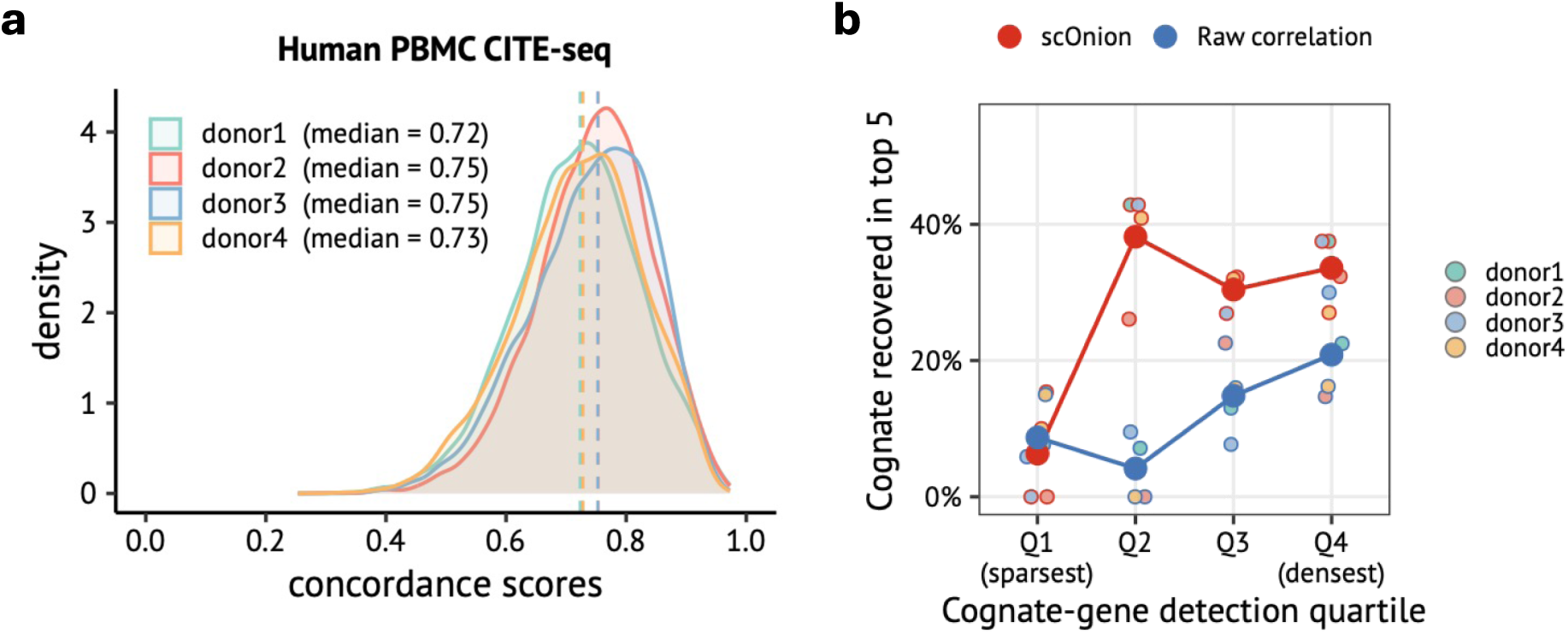
Concordance distributions and sparsity-resolved cognate recovery in CITE-seq. **a**, Concordance score distributions for a human PBMC CITE-seq dataset across four donors (median 0.72 to 0.75). **b**, Cognate-gene recovery in the top 5 as a function of cognate-transcript detection, with proteins binned into quartiles from sparsest to densest cognate-transcript detection, for scOnion and raw Pearson correlation across four donors, restricted to well-marking proteins, those with mean antibody abundance above the per-donor median (104 per donor, 13 to 40 per quartile). In the sparsest quartile both methods recover fewer than 10%. Above it, scOnion recovers the cognate at 30% to 38% in every quartile (mean over donors), whereas correlation im-proves only as detection increases (4% to 21%). Points are the donor mean, and the four per-donor rates are shown.

